# mCranberry: A rationally engineered monomeric red fluorescent protein with 85% quantum yield

**DOI:** 10.64898/2026.10.01.755472

**Authors:** Shivam Chaubey, Srijit Mukherjee, Anita Adams, Alice R. Walker, Melonnie L. M. Furgason, Heman Kumar Ramiya Ramesh Babu, Irimpan I. Mathews, Premashis Manna

## Abstract

Fluorescent proteins (FPs) are the workhorse of modern-day cellular imaging and biosensing. However, rationally predicting and engineering the photophysical properties of FPs remain challenging, as these are governed by complex three-dimensional microenvironments and post-translational modifications. Here, we present a novel surface-mutation-guided engineering strategy combined with iterative rounds of rational mutagenesis to improve key photophysical properties of red fluorescent proteins (RFPs). This approach generated two bright, monomeric RFP variants from the mFruits lineage, mCranberry and mCranberry2, with quantum yields of 85% and 80%, respectively. Strategic incorporation of charged amino acid residues on the protein surface enhanced fluorescence quantum yield without compromising photostability. Comprehensive photophysical characterization and live-cell microscopy demonstrate enhanced performance of these variants and establish their utility as robust probes for fluorescence imaging. However, the exact mechanism of such improvements is subject to further investigation. Collectively, this work introduces a strategy for optimizing fluorescence properties and provides a framework that may be broadly applicable to other pigment-protein systems such as genetically encoded metal-ion indicators and photosynthetic light-harvesting complexes.

## Introduction

Fluorescent proteins (FPs) have transformed biological imaging and biosensing by enabling genetically encoded visualization of cellular and molecular processes.^1^ In the last decade, significant progress has been made in optimizing their photophysical properties such as brightness,^2–5^ photostability,^6,7^ pH resistance,^8^ oligomerization tendencies,^9^ etc. Brightness and photostability of an FP dictate the quality and duration of fluorescence imaging. Improving both properties simultaneously has been proven challenging as they are often found to be inversely correlated.^10^ Although recent engineering efforts have produced green and yellow fluorescent proteins with substantial improvements in both properties, comparable advances in RFPs have remained limited.^11,12^ This may be attributed to the presence of significant light-induced non-fluorescent or dark states in RFPs, particularly in the so-called mFruit series.^13,14^ Therefore, novel strategies are required to overcome this limitation to engineer bright and photostable RFPs.

Recent advances in artificial intelligence (AI) yielded novel pigment-binding proteins or FPs with reduced sizes.^15,16^ However, AI-driven enhancement of quantum yield and brightness in FPs remains elusive.^17^ For instance, a protein language model-guided discovery of esmGFP, which is evolutionarily distant from previously characterized green fluorescent proteins, shows lower brightness and slower chromophore maturation than the wild-type.^18^ Accurately predicting photophysical properties in FPs is challenging, as they are governed not only by accurate folding but also by post-translational modifications of the chromophore-forming tripeptides and complex interactions with their neighboring residues.

Here, we report a rational engineering strategy for improving the performance of RFPs. We reasoned that charged amino acid residues positioned along the plane of the chromophore transition dipole moment (TDM) of the FPs could modulate the radiative and non-radiative transition rate and thereby influence fluorescence quantum yield.^19^ Following this approach, we demonstrate that targeted substitution of surface residues distal from the chromophore, in some cases through a single amino acid substitution, can substantially increase fluorescence quantum yield while preserving photostability. This strategy yielded a series of improved RFP variants, including mCranberry and mCranberry2, with markedly enhanced quantum yields. We performed extensive photophysical measurements, structural analysis, and live-cell imaging to validate their robustness as imaging probes. Molecular dynamics (MD) simulations of the variants provide mechanistic insights into the role of surface mutations in tuning fluorescent protein properties. However, whether the observed improvements in the mCranberry series are indeed due to hypothesized modulation of TDM amplitude and/or orientation is subject to further investigation.

## Results & Discussion

### Rational engineering of mCranberry

The engineering strategy leading to mCranberry is summarized in Fig. 1. As the starting scaffold, we selected the R125K variant of mCherry-XL, which we call mCherry-XL R125K. R125K mutation appeared during a random mutagenesis screening, which improved the cellular brightness in yeast of its precursor, mCherry-XL. mCherry-XL is an engineered mCherry variant with a fluorescence quantum yield of 70%.^20^ It was previously developed by optimizing the excited-state lifetime of mCherry through directed evolution, resulting in four chromophore-proximal substitutions (W143S, I161V, Q163Y, and I197R) that increased the quantum yield from 22% to 70%. Subsequent site-directed mutagenesis of residues surrounding the chromophore produced only marginal improvements, suggesting that the local chromophore environment had approached an optimal balance of chromophore maturation, folding, and fluorescence quantum yield. We therefore reasoned that further enhancement might require targeting residues distal to the chromophore. Our engineering strategy was motivated by the hypothesis that surface residues can modulate the electrostatic environment within the β-barrel across the chromophore. As the TDM influences both the extinction coefficient and the radiative decay rate,^21^ we reasoned that strategically positioned charged residues could alter its amplitude and/or orientation and thereby enhance fluorescence brightness. Importantly, mutations on the exterior of the β-barrel were expected to perturb the electrostatic environment while minimizing direct effects on chromophore maturation, folding, or other structural determinants of fluorescence.^5^ We further hypothesized that residues located approximately within the plane of the TDM would exert the greatest influence on its magnitude.

**Fig. 1.**
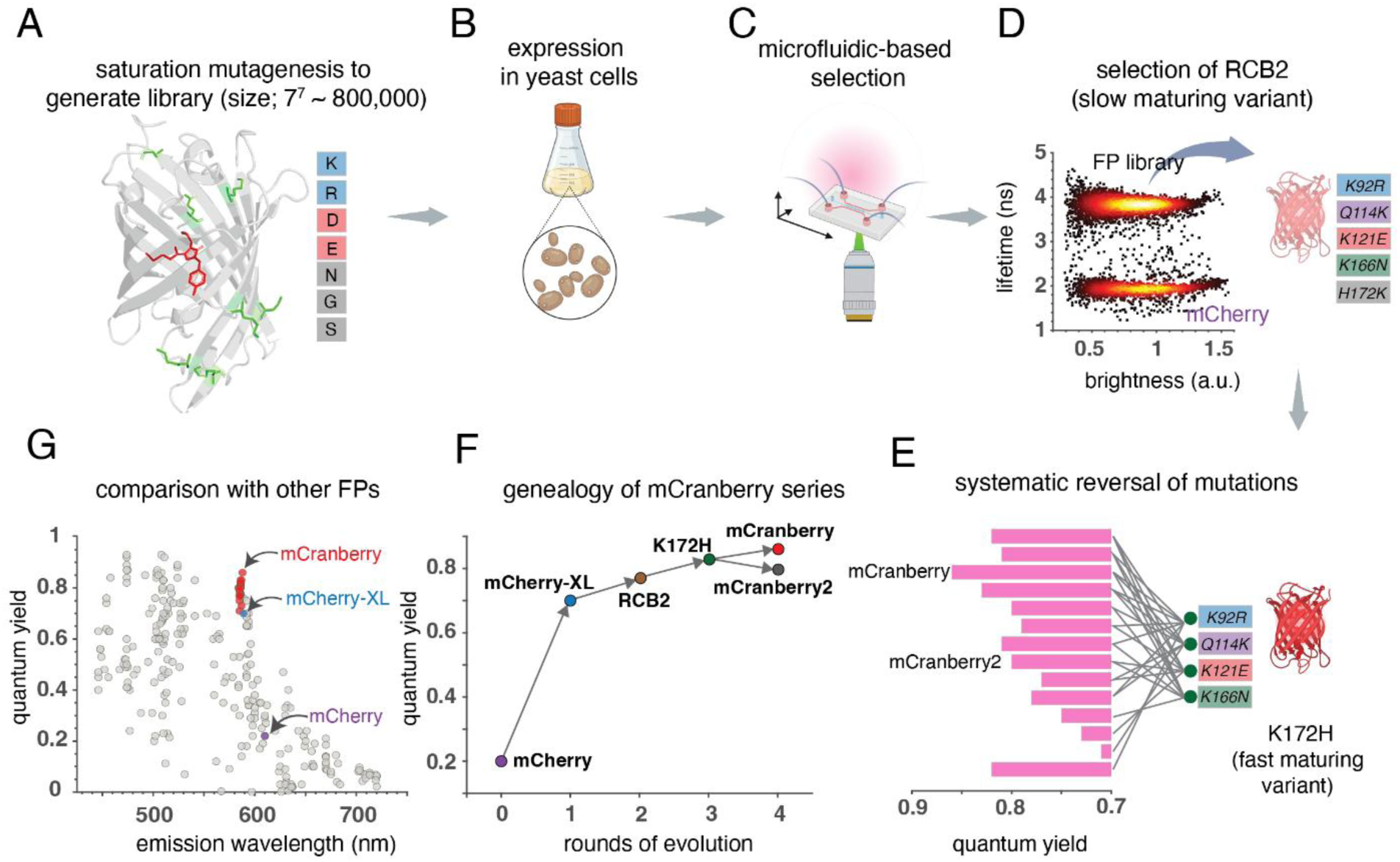
Directed evolution of mCranberry and mCranberry2. **A,** Amino acid residues targeted to create a protein library are shown in the mCherry crystal structure (PDB: 2H5Q).^23^ The target residues are shown in green, the protein backbone in grey, and the chromophore in red. Seven positions (Table 1) were saturated with two negatively charged (D, E), two positively charged (K, R), and asparagine, serine, and glycine. The total library size was 7^7^ ∼ 800,000. **B, C,** The resulting library was expressed in yeast cells and sorted on a custom-made high-throughput microfluidic platform based on fluorescence lifetime and intensity of the individual cells. **D,** Scatter plot displaying the lifetime and brightness of mCherry and FP library expressed in yeast. RCB2 mutant was selected from this FP library. The mutations of RCB2 compared to mCherry-XL R125K are shown in colored boxes. **E,** Systematic reversal of mutations of RCB2 to mCherry like residues yielded a fast-maturing variant of RCB2 (shown as K172H), where the H172K mutation was reverted. A series of RFP mutants was generated by a combinatorial library where all the mutations of K172H were reverted to mCherry-like residues. The quantum yields of the mutants are shown in pink bar plots. mCranberry is characterized by the highest quantum yield of this series (85%), whereas mCranberry2 shows faster chromophore maturation**. F,** The genealogy of mCranberry and mCranberry2 mutants**. G,** Quantum yield and emission wavelength of the fluorescent proteins from the FPbase database (https://www.fpbase.org/). All the mutants selected are constitutively fluorescent and monomeric in nature. Red filled circles represent mCranberry variants, blue circles indicate mCherry-XL and purple circle denote mCherry.

To identify candidate positions, we examined mutations that emerged during the evolution of several bright RFPs, including mRuby3,^5^ mScarlet,^2^ and mCherry-d.^13^ This analysis revealed recurring substitutions at solvent-exposed positions distal to the chromophore. For instance, the N98K and K166R substitutions appeared during the evolution of brighter C4-derived variants, including mCherry-d, with K166R conserved across all C4PB variants.^13,22^ Similarly, the K166R mutation was introduced during the evolution of mScarlet,^2^ whereas the evolution of mRuby3 introduced both the C114E substitution near the imidazolinone-end of the chromophore and the S171H substitution near the phenolate-end.^5^ The repeated emergence of charged substitutions at these positions suggested that distal electrostatic interactions may contribute to improved brightness.

As the TDM reflects the redistribution of electric charge between the ground and excited states, we further reasoned that residues flanking both ends of the chromophore could influence its amplitude and/or orientation. We therefore designed a saturation mutagenesis library targeting solvent-exposed residues surrounding both the phenolate and imidazolinone-ends of the chromophore (Fig. 1A). The selected positions and the rationale for their inclusion are summarized in Table 1.

**Table 1.** Targets for the FP library and the rationales for including them. Numbering is based on the mCherry crystal structure (PDB: 2H5Q). ^23^

| Residues | Position | Rationale |
| --- | --- | --- |
| 92 | Imidazolinone-end | Appeared in mApple (K92R), improves brightness |
| 98 | Phenolate-end | Appeared in mCherry-d, improves brightness |
| 114 | Imidazolinone-end | Appeared in mRuby3 (C114E) |
| 121 | Imidazolinone-end | Appeared in mRuby3 (H121N) |
| 162 | Phenolate-end | Appeared in mRuby3 |
| 166 | Phenolate-end | Appeared in mScarlet, mCherry-d |
| 172 | Phenolate-end | Appeared in mScarlet and mRuby3 |

To test our hypothesis, we constructed saturation mutagenesis libraries enriched for charged amino acid substitutions. Although our initial goal was to restrict the libraries to the charged residues lysine (K), arginine (R), glutamate (E), and aspartate (D), analysis of codon degeneracy showed that this could not be achieved without introducing a small number of additional amino acids. As histidine is partially protonated near physiological pH (pKa ≈ 6), we excluded it from the library design. We therefore employed the degenerate RRN codon, which encodes the four desired charged residues together with three uncharged residues (serine, asparagine, and glycine) while maintaining a relatively uniform amino acid distribution. For the combinatorial library, seven solvent-exposed positions were simultaneously diversified using the RRN codon, yielding seven possible amino acids (K, R, D, E, S, N, and G) at each site. The theoretical library size was therefore 7^7^ or ∼8.2 × 10^5^ variants. The resulting FP library was constructed and expressed in *Saccharomyces cerevisiae* for high-throughput screening as described elsewhere (Fig. 1B).^24^

Given the large size of the combinatorial library, we first enriched for fluorescent variants by fluorescence-activated cell sorting (FACS), collecting the brightest 0.3% of the population.^25^ The enriched FP library was subsequently screened using a custom-built microfluidic lifetime-sorting platform (Fig. 1C).^24^ As Figure 1D displays, fluorescence lifetime analysis revealed two major populations: one centered at approximately 2 ns, corresponding to mCherry-like variants, and a second population centered near 4 ns, consistent with mCherry-XL-like variants. Variants from the ∼ 4 ns lifetime population were further enriched, plated on galactose-containing agar, and individually characterized based on fluorescence brightness and lifetime. Among the selected variants, RCB2 exhibited the greatest improvement in fluorescence brightness in yeast, displaying approximately sixfold higher fluorescence than mCherry during FACS analysis. Relative to mCherry-XL R125K, RCB2 was characterized with five amino acid substitutions: K92R, Q114K, K121E, K166N, and H172K. Surprisingly, despite its favorable photophysical properties, RCB2 showed only modest improvements in fluorescence brightness in HeLa cells. We hypothesized that this discrepancy in brightness between yeast and HeLa cells resulted from slower chromophore maturation, consistent with the delayed appearance of its characteristic pink coloration compared with mCherry during protein expression.

To identify mutations responsible for the slow maturation, we individually reverted each RCB2 substitution to the corresponding residue in mCherry-XL R125K. Among these variants, reversion of lysine at 172 to histidine (i.e. K172H) substantially accelerated chromophore maturation while preserving the enhanced fluorescence properties (Fig. 1E). We next examined the remaining substitutions by systematically reverting individual and combined mutations, generating a total of 14 variants, collectively referred to as the **mCranberry** series.

Photophysical characterization of the mCranberry series identified two particularly promising variants (Fig. 1E). Variant G12 exhibited a fluorescence quantum yield of 85%, the highest reported for a monomeric, constitutively fluorescent RFP to date. Variant G7 displayed substantially faster chromophore maturation, producing visible fluorescence with kinetics comparable to mCherry during bacterial expression. We designated G12 and G7 as **mCranberry** and **mCranberry2**, respectively. Their evolutionary relationship is summarized in Fig. 1F, whereas their positions relative to previously reported monomeric, constitutively fluorescent RFPs are shown in Fig. 1G.

### Photophysical and in-vivo characterization of the mCranberry variants

The photophysical properties of the mCranberry variants are summarized in Table 2. Figures 2A and 2B display absorption, excitation and emission spectra of mCranberry and mCranberry2, respectively. The introduction of surface mutations in mCherry-XL R125K (K121E in mCranberry; K92R and Q114K in mCranberry2) improved the quantum yield from 72% to 85% and 80%, respectively, without altering absorption/emission peaks, spectral broadness, and Stokes shifts (Extended Data Fig. 1). Both the variants were characterized with an excited state lifetime of 3.8 ns and fluorescence decays followed single-exponential kinetics (Figure 2C, Table 2, Extended Data Fig. 5). From the quantum yield and lifetime values, radiative and non-radiative rate constants of the RFPs are quantified (Fig. 2D, Extended Data Fig. 6). We observe that radiative rate constant of mCranberry is higher compared to other RFP variants which might contribute to its higher quantum yield. pKa values of mCranberry and mCranberry2 were measured as 4.9 and 5.1, respectively, making them suitable probes even in acidic environments (Fig. 2E). Finally, size-exclusion chromatography of the RFPs shows a similar elution profile as the parent mCherry, indicating that the surface mutations introduced in mCranberry variants do not alter their monomericity (Fig. 2F).

**Fig. 2.**
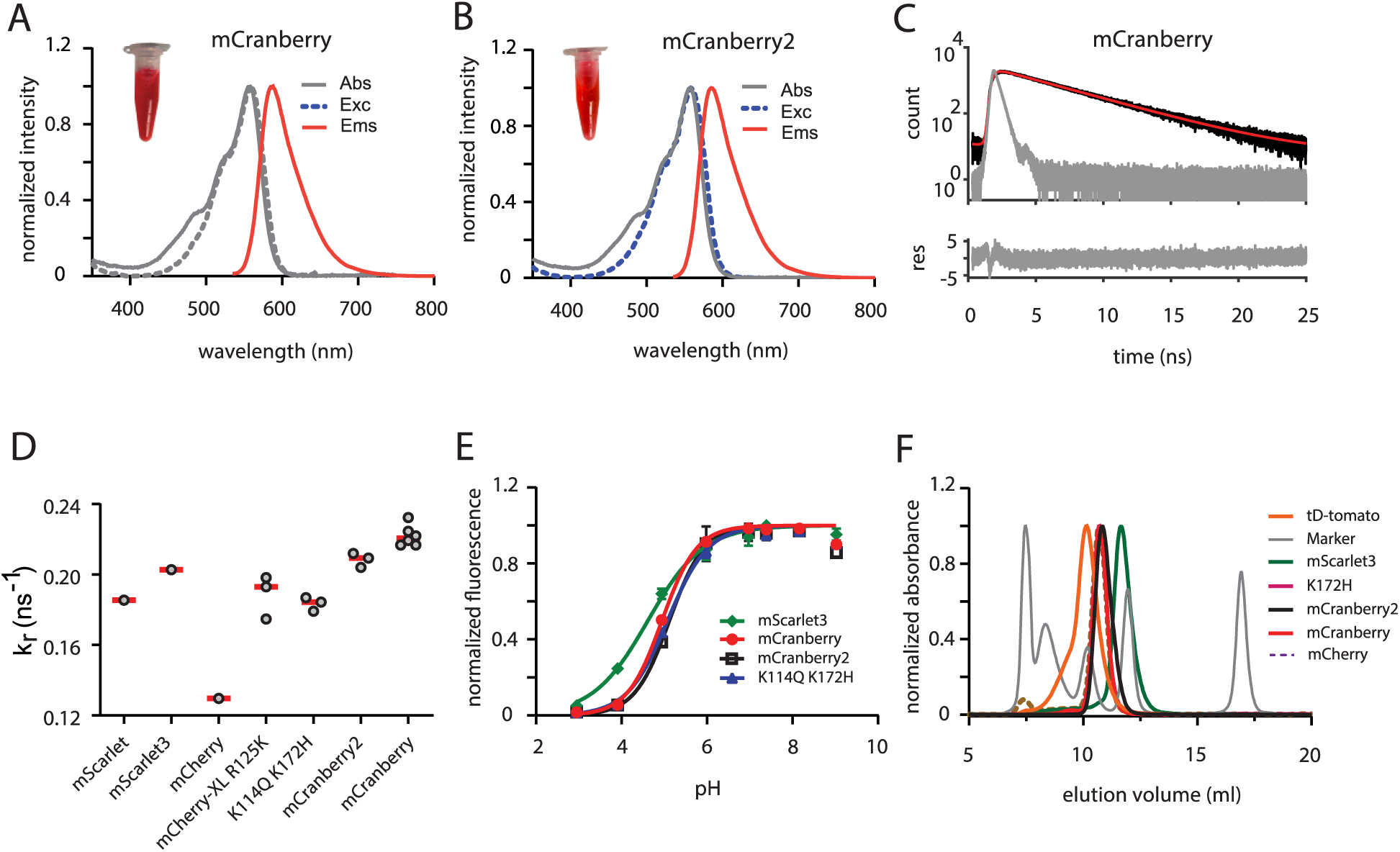
*In Vitro* characterization of mCranberry mutants. **A**, **B**, Normalized absorbance (Abs), excitation (Exc) and emission (Em) spectra of mCranberry (**A**) and mCranberry2 (**B**)**. C,** Excited state lifetime decays (black line) of mCranberry and the single-exponential fit (red line). The instrument response function is shown in grey and the residuals of the fit are shown at the bottom. **D,** Radiative rate constants of the FPs obtained from the experimental quantum yields and fluorescence lifetimes. **E,** pH-dependent fluorescence intensities of the RFPs with error bars indicating the standard deviations across three biological replicates. The solid lines represent the sigmoidal non-linear regression fits used to determine the apparent pKa for each variant. **F**, Size-exclusion chromatography of the purified proteins was monitored with absorbance at 280 nm. The marker peaks (Bio-Rad gel filtration standard, 1511901), from left to right, corresponded to thyroglobulin (670,000 Da), γ-globulin (158,000 Da), ovalbumin (44,000 Da), myoglobin (17,000 Da), and vitamin B12 (1,350 Da). mCranberry eluted at the same volume as mCherry, indicating that the introduced surface mutations did not alter its oligomeric state and that the protein remained monomeric.

**Table 2.**
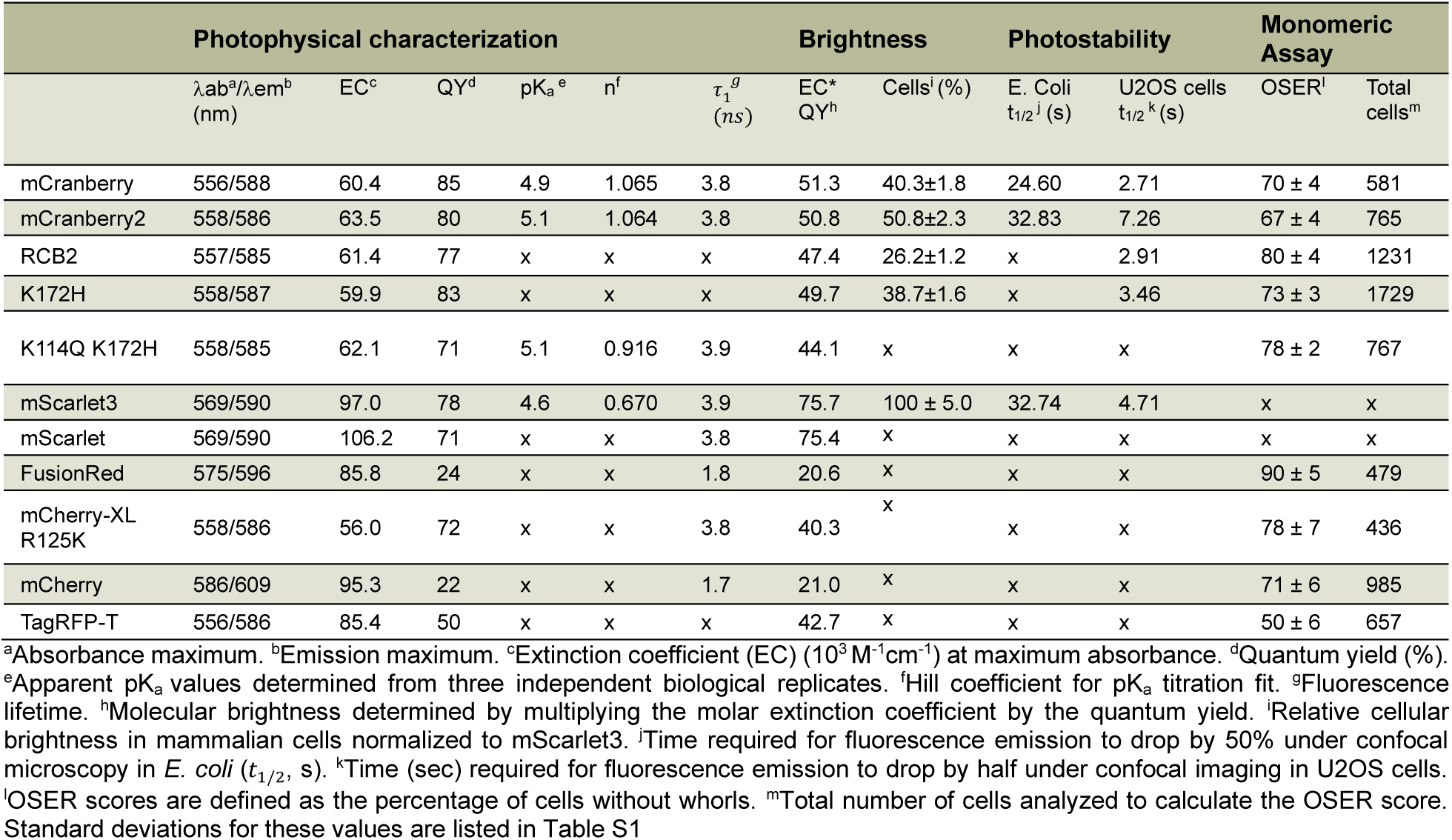
Characterizations of mCranberry and other reference RFPs.

|  | Photophysical characterization |  |  |  |  |  | Brightness |  | Photostability |  | Monomeric Assay |  |
| --- | --- | --- | --- | --- | --- | --- | --- | --- | --- | --- | --- | --- |
| | $\lambda_{ab}^a/\lambda_{em}^b$<br>(nm) | EC<sup>c</sup> | QY<sup>d</sup> | pK<sub>a</sub><sup>e</sup> | n<sup>f</sup> | $\tau_1^g$<br>(ns) | EC*<br>QY<sup>h</sup> | Cells<sup>i</sup> (%) | E. Coli<br>$t_{1/2}^j$ (s) | U2OS cells<br>$t_{1/2}^k$ (s) | OSER<sup>l</sup> | Total<br>cells<sup>m</sup> |
| mCranberry | 556/588 | 60.4 | 85 | 4.9 | 1.065 | 3.8 | 51.3 | 40.3±1.8 | 24.60 | 2.71 | 70 ± 4 | 581 |
| mCranberry2 | 558/586 | 63.5 | 80 | 5.1 | 1.064 | 3.8 | 50.8 | 50.8±2.3 | 32.83 | 7.26 | 67 ± 4 | 765 |
| RCB2 | 557/585 | 61.4 | 77 | x | x | x | 47.4 | 26.2±1.2 | x | 2.91 | 80 ± 4 | 1231 |
| K172H | 558/587 | 59.9 | 83 | x | x | x | 49.7 | 38.7±1.6 | x | 3.46 | 73 ± 3 | 1729 |
| K114Q K172H | 558/585 | 62.1 | 71 | 5.1 | 0.916 | 3.9 | 44.1 | x | x | x | 78 ± 2 | 767 |
| mScarlet3 | 569/590 | 97.0 | 78 | 4.6 | 0.670 | 3.9 | 75.7 | 100 ± 5.0 | 32.74 | 4.71 | x | x |
| mScarlet | 569/590 | 106.2 | 71 | x | x | 3.8 | 75.4 | x | x | x | x | x |
| FusionRed | 575/596 | 85.8 | 24 | x | x | 1.8 | 20.6 | x | x | x | 90 ± 5 | 479 |
| mCherry-XL<br>R125K | 558/586 | 56.0 | 72 | x | x | 3.8 | 40.3 | x | x | x | 78 ± 7 | 436 |
| mCherry | 586/609 | 95.3 | 22 | x | x | 1.7 | 21.0 | x | x | x | 71 ± 6 | 985 |
| TagRFP-T | 556/586 | 85.4 | 50 | x | x | x | 42.7 | x | x | x | 50 ± 6 | 657 |

<sup>a</sup>Absorbance maximum. <sup>b</sup>Emission maximum. <sup>c</sup>Extinction coefficient (EC) ( $10^3 \text{ M}^{-1} \text{ cm}^{-1}$ ) at maximum absorbance. <sup>d</sup>Quantum yield (%). <sup>e</sup>Apparent pK<sub>a</sub> values determined from three independent biological replicates. <sup>f</sup>Hill coefficient for pK<sub>a</sub> titration fit. <sup>g</sup>Fluorescence lifetime. <sup>h</sup>Molecular brightness determined by multiplying the molar extinction coefficient by the quantum yield. <sup>i</sup>Relative cellular brightness in mammalian cells normalized to mScarlet3. <sup>j</sup>Time required for fluorescence emission to drop by 50% under confocal microscopy in *E. coli* ( $t_{1/2}$ , s). <sup>k</sup>Time (sec) required for fluorescence emission to drop by half under confocal imaging in U2OS cells. <sup>l</sup>OSER scores are defined as the percentage of cells without whorls. <sup>m</sup>Total number of cells analyzed to calculate the OSER score. Standard deviations for these values are listed in Table S1

Consistent with our analytical size-exclusion chromatography data, the organized smooth endoplasmic reticulum (OSER) assay^26,27^ confirm that the mCranberry variants exhibit a favorable monomeric profile (Figure 3A). The RFP variants were expressed as CytERM fusion constructs in U2OS cells, followed by live-cell imaging and quantitative image analysis (SI Sec 2, Figure S2A & S2B).^24^ mCranberry and mCranberry2 exhibited similar OSER scores as their precursor mCherry, indicating that the engineered surface mutations do not compromise the monomeric behavior of the FPs. Figure 3B displays representative OSER images of the mCranberry variants along with FusionRed (positive control) and TagRFP-T (negative control).

**Fig. 3.**
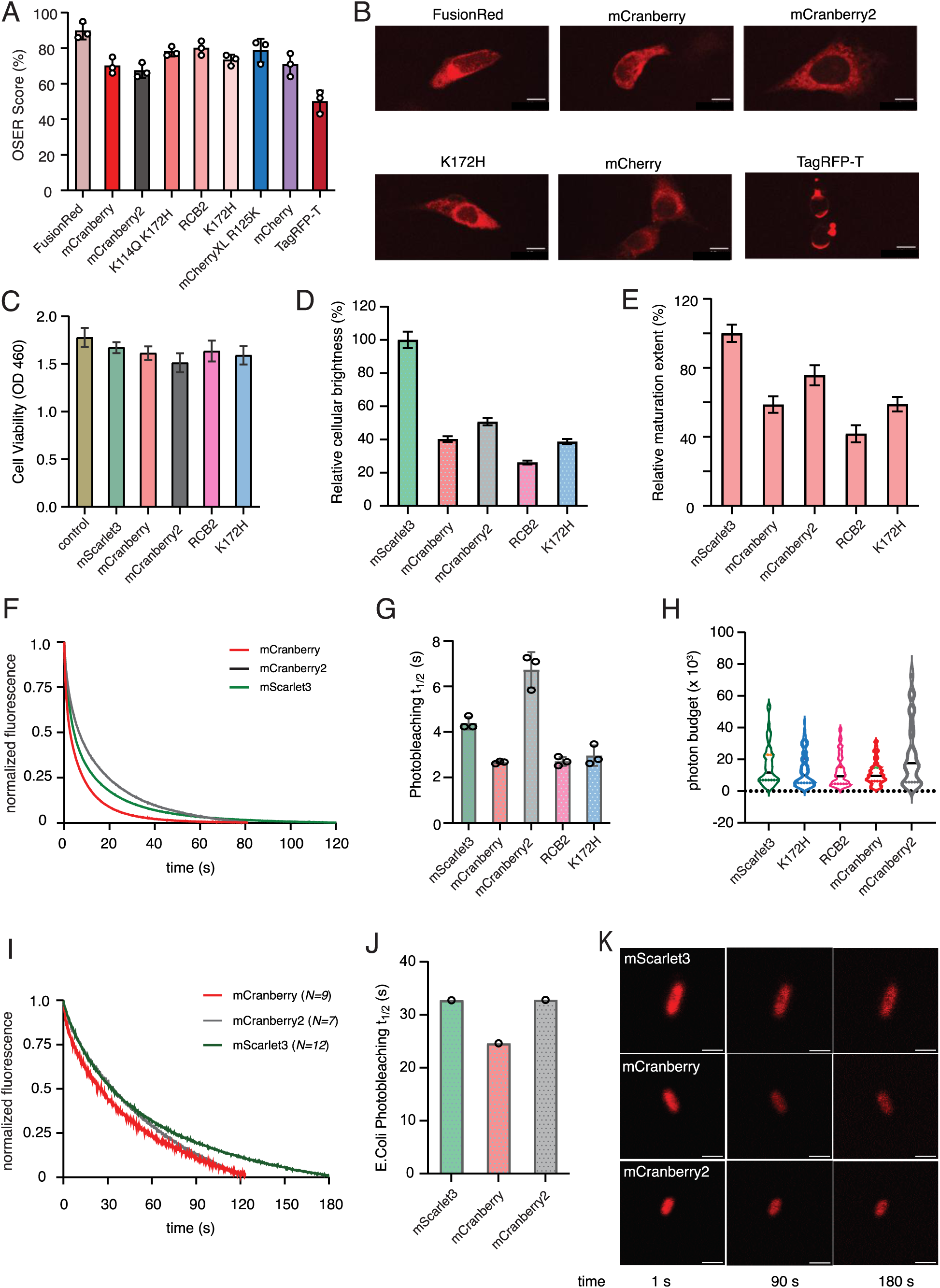
Cellular performance of mCranberry and mCranberry2. **A,** OSER scores for proteins assessed 24 hours post-transfection with CytERM-RFP in U2OS cells. Scores represent the percentage of cells lacking whorls, with error bars reflecting standard deviations from three independent biological replicates. **B,** Representative fluorescence images of CytERM-RFP construct in transfected U2OS cells. Scale bar, 10 μm. Red fluorescence intensity is displayed with uniform brightness and contrast adjustments across all panels. CytERM-mCranberry displays proper reticular morphology, whereas CytERM-TagRFP-T forms small whorl-like structures due to its in vivo oligomerization tendency. **C,** Cellular cytotoxicity following 48-hour transfection of U2OS cells with pDRESS_mTurquoise2_spatial-linker-P2A_RFP was assessed by measuring the reduction of WST-8 to water-soluble formazan, with absorbance read at 460 nm. Results are reported as mean absorbance at 460 nm ± s.d. across six independent transfections. **D,** Relative cellular brightness (means ± s.d) of RFPs in U2OS cells 48 hours post-transfection with pDRESS_mTurquoise2_spatial-linker-P2A_RFP. To determine the relative brightness within mammalian cells, the raw red-to-cyan ratios were normalized by accounting for the specific spectral throughput and detection efficiency of the imaging system. The brightness here is shown relative to mScarlet3. **E,** Relative maturation extent of RFPs in U2OS cells determined by dividing cellular brightness by molecular brightness (normalized to mScarlet3). Data are shown as mean ± s.d. **F**, Confocal photobleaching decay in U2OS cells. Normalized fluorescence intensity is plotted as a function of continuous illumination time (seconds). Lines represent the mean decay curve obtained from three independent biological replicates (20-25 cells analyzed per replicate, Extended Data Fig. 6). The time axis was corrected as described in the methods to account for RFP dependent differences in extinction coefficient, quantum yield, and excitation efficiency. **G,** U2OS photobleaching half-times (*t*_1/2_) indicate the time required for fluorescence intensity to decay to 50% of its initial intensity. **H,** Single-cell photon budget distributions using violin scatter plots to evaluate the total number of photons emitted by the RFPs before photobleaching. **I**, Confocal photobleaching decay in fixed *E. coli*, Normalized remaining fluorescence intensity (% of initial signal) as a function of continuous illumination time (seconds). *N* represents the number of cells analyzed for each mutant. **J,** E.Coli Photobleaching half-times (*t*_1/2_), indicate the time required for fluorescence emission to decay to 50% of its initial intensity. **K,** Representative confocal microscopy images of RFP-expressing *E. coli* at three time points during photobleaching. Scale bar: 2 μm. Red fluorescence intensity is displayed with uniform brightness and contrast adjustments across all panels.

While FPs serve as standard whole-cell labels and fusion tags, high-level expression often triggers cytotoxic stress. This issue is especially pronounced among RFPs, which exhibit substantially higher cytotoxicity in whole-cell labels when compared to mEGFP in both bacterial and mammalian expression systems.^28^ Establishing minimally cytotoxic RFP variants is therefore an important parameter in their directed evolution. Cell viability assays performed 48 h post-transfection with pDRESS-mTurquoise2–spatial linker–P2A-RFP (Fig. 3C) and 24 h with CytERM-RFP (Extended Data Fig.10) demonstrated that the mCranberry RFP variants exhibited no significant cytotoxicity relative to control cells.

We next characterized the intracellular brightness of the mCranberry variants by confocal fluorescence microscopy. To account for differences in expression levels between cells, each RFP was co-expressed with the cyan fluorescent protein mTurquoise2,^4^ and the red fluorescence intensity was normalized to the corresponding cyan fluorescence (Extended Data Fig.8).^3^ Although mCranberry and mCranberry2 were characterized with record fluorescence quantum yield, their brightness in U2OS cells were found to be 40-50% of mScarlet3 (Fig. 3D, Table 2). The relative maturation extent of the RFPs, which measures the ratio of cellular brightness to the molecular brightness (extinction coefficient x quantum yield), was found to be lower compared to mScarlet3, indicating slower/incomplete chromophore maturation (Fig. 3E). Lower molecular brightness of mCranberry variants relative to mScarlet3 is due to their lower extinction coefficients compared to mScarlet3 (∼ 60,000 M^-1^cm^-1^ vs. 97,000 M^-1^cm^-1^). It appears that, although surface mutations improved the quantum yield of the variants, but they reduced their extinction coefficients–a mechanism that is not clear and subject to future investigation.

Fig. 3F shows the photobleaching decays of mCranberry, mCranberry2 and mScarlet3 in U2OS cells. We observed that mCranberry2 is 1.7-fold more photostable than mScarlet3 (Fig. 3G and Table 2). This resulted in a higher median photon budget (total number of photons released before photobleaching) of mCranberry2 compared to other variants (Fig. 3H). However, in bacteria, mCranberry2 and mScarlet3 were found to have similar photostability (Fig. 3I and 3J). Fig. 3K displays representative time-lapse confocal images of bacteria expressing RFPs.

In general, it has been observed a negative correlation between photostability and brightness in RFPs.^10^ For instance, the improved photostability in mScarlet-H,^2^ mScarlet3-H,^6^ mScarlet3-S2^29^ have been achieved with compromising their quantum yield and thereby brightness. On the contrary, surface mutations introduced in mCranberry and mCranberry2 improved their quantum yields without negatively affecting their photostability. Moreover, surface mutations in mCranberry2 enhanced its photostability. To the best of our knowledge, this is the first observation of such a phenomenon, which may be beneficial in engineering other FPs or pigment-protein complexes, in general. In order to understand the mechanism behind such effect, we performed structural analysis and molecular dynamics (MD) simulations of the RFPs.

### Structural analysis of the engineered variants

To investigate how distal surface mutations influence fluorescence quantum yield, we determined the crystal structure of mCranberry2 (Fig. 4A, 4B). Diffraction data were collected, and the structure was refined to a resolution of 3.2 Å. The overall architecture of mCranberry2 retained the canonical fluorescent protein fold, consisting of an eleven-stranded β-barrel surrounding the autocatalytically formed chromophore (Fig. 4B). Mapping the engineered substitutions onto the structure revealed that the mutations responsible for the initial increase in quantum yield from mCherry to mCherry-XL (W143S, I161V, Q163Y, and I197R) are clustered around the chromophore, consistent with previous engineering efforts that primarily targeted the chromophore environment (Fig. 4B). In contrast, the additional substitutions identified in this study, including K92R and Q114K, are located on the protein surface and are spatially separated (12 and 19 Å, respectively) from the chromophore.

**Fig 4.**
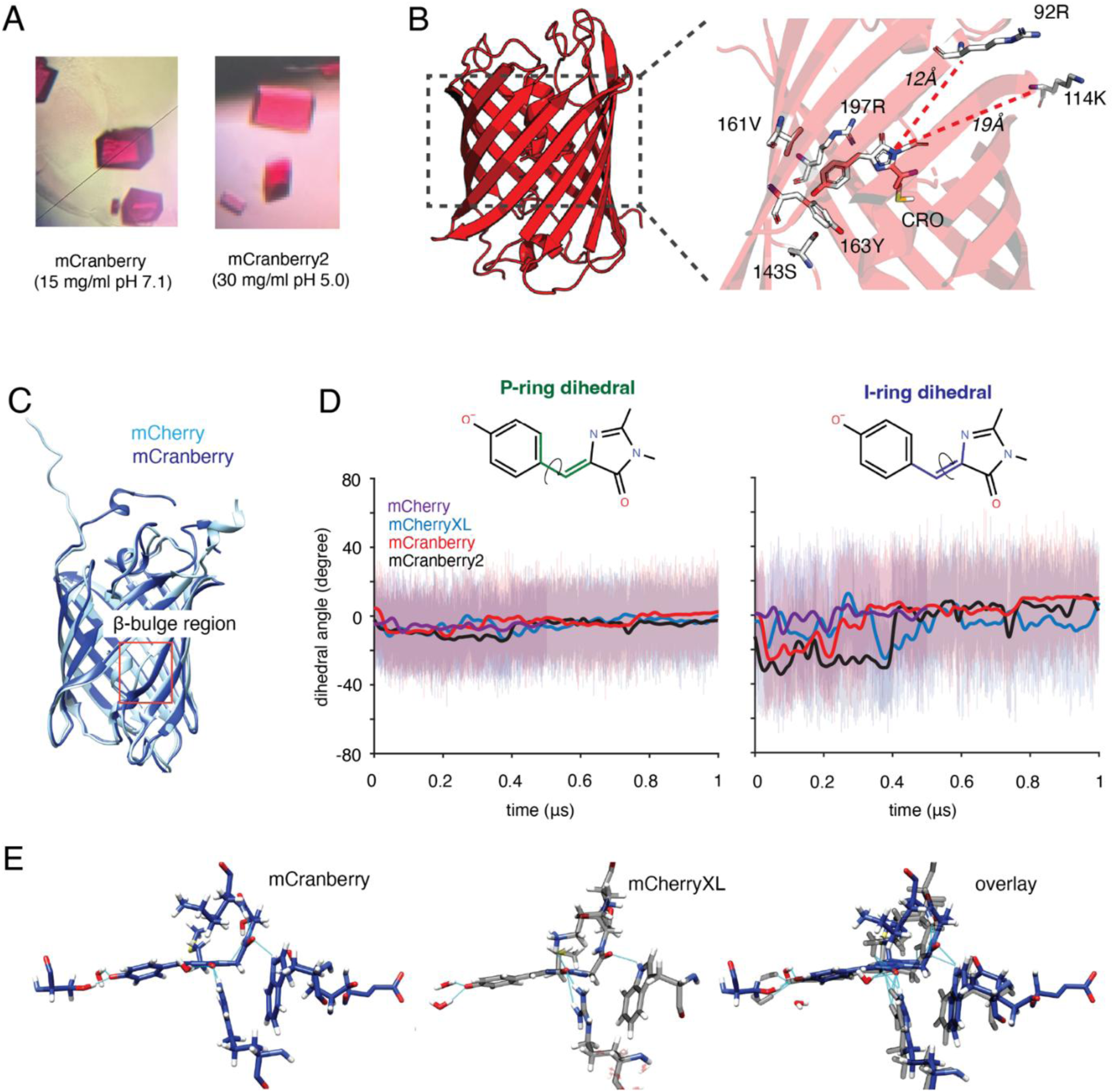
Crystal structure and MD simulation of mCranberry mutants. **A,** Representative stereomicroscopic views of protein crystals obtained for mCranberry and mCranberry2. mCranberry crystals were obtained at 15 mg ml⁻¹ protein in 5% (w/v) choline acetate and 30% PEG 400 at pH 7.1, whereas mCranberry-2 crystallized at 30 mg ml⁻¹ protein in 22% (w/v) PEG 3350 and 0.1 M sodium citrate tribasic dihydrate at pH 5.0. **B,** Crystal structure of mCranberry2 determined at a resolution of 3.2 Å, highlighting the chromophore and surface-mutated amino acid residues. **C**, Overlay of representative molecular dynamics structures of wild type mCherry shown in cyan and mCranberry in dark blue, highlighting differences in the β-bulge region (Gly145-Ser149). **D**, Phenolate-ring (left) and imidazolinone-ring (right) dihedral angles as a function of time for mCherry (purple), mCherry-XL (cyan), mCranberry (red) and mCranberry2 (black) as obtained from MD simulations. The solid lines are the moving averages of the traces. The dihedral angles quantified are shown for each plot. **E**, Representative frames for mCranberry (blue) and mCherry-XL (black) systems, demonstrating subtle differences in packing and water content that improve I-ring stability in mCranberry. For example, Lys73 has 40-50% higher hydrogen bonding occupancy with the O3 atom of the chromophore in mCranberry as compared to all others in the series (Table S9).

The structural analysis therefore demonstrates that substantial improvements in fluorescence quantum yield and photostability can be achieved through mutations distant from the chromophore, supporting the concept that electrostatic effects contribute to fluorescence performance.^19^ Analysis of quantum yield and fluorescence lifetime provides indirect evidence that such improvements might be due to increased radiative rates in mCranberry variants (Fig. 2D). Although the crystal structure does not directly reveal the molecular origin of the reduced non-radiative decay (Extended Data Fig. 5), it provides a structural framework for investigating how distal surface mutations alter the chromophore environment. Ongoing two-photon absorption measurements and NMR studies of mCranberry variants are expected to further elucidate this novel finding.

**Fig 4**| **Crystal structure and MD simulation of mCranberry mutants**. **A,** Representative stereomicroscopic views of protein crystals obtained for mCranberry and mCranberry2. mCranberry crystals were obtained at 15 mg ml⁻¹ protein in 5% (w/v) choline acetate and 30% PEG 400 at pH 7.1, whereas mCranberry-2 crystallized at 30 mg ml⁻¹ protein in 22% (w/v) PEG 3350 and 0.1 M sodium citrate tribasic dihydrate at pH 5.0. **B,** Crystal structure of mCranberry2 determined at a resolution of 3.2 Å, highlighting the chromophore and surface-mutated amino acid residues. **C**, Overlay of representative molecular dynamics structures of wild type mCherry shown in cyan and mCranberry in dark blue, highlighting differences in the β-bulge region (Gly145-Ser149). **D**, Phenolate-ring (left) and imidazolinone-ring (right) dihedral angles as a function of time for mCherry (purple), mCherry-XL (cyan), mCranberry (red) and mCranberry2 (black) as obtained from MD simulations. The solid lines are the moving averages of the traces. The dihedral angles quantified are shown for each plot. **E**, Representative frames for mCranberry (blue) and mCherry-XL (black) systems, demonstrating subtle differences in packing and water content that improve I-ring stability in mCranberry. For example, Lys73 has 40-50% higher hydrogen bonding occupancy with the O3 atom of the chromophore in mCranberry as compared to all others in the series (Table S9).

### Molecular dynamics (MD) simulations of the mCranberry variants

In order to understand the mechanistic implications of mCranberry variants more closely, we modeled a series of systems (mCherry, mCherry-XL, RCB2, K114Q K172H, mCranberry, and mCranberry2) with classical MD. The classical MD primarily revealed key structural differences both in the overall protein β barrel fold and close to the chromophore.

In general, the dynamic fluctuations of all the proteins were relatively small: root mean square fluctuation (RMSF) values for all systems were below 2.5 Å when leaving aside the relatively disordered N/C termini (Figure S5). This is expected as protein rigidity and improved fluorescence quantum yield have been shown to be linked in previous work.^20,30–32^ The largest deviations were seen in the β-bulge region (Gly145-Ser149), where there is a general correlation between increased quantum yield and structural changes in this region (Fig. 4C). mCranberry showed the largest deviation from mCherry.

The consequence of opening this region is twofold. First, the residue Ser146 can hydrogen-bond with the phenolate end of the chromophore (P-ring) in 17-40% of the obtained trajectories (Table S7), which is generally agreed to increase the stability of the P-ring in the excited state. We attribute some of the large increase in quantum yield (QY) of mCherry-XL (QY; 70%) relative to mCherry (QY; 22%) to the resulting stabilization. Secondly, there is an overall increase in hydration within the β-barrel, causing a 10-20% increase in water-bridging hydrogen bonds to Ser143 and Ser146 for K114Q K172H, RCB2, mCranberry, and mCranberry2 (Table S8).

Most of these changes we attribute to the base mutations in mCherry-XL (W143S, I161V, Q163Y, I97R), which generally increase bulk and rigidity within the barrel.^33^ Our fundamental question, however, was the mechanism by which the key surface mutations in the new series, especially K121E for mCranberry, alter the fluorescence quantum yield of the system. To probe this, we first tested the chromophore stability for the imidazolinone (I-ring) and P-ring dihedral angles of the chromophore.

The P-ring is generally stable around 0° for all the systems studied (Fig. 4D, left). The I-ring, however, shows large fluctuations in the initial half of each trajectory from 0 to −35° before converging (Fig. 4D, right). Once converged, K114Q K172H and mCherry-XL— FPs with lower quantum yields--remain around −15°, while RCB2, mCranberry, and mCranberry2—FPs with relatively higher quantum yields--all converge to +5°. Mutating the surface residues close to the I-ring, then, has slightly altered the inner volume of the barrel and subsequent packing, allowing for a more stabilized I-ring. While subtle, even small changes in the respective packing and hydrogen bonding can have substantial effects on the eventual excited state dynamics. We see the relatively larger proportion of hydrogen bonds and water bridges for mCranberry, particularly near the I-ring, as key for its improved quantum yield relative to the other FPs (Fig. 4E, Table S9). Whether TDM of the mCranberry variants are altered via surface mutation as hypothesized before will be investigated with QM/MM calculations.

## Conclusion

In conclusion, here we report two red fluorescent proteins, mCranberry and mCranberry2, exhibiting record quantum yields reported to date, robust in-vitro and in-vivo monomeric character, and low cellular cytotoxicity. Notably, mCranberry2 demonstrates 1.7-fold higher photostability compared to mScarlet3 and has a higher photon budget. While traditional approaches typically modulate RFP quantum yield by rigidifying the chromophore,^34,35^ our findings demonstrate that quantum yields and photostability can also be enhanced through alternative mechanisms. Here, we showcased that strategically introduced charged surface mutations tune chromophore photophysics without compromising its maturation or protein monomericity. Future efforts will involve augmenting their cellular brightness and understanding the mechanism behind surface charge induced change in quantum yield. Overall, this work introduces a unique strategy for tuning photophysical properties of RFPs through surface mutations, successfully enhancing both quantum yield and photostability. Deeper mechanistic insights into these improvements will enable the broader application of this engineering approach to a wider range of FPs and other pigment-binding proteins such as FP-based metal-ion indicators^36,37^ and photosynthetic light-harvesting complexes.^38^

## Methods

### Plasmid construction and mutagenesis

To generate mCranberry variants with improved quantum yield, mCherry-XL was used as the starting template, and pBAD-mCherry-XL served as the parent plasmid. Single-site substitutions were introduced using the Q5 Site-Directed Mutagenesis Kit (NEB, E0554S) with custom primers obtained from Integrated DNA Technologies. To accelerate construction of variants containing multiple substitutions, synthetic DNA fragments spanning the region flanked by NheI and HindIII restriction sites were obtained from Twist Bioscience and ligated into the pBAD backbone using T4 DNA ligase (NEB, M0202S). To generate plasmids for OSER assays, mScarlet3-CytERM-N17 (Addgene plasmid #189778) was used as the recipient backbone. DNA fragments encoding the protein of interest were inserted into this vector using the AgeI and BsrGI restriction sites. For photobleaching and brightness measurements, pDRESS_mTurquoise2_spatial-linker-P2A_mScarlet3 (Addgene plasmid #189752) served as the expression backbone. DNA fragments encoding the protein of interest were cloned into this vector similarly as described above. All plasmid constructs were verified by whole plasmid sequencing (Plasmidsaurus) before use in experiments.

### Protein expression and purification

His-tagged recombinant red fluorescent proteins (RFPs) were expressed and purified from *Escherichia coli* as described previously.^39^ Briefly, electrocompetent *E. coli* Top10 cells were transformed with the pBAD-His plasmid encoding the fluorescent protein of interest using a Bio-Rad Micropulser (Catalog #165-2100). Following transformation, cells were recovered in antibiotic-free 2XYT medium at 37 °C for 1 h, and aliquots were plated onto LB agar supplemented with ampicillin (100 μg ml^-1^) and L-arabinose (0.2%; GoldBio, A-300-25). After overnight incubation at 37 °C, single fluorescent colonies were picked and inoculated into 1 ml 2xYT medium containing ampicillin and grown overnight. The resulting starter culture (1 ml) was then transferred into 100 ml fresh 2xYT medium supplemented with ampicillin and incubated at 37 °C until the culture reached an OD of ∼0.6. Protein expression was induced by addition of L-arabinose to a final concentration of 0.2%, after which cultures were shifted to 28 °C and grown for 20-30 h at 200 rpm to promote protein folding and chromophore maturation.

The induced cultures were harvested by centrifugation at 7,000 rpm for 30 min at 4 °C, and the resulting cell pellets were stored at −80 °C to facilitate lysis. Cells were lysed in B-PER bacterial protein extraction reagent (Thermo Fisher, PI78243) supplemented with EDTA-free protease inhibitor mini tablets (Thermo Fisher, A32955) for 30 min on ice. Lysates were clarified by centrifugation at 10,000 rpm for 1 h at 4 °C, and the supernatant containing the 6×His-tagged protein was filtered through a 0.2 μm syringe filter. The clarified lysate was then incubated with Ni-NTA agarose resin (Cube Biotech, 311103) for 1 h on ice. The resin was loaded onto a column, washed with 100 mM Tris-HCl (pH 7.4) containing 10 mM and then 20 mM imidazole, and bound proteins were eluted with 100 mM Tris-HCl (pH 7.4) containing 250 mM imidazole. Imidazole was removed by buffer exchange into 100 mM Tris-HCl (pH 7.4) using pierce protein concentrators. Purified samples were subsequently used for in vitro photophysical analyses. Proteins were stored at 4 °C for short-term use or flash-frozen in liquid nitrogen and kept at −80 °C for long-term storage.

### Spectral measurements

Absorption spectra were recorded on a Thermo Scientific Evolution One UV–Vis spectrophotometer using a 1 cm quartz cuvette. Measurements were acquired over 250-750 nm with a spectral bandwidth of 1 nm, a data interval of 1 nm, an integration time of 0.5 s, followed by blank subtraction. Samples were diluted in 100 mM Tris-HCl buffer (pH ∼7.4). Excitation and emission spectra were collected on an Agilent Cary Eclipse fluorescence spectrophotometer using a 1 cm path length cuvette. For excitation spectra, emission was monitored at 670 nm while excitation was scanned from 350 to 650 nm using 5 nm excitation and emission slit widths and a scan rate of 120 nm/min. For emission spectra, samples were excited at 520 nm, and emission was recorded from 535 to 800 nm using 5 nm excitation and emission slit widths and a scan rate of 120 nm/min.

### Extinction coefficient

The extinction coefficient of the RFP variants was determined via the alkali denaturation method as previously described.^20^ Absorbance spectra of the purified RFPs in 100 mM Tris-HCl buffer (pH 7.4) were acquired using a Thermo Scientific Evolution One UV-Vis spectrophotometer. All measurements were performed in a 1 cm quartz cuvette across a wavelength range of 250-750 nm, utilizing a 1 nm spectral bandwidth, a 1 nm data interval, and an integration time of 0.5 seconds. Typically, protein samples were prepared in parallel under denatured and native conditions using a starting volume of 1.2 ml of protein stock solution OD ∼ 0.11. For the denatured sample preparation, a 500 μl aliquot of the protein stock was transferred to a designated tube and mixed with 30 μl of 10 M NaOH. The absorbance spectrum was subsequently recorded, using a baseline blank consisting of 500 Tris buffer mixed with 30 μl of 10 M NaOH. Concurrently, for the native sample preparation, a separate 500 μl aliquot of the protein stock was transferred to a native sample tube and mixed with 30 μl of Tris buffer. The absorbance spectrum for the native protein was then recorded using Tris buffer alone as the baseline blank. The denatured FPs exhibit an absorption peak with known extinction coefficients of 44,000 M^-1^ cm^-1^ near 450 nm. The value of the peak extinction coefficient was determined according to the ratiometric relationship expressed below:

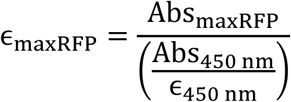

### Quantum yield

Fluorescence quantum yield of the RFPs were measured by serial dilution method as described previously.^24^ Briefly, absorbance spectra of the FPs were acquired using a Thermo Scientific Evolution one UV-Vis spectrophotometer and a 2 cm pathlength quartz cuvette, followed by baseline correction. ^24^ Briefly, absorbance spectra of the FPs were acquired using a Thermo Scientific Evolution One UV-Vis spectrophotometer and a 2 cm pathlength quartz cuvette, followed by baseline correction. For each RFP variant, a dilution series consisting of 4-5 different concentrations were prepared. The most concentrated sample was adjusted to yield a maximum optical density OD_520_ of approximately 0.15 in the 2 cm cuvette. OD_520_ of the step-dilution series were measured and subsequently integrated fluorescence of the samples were obtained across the entire emission band (535 - 800 nm). Emission spectra were recorded on a Cary Eclipse fluorescence spectrophotometer using a 1 cm pathlength cuvette and the following parameters: excitation at 520 nm, emission scanning from 535 to 800 nm, excitation and emission slit widths of 2.5 nm, and a scan speed of 120 nm/min. All emission spectra were baseline-corrected, and the integrated fluorescence was quantified by calculating the area under the curve. To determine the quantum yield, the integrated fluorescence values were plotted against the corresponding OD_520_ values. The resulting data were fit to a straight line with the regression lines constrained through the origin, according to the following equation: *y* = *slope* × *x* where *y* is the integrated fluorescence and *x* is the OD_520_. Finally, fluorescence quantum yield ( *φ_RFP_*) of the RFPs were calculated from the below equation:

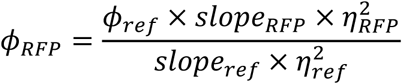

where *η_RFP_* and *η_ref_* are the refractive index of the RFP and the reference sample. Here, mScarlet (*φ* = 0.71) are used as the reference for all protein except FusionRed and TagRFP-T for which mCherry (*φ* = 0.22) was used as reference.

### pKa measurement

For pKa measurements, a series of buffers spanning pH 2-12 was prepared, and the pH of each solution was verified using a calibrated pH meter. Fixed amounts of concentrated pure protein samples were added to 1 ml of buffer in a quartz cuvette with a 1 cm path length, and a fluorescence spectrum was recorded in each case with excitation at 520 nm. After addition of proteins the samples were left at room temperature for an hour to reach equilibrium. Instrument settings used for these measurements were: Excitation: 520 nm, Emission: 535-800nm, slit width (both): 5 nm, speed: 120 nm/min). To account for variations in emission maxima across pH conditions, individual emission spectra were normalized relative to the maximum fluorescence intensity observed across all tested pH buffers. Normalized fluorescence values were plotted as a function of pH and fitted in GraphPad Prism using a sigmoidal dose-response model with variable slope [log(agonist) versus normalized response]. The resulting fits were used to calculate pKa values.

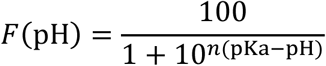

For curve fitting, *F*(pH) denotes the normalized peak fluorescence of the sample at a given pH, and *n* represents the Hill slope, which describes the steepness of the fitted sigmoidal transition and pH is the independent variable (x-axis). In plots of peak fluorescence versus solution pH, the pKa of each fluorescent protein was defined as the pH at which the fluorescence intensity reached half of its maximum value. The pKa value for each RFP was determined from three biological replicates. Buffers below pH 6 were prepared from 0.1 M sodium citrate and 0.1 M citric acid. Buffers between pH 6 and 8 were prepared using 0.2 M potassium dihydrogen phosphate and 0.2 M disodium hydrogen phosphate, whereas buffers above pH 8 were prepared from 0.2 M glycine and 0.2 M sodium hydroxide.

### Absorption spectra at varying pH

Series of buffers ranging from pH 3.0 to 12.0 were prepared as previously detailed in pKa measurement section. Protein samples were prepared by adding fixed amounts of concentrated stock to 1.5 ml of the respective buffer in a 4 cm path-length quartz cuvette. To allow full equilibration, the mixtures were maintained at room temperature for 5-6 h. Absorbance spectra were subsequently collected on a Thermo Scientific Evolution One spectrophotometer from 250 to 750 nm (bandwidth: 1 nm; data interval: 1 nm; scan speed: 117.65 nm/min; integration time: 0.5 s). Background subtraction was performed using the corresponding buffer controls.

### Fluorescence lifetime

Fluorescence lifetime of the purified proteins was measured using a home-built time-correlated single-photon counting (TCSPC) setup. The setup uses a 515 nm pulsed laser (LDH-D-C-520, PicoQuant, Germany) operated at 40 MHz using a PDL 828 “Sepia II” driver (PicoQuant, Germany) to excite the samples. Emitted photons are collected with an avalanche photodiode (APD; SPCM-AQRH-14-TR, Excelitas), which is connected to a TCSPC module, Picoharp 330 (PicoQuant, Germany). For the measurement, 30 µl purified protein (in 1XPBS) was placed in a precision cover glass #1.5 (CG15KH1, Thorlabs), the laser was focused 25 µm into the sample with the help of a Piezostage (BIO3, Piezoconcept, France) using an Olympus UplanSApo 100X oil immersion objective. The filters and dichroic mirrors (DM) used for the measurement include a laser cleanup filter: FBH515-10 (ThorLabs); DM1: ZT532/640/NIR_rpc (Chroma); Emission filter: ET630/75m (Chroma). The detailed schematic of the setup is shown in Supplementary Figure 3. The resulting fluorescence decays were fit to a single-exponential function by the iterative reconvolution method. The fit values are shown in Table 2.

### Oligomerization analysis by Size Exclusion Chromatography (SEC)

Purified RFP samples were buffer-exchanged into 100 mM Tris-HCl, 100 mM NaCl, pH 7.4, which also served as the SEC running buffer. A 500 μl aliquot of concentrated protein was loaded onto a pre-equilibrated Superdex 75 10/300 GL column mounted on an FPLC system. Chromatographic separation was performed at a flow rate of 0.3 ml min^-1^, and protein elution was monitored by absorbance at 280 nm. tdTomato, produced in-house, was used as a dimeric standard, whereas mScarlet3 served as a monomeric standard. A gel filtration standard (Biorad,1511901) was also included as a molecular size marker.

### Cellular cytotoxicity

Cytotoxicity induced by RFP expression in U2OS cells was observed using a Cell Counting Kit-8 (Sigma-Aldrich, 96992). This colorimetric assay quantifies viable cells by measuring the reduction of WST-8 to water-soluble formazan with absorbance read at 460 nm. On day 0, 3,000 U2OS cells were seeded per well in 100 µl of DMEM supplemented with 10% fetal bovine serum and 100 U ml⁻¹ penicillin-streptomycin. Cells were plated in Falcon 96-well clear flat-bottom TC-treated culture microplates (353072) and maintained at 37 °C in a humidified incubator with 5% (v/v) CO₂. On day 1, 100 ng CytERM-RFPs and pDRESS_mTurquoise2_spatial-linker-P2A_RFPs were transiently transfected using Lipofectamine 3000 (Invitrogen, L3000001) following the manufacturer’s instructions. Six wells were seeded with untransfected U2OS cells to serve as positive controls, and wells containing 100 µl of DMEM alone were used as blanks. After transfection, the 96-well plate was returned to the humidified cell incubator. Diluted mix of cell counting kit was prepared by adding 1 ml WST-8 mix from cell counting kit-8 and 4.5 ml of DMEM. 50 μl of this cell counting mix was added in each well on day 2 for CytERM-RFPs and on day 3 for pDRESS_mTurquoise2_spatial-linker-P2A_RFP plasmid followed by 2 h incubation at 37 °C and 5% CO_2_. Absorbance at 460 nm was recorded using a SpectraMax M5 plate reader. The mean absorbance (± s.d.) of six replicate wells transfected with the same plasmid was calculated and blank-corrected to quantify cellular cytotoxicity.

### Assessment of protein oligomerization using OSER method

U2OS cells were routinely maintained in DMEM (Thermo Fisher Scientific, 10566-016) supplemented with 10% fetal bovine serum (Thermo Fisher Scientific, A52567) and 100 U ml⁻¹ penicillin-streptomycin (Thermo Fisher Scientific, 15-140-122) at 37 °C in a humidified 5% CO₂ atmosphere. For imaging experiments, cells were seeded in 35 mm cell culture microscopy dishes with a #1.5 polymer coverslip (ibidi, 81156). CytERM constructs were transiently transfected with 1 µg plasmid DNA using Lipofectamine 3000 (Invitrogen, L3000001) for 24 h, following the manufacturer’s instructions. Before live-cell imaging, cells were transferred to 1× DMEM (Thermo Fisher Scientific, 31053-028) supplemented with 10% fetal bovine serum and 100 U ml⁻¹ penicillin-streptomycin. Analysis of approximately 300 cells was sufficient to obtain a robust OSER score. Imaging was performed in an environmentally controlled chamber (Oko Labs; 37 °C, 5% CO₂) on a Nikon A1R confocal system (Nikon Ti-E) using a 20×Plan Apo Lambda dry objective (NA 0.75) with 2x optical zoom. Samples were illuminated by a 561 nm laser, and emission was collected using a 595/50 nm bandpass filter, with an acquisition rate of 3.9 frames per second (frame time 259.9 ms). Large fields (1024 × 1024 pixels) were acquired while scanning in z-stacks to ensure optimal focus for all cells in the field of view. The typical z-range was 8 µm, sampled with a 1 µm step size, yielding 9 layers. Following the acquisition, maximum intensity projections of the z-stacks were generated using ND Processing, and these projections were used for quantification of the OSER score. The OSER score is calculated from at least three biological replicates for each CytERM construct. To quantify the OSER, we developed an automated image analysis pipeline utilizing CellProfiler,^40^ with a summary of the pipeline workflow and its comprehensive parameters presented in Supplementary Figure 2.

### Brightness in Mammalian Cells

The brightness of the diverse RFPs was determined in U2OS cells. U2OS cells were routinely maintained as described in the assessment of protein oligomerization using OSER method. For imaging experiments, cells were seeded in 35 mm cell culture microscopy dishes with a #1.5 polymer coverslip (ibidi, 81156). During passage, cells were seeded at a density of 2.5 x10^5^ cells ml^-1^ in a 400 µl volume at the center of the microscopy dish. Following a 20-minute incubation period to allow for initial attachment, the volume was brought to 2.0 ml of complete medium. Transient expression of the bicistronic pDRESS_mTurquoise2_spatial-linker-P2A_RFP vectors (1:1 mTurquoise2:RFP ratio) was achieved using 1 μg of plasmid DNA and Lipofectamine 3000 (L3000001, Invitrogen) for 48 h following the manufacturer’s instructions.

Before live-cell imaging, cells were transferred to 1× DMEM supplemented with 10% fetal bovine serum and 100 U ml⁻¹ penicillin-streptomycin. Imaging was performed in an environmentally controlled chamber (Oko Labs; 37 °C, 5% CO₂) on a Nikon A1R confocal system (Nikon Ti-E) under resonant scanning mode. Fluorescence imaging was performed 48 h post-transfection using a confocal microscope equipped with 402, 487, and 561 nm laser lines. Triple-channel acquisition was configured for cyan, yellow, and red using emission collection windows of 425-475 nm, 500-550 nm, and 570-620 nm, respectively. Images were captured at a resolution of 1024 x 1024 pixels using a 10x Plan Apo Lambda dry objective (NA 0.45, WD 4 mm). Laser power at the objective plane was measured with a Thorlabs PM400 optical power meter and was maintained at 70 µW (402 nm), 250 µW (487 nm), and 38 µW (561 nm) to optimize the signal-to-noise ratio while reducing potential phototoxicity.

The red-to-cyan fluorescence intensity ratios were quantified using a custom-developed CellProfiler pipeline (Supplementary Figure 5).^40^ To determine the relative brightness within mammalian cells, these raw ratios were normalized by accounting for the specific spectral throughput and detection efficiency of the imaging system.

### Confocal photobleaching in fixed *E.Coli*

*E.coli* strains with either the pBAD-His or pDx-mScarlet3 plasmid were streaked onto agar plates and incubated to isolate single colonies. A single colony was inoculated into 5 ml of 2xYT medium containing the appropriate selection of antibiotics (ampicillin for pBAD-His; kanamycin for pDx-mScarlet3) and grown overnight. To initiate expression cultures, 50 µl of the overnight culture was inoculated into 5 ml of fresh 2xYT medium supplemented with the corresponding antibiotic and grown until reaching an OD600 of approximately 0.6. Protein expression was subsequently induced by the addition of 0.2% (w/v) L-arabinose (GoldBio, A-300-25) for pBAD-His or 0.2% (w/v) L-rhamnose for pDx-mScarlet3. Cultures were then incubated for an additional 24 h at 28 °C. Following induction, 100 µl of cell culture was harvested, washed three times with 1X PBS, and resuspended in 1X PBS buffer for confocal photobleaching experiments. 5 μl of cell suspension in 1X PBS were mounted on Poly-L-Lysine-coated slides (Expredia, 6776216) using microscope coverslips (VWR, 16004-302). Confocal imaging was performed on a Nikon A1R microscope operated in resonant scanning mode. Samples were visualized using a 60x oil-immersion Plan Apo Lambda objective (NA 1.4, WD 0.13 mm) at a 1024 × 1024-pixel resolution. Regions of interest (ROIs) corresponding to individual cells and background regions were selected under minimal laser power to prevent initial RFP photobleaching. Time-series acquisitions were executed in ND Acquisition mode using continuous 561 nm laser excitation at 42 μW laser power at the objective plane, measured with a Thorlabs PM400 optical power meter. ROIs were continuously illuminated for 180 s, during which 694 images were captured. Mean fluorescence intensity within each ROI was tracked over time, corrected for background fluorescence at each time point, and normalized to the initial intensity at *t* = 0 s. Photobleaching half-times (*t*_1/2_) were defined as the duration required for fluorescence emission to decay to 50% of its initial value. To correct for intrinsic differences in extinction coefficient, quantum yield, and excitation efficiency, raw time coordinates were scaled relative to mScarlet3 by multiplying each trace by its relative molecular brightness factor (B_RFP_/B_mScarlet3_) as described previously.^29^

### Confocal photobleaching in mammalian Cells and Photon Budget

U2OS cells were maintained, and transfections were performed using pDRESS_mTurquoise2_spatial-linker-P2A_RFP vectors as described in the Brightness in mammalian cells method section. Before live-cell imaging, cells were transferred to 1× DMEM supplemented with 10% fetal bovine serum and 100 U ml⁻¹ penicillin-streptomycin. Imaging was performed in an environmentally controlled chamber (Oko Labs; 37 °C, 5% CO₂) on a Nikon A1R confocal system (Nikon Ti-E) under resonant (Update) scanning mode. Fluorescence imaging was performed 48 h post-transfection. The samples were imaged using a 40x oil-immersion Plan Fluor objective (NA 1.3, WD 0.2 mm) at 512 × 512-pixel resolution. A Perfect Focus System (PFS) was used throughout acquisition to prevent focus drift. Multiple cells were selected as regions of interest (ROIs) using the cyan channel under low laser power to minimize RFP photobleaching prior to confocal bleaching, and background regions were also selected for correction. In ND Acquisition time-measurement mode, the 561 nm laser was operated at an objective-plane power of 0.7 mW, which was measured using a Thorlabs PM400 optical power meter. Under continuous 561 nm excitation with PFS enabled, ROIs were illuminated for 2 min, and 923 images were acquired. The mean ROI intensity was measured over time for each cell, corrected for background fluorescence at each time point, and normalized to the intensity at t = 0 s before plotting. Photobleaching half-times (t_1/2_) were determined by numerical interpolation as the time required for fluorescence emission to decay to 50% of its initial value. Photostability measurements were conducted across three biological replicates (independent transfections) for each construct. Photobleaching decay traces are presented as mean curves, and half-times (t_1/2_) were calculated individually for each replicate and reported as mean ± s.d. (n=3). As excitation irradiance was kept constant across all measurements, raw time coordinates were scaled relative to mScarlet3 by multiplying each trace by its relative molecular brightness (B_RFP_/B_mSc-3_) to correct for intrinsic differences in extinction coefficient, quantum yield, and excitation efficiency as described previously.^29^

Photon budgets were quantified from continuous photobleaching time-lapse series. Raw fluorescence intensity decay curves over time (seconds) were first obtained for individual regions of interest (ROIs). To account for system-specific optical efficiency, measured intensity values were multiplied by the total spectral throughput of the imaging system. The total photon budget for each individual cell or ROI was subsequently determined by calculating the integrated area under the corrected fluorescence decay curve over the duration of the photobleaching experiment. Single-cell photon budget distributions were then combined and visualized using violin scatter plots to evaluate the overall photon emission capacity and photostability of the fluorescent proteins.

The red-to-cyan fluorescence intensity ratios were quantified using a custom-developed CellProfiler pipeline.^40^ To determine the relative brightness within mammalian cells, these raw ratios were normalized by accounting for the specific spectral throughput and detection efficiency of the imaging system.

### Structure Determination

A batch of mCranberry, mCranberry2, and K114Q K172H protein samples were specifically expressed and purified for crystallization studies as described below.

### Protein purification

Protein expression was performed as described in the Methods section. For cell disruption, a non-chemical lysis approach was used with an EmulsiFlex homogenizer. Thawed cell pellets were resuspended in 100 mM Tris-HCl (pH 7.4) supplemented with EDTA-free Protease Inhibitor Mini Tablets (Thermo Fisher, A32955). The suspension was passed through the EmulsiFlex at least three times at 15,000–20,000 psi to achieve complete lysis. Lysates were clarified by centrifugation at 10,000 rpm for 1 h at 4 °C, and the supernatant containing the 6x His-tagged protein was filtered through a 0.2 μm syringe filter. The clarified lysate was then incubated with Ni-NTA agarose resin (Cube Biotech, 311103) for 1 h on ice. The resin was loaded onto a column, washed with 100 mM Tris-HCl (pH 7.4) containing 10 mM and then 20 mM imidazole, and bound proteins were eluted with 100 mM Tris-HCl (pH 7.4) containing 250 mM imidazole. Imidazole was removed by buffer exchange into 20 mM Tris-HCl, 50 mM NaCl, 2 mM CaCl_2_ (pH 8.0) using Pierce protein concentrators.

The N-terminal His tag was removed by digestion with Enterokinase, light chain (NEB, P8070) for 16 h at 25 °C. To separate cleaved from uncleaved protein, the reaction mixture was subjected to a second round of Ni-NTA affinity purification step, in which the cleaved protein was collected in the flow-through and uncleaved protein was retained in the elution fraction. The flow-through fractions were pooled and buffer-exchanged into Buffer A (20 mM Tris-HCl, pH 7.4). The concentrated protein sample was loaded onto a Resource Q anion-exchange column (Cytiva, 17117901) on an FPLC system at a flow rate of 0.6 ml min⁻¹, and elution was monitored by absorbance at 280 nm. Bound protein was eluted with Buffer B (20 mM Tris-HCl, 1 M NaCl, pH 7.4) using a linear gradient. Fractions containing the target protein were pooled and buffer-exchanged into 50 mM HEPES (pH 7.9) for crystallization trials.

### Methods of mCranberry2 Crystallography, Diffraction, and Analysis

Initial conditions were identified by sparse-matrix screening in 96-well plates (Greiner Bio-One, 07-000-158) using a mosquito crystallization robot (TTP LabTech) in hanging-drop format. Initial hits were subsequently optimized in 24-well VDX plates (Hampton Research, HR3141) using siliconized square cover slides (Hampton Research, HR3-215) at room temperature. mCranberry crystals were obtained at 15 mg ml⁻¹ protein in 5% (w/v) choline acetate and 30% PEG 400 at pH 7.1, whereas mCranberry2 crystallized at 10-15 mg ml⁻¹ protein in 25% (w/v) PEG 3350 and 0.1 M sodium citrate tribasic dihydrate at pH 5.0. For mCranberry2, reservoir solutions contained 0.1 M sodium citrate tribasic dihydrate at pH 5.0-6.0 and 15-30% (w/v) PEG 3350. Drops were prepared by mixing reservoir solution with protein solution at 1:1 and 1:2 (v/v) ratios and equilibrated against 1 ml of reservoir. Protein concentrations of 10-30 mg ml^-1^ were screened. The crystals used for data collection for mCranberry2 grew from 0.1 M sodium citrate tribasic dihydrate, pH 5.0, 25% (w/v) PEG 3350, at a protein concentration between 10-15 mg ml^-1^ with crystals confirmed after 13 days and subsequently diffracted in the next 2-10 days. The morphology of crystals was rod-like. All crystals of sufficient quality were placed on 0.1 – 0.4 mm Mounted CryoLoops (Hampton Research), briefly (1-2 seconds) incubated in polyfluoroether cryo-oil (Hampton Research), rapidly frozen in liquid nitrogen, and placed into a Stanford Synchrotron Radiation Lightsource (SSRL) cassette or loaded directly on the beamline for X-ray diffraction data collection. X-ray diffraction was performed at SSRL (Menlo Park, CA) Beam Line 12-2 at 100K. Further detailed information on the data collection strategy can be found in Romei et al.^41^ Data processing and scaling were performed using the autoproc software^42^,Further detailed information on the data collection strategy can be found in Romei et al.^41^ Data processing and scaling were performed using the autoproc software^42^ providing the correct C2 space group. Molecular replacement was performed in Phenix-1.21.2^43^ with phenix.phaser using the mCherry (PDB Entry: 2H5Q)^23^ structure as the search model. Numerous rounds of model building and refinement were carried out in Coot^44^ and then with phenix.refine. Refinement continued until the R-free score could no longer be lowered. The X-ray collection parameters and refinement statistics are given in Table S6.

### Computational Methods

#### Classical MD

Classical molecular dynamics (MD) simulations were performed in triplicate for the following systems: mCherryXL,^20^ mCranberry, mCranberry2, K114Q K172H, and RCB2. Previously published MD simulations for wild-type mCherry, using the same parameters, were taken from previous work as a comparison.^13^

Each system was prepared in two ways: first, by using the appropriate Dunbrack rotamers to replace residues and rotate sidechains from the wild-type mCherry crystal structure (PDB ID: 2H5Q).^23,45^ Second, by using the crystal structure obtained in this work for mCranberry2 and replacing the modified residues in the same manner for each system. N/C termini sequences for all systems were replaced using AlphaFold3.^46^ The crystal structure of mCranberry2 and the simulations of mCranberry2 developed from wild type mCherry matched very closely (sub 1 Å RMSD), and there were not large deviations between systems, so we consider them as replicate simulations in our analysis.

Once initial structures were generated, they were initially protonated and hydrogen-bonding networks optimized using MolProbity.^47,48^ Each system was checked with propka3 to ensure correct residue protonation states.^49^ Chromophore force field parameters were generated with AutoParams in previous work.^13,50^ The rest of the protein was simulated with the ff14SB force field, with solvent simulated as TIP3P water.^51,52^ Systems were neutralized to a net charge of 0 by adding 7 or 8 Na^+^ ions depending on the mutant. Each system was simulated in triplicate for 1 μs each, for a total of 3 μs of simulation time per system and 15 μs of simulation overall, using the pmemd.cuda module of the Amber24 simulation package.^53^ After an initial solvent density equilibration, heating, and re-equilibration, production was run in the NVT ensemble using a 2 fs timestep and the SHAKE/RATTLE algorithms for hydrogens.^54^ A Langevin thermostat was used with the friction parameter set to 5.0 ps^-1^, and smooth particle mesh Ewald was used for electrostatics, with a 10 Å nonbonded interaction cutoff.^55,56^ Simulations were run with periodic boundary conditions.

## Author Contributions

P.M., S.C., and S.M. conceptualized the study. A.A. and A.R.W. designed, performed, and analyzed the simulations. S.C., P.M., S.M., M.F., H.K.R.R.B. and I.I.W. designed and performed the experiments. P.M. and A.R.W. supervised the study. S.C. and P.M. wrote the first draft of this manuscript.

## Notes

The authors declare no competing financial interests.

## Acknowledgement

S.C. was supported by the Faculty Start-up Grant PG100125 from The Ohio State University (OSU), awarded to P.M. A. A. was supported by R35GM154949, awarded to A.R.W. S.M. was partially supported by NIH Grant R35GM118044 to Prof. Steven G. Boxer (Stanford). Use of the Stanford Synchrotron Radiation Lightsource (SSRL/SLAC) was supported by the DOE Office of Science under Contract DE-AC02-76SF00515 and NIH Grant P30GM133894. P.M. thanks Prof. Ralph Jimenez (JILA, NIST & University of Colorado Boulder), Pia Friis (JILA), and Nancy Douglas (JILA) for providing the RCB2 and other FP constructs and for valuable discussions. P.M. and S.C. thank Prof. Vladislav Belyy (OSU) for providing U2OS cells, Prof. Charles Bell (OSU) for helping with crystallization, and Prof. Thomas Magliery (OSU) for providing the FPLC facility. The authors thank Prof. Amy E. Palmer (University of Colorado Boulder), Prof. Sheng-Ting Hung (NSYSU), Prof. Mark Foster (OSU), Prof. Venkat Gopalan (OSU), and Prof. Steven G. Boxer (Stanford) for valuable discussions.

## Extended Data

### 1. Shifts in absorption and emission spectra

**Extended Data Fig. 1.**
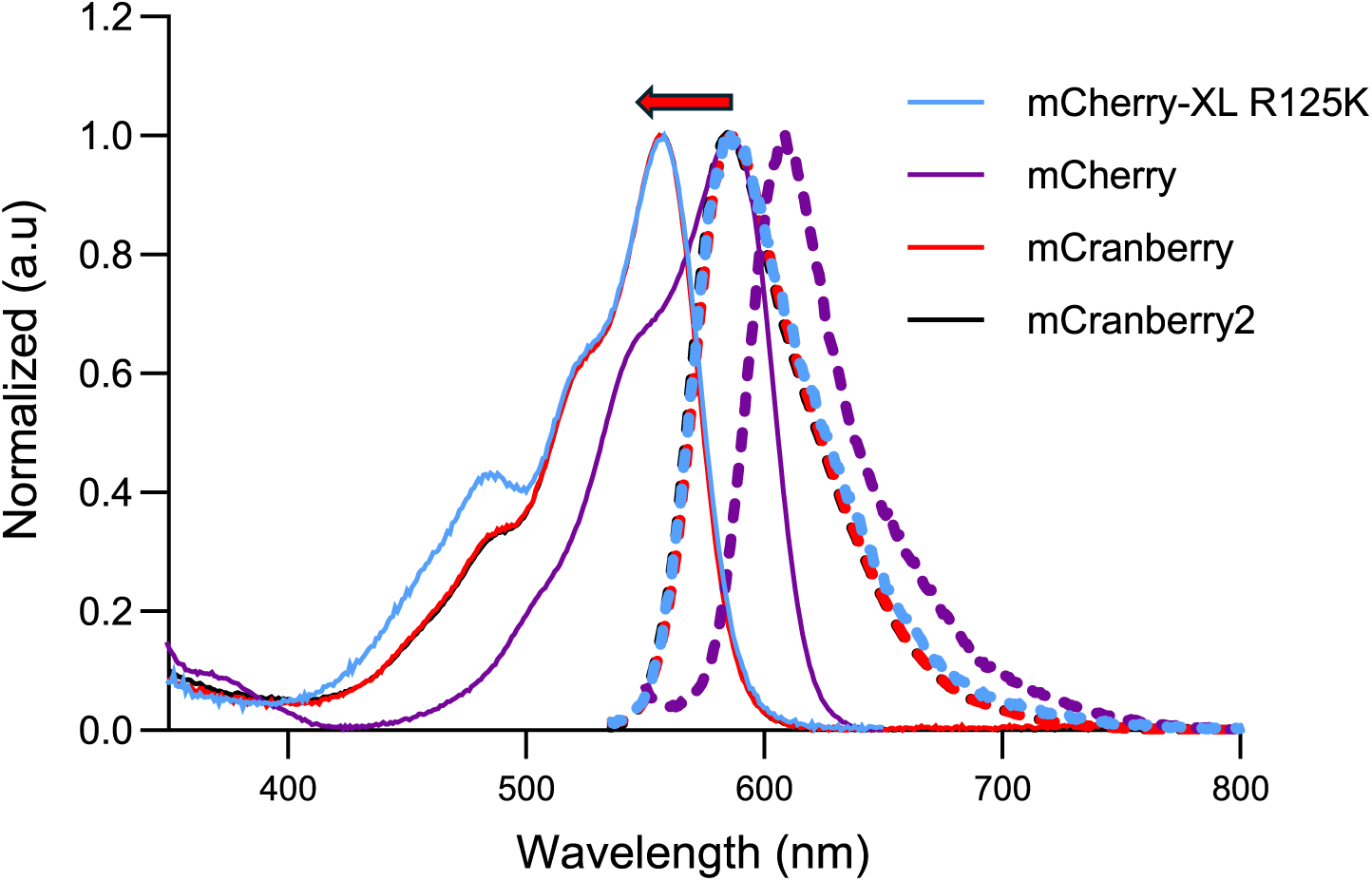
Shifts in absorption and emission spectra. Normalized absorption (solid lines) and emission spectra (dashed lines) of mCranberry and parent mutants are shown. Compared to mCherry, the absorption and emission maxima of mCherry-XL R125K, mCranberry, and mCranberry2 are blue-shifted.

### 2. pH dependence changes in absorption and emission

**Extended Data Fig. 2.**
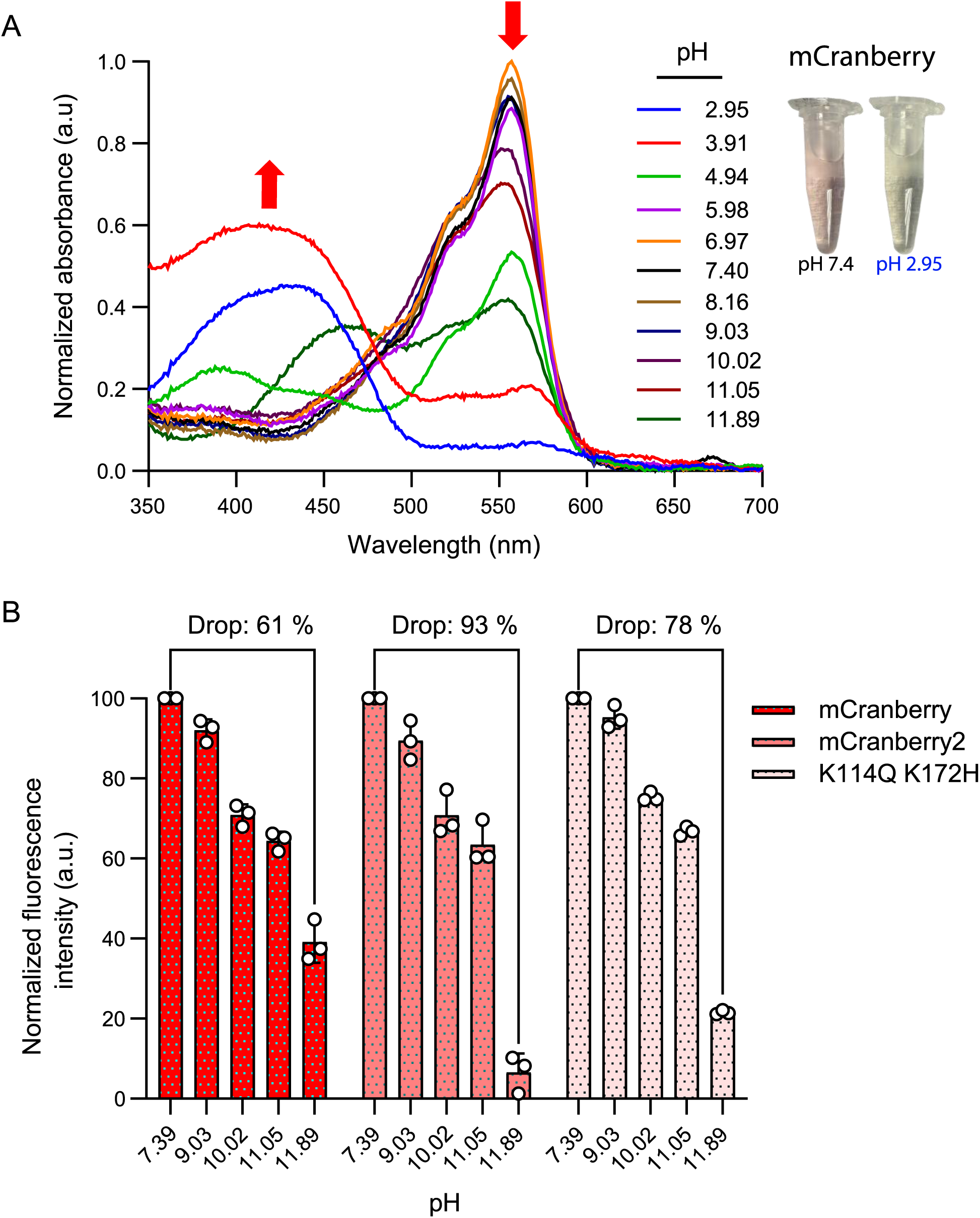
pH Dependence changes in absorption and emission. **A,** pH-dependent absorption spectra of mCranberry. Normalized absorbance of the mCranberry was recorded at varying pH levels ranging from 2.95 to 11.89. **B,** Comparison of fluorescence intensity loss at alkaline pH across mCranberry variants. A bar graph illustrates the normalized fluorescence intensity of three independent biological replicates. Fluorescence for each FP is normalized to an initial maximum of 100 at pH 7.39. The annotations above each dataset represent the total percentage decrease in fluorescence intensity when transitioning from pH 7.39 to pH 11.89.

### 3. Quantum yield and Extinction Coefficient

**Extended Data Fig. 3.**
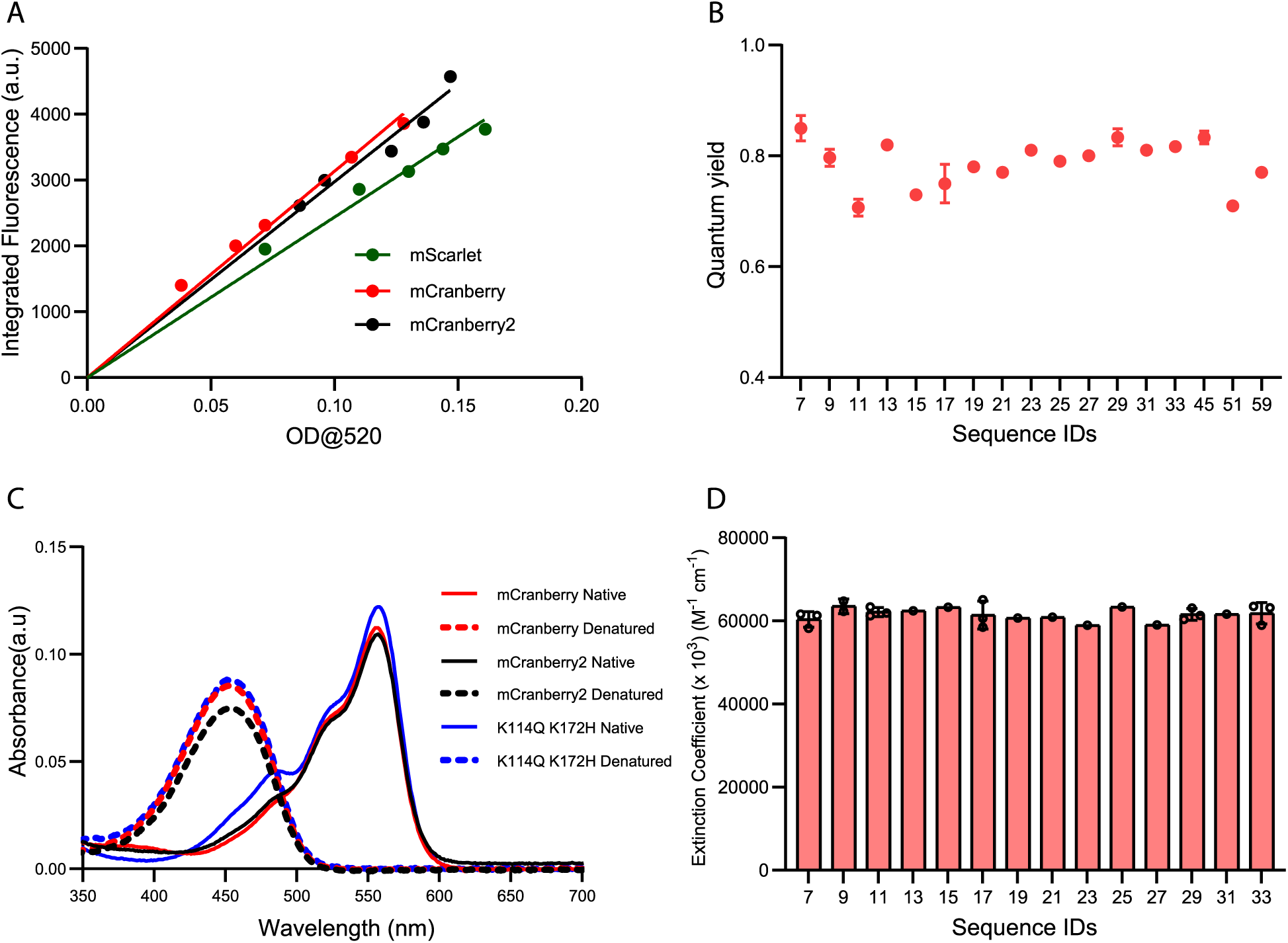
Quantum yield and Extinction coefficient measurements. **A,** Linear regressions of integrated fluorescence emission (535–800 nm) versus absorbance (OD_520_) across serial dilutions of mCranberry mutants and reference standard mScarlet. Data shown are representative of one of three biological replicates, while relative quantum yields in the main text represent values from all three replicates. mCranberry shows the steepest slope consistent with its highest quantum yield. **B,** Relative quantum yields of mCranberry variants from the fourth round of evolution. Sequence IDs correspond to individual protein mutants listed in Supplementary Table 1. Key mutants include Seq ID7 (mCranberry), Seq ID9 (mCranberry2), and Seq ID51 (mScarlet). **C,** To determine the peak extinction coefficients of the mCranberry variants, the alkali-denaturation method was employed to compare the absorbance profiles of the proteins in their native and denatured states. Upon the addition of 10 M NaOH (dashed lines), complete denaturation of the beta-barrel structure was achieved for all three variants, resulting in a hypsochromic shift. The native major absorption peak at ∼557 nm was entirely reduced, replaced by a distinct denatured peak centered at approximately 450 nm, which corresponds to the denatured anionic chromophore state. **D**, Extinction coefficient of mCranberry variants from the fourth round of evolution. Sequence IDs correspond to individual protein mutants listed in Supplementary Table 1. Key mutants include Seq ID (mCranberry), Seq ID9 (mCranberry2).

### 4. Excited-state lifetime

**Extended Data Fig. 4.**
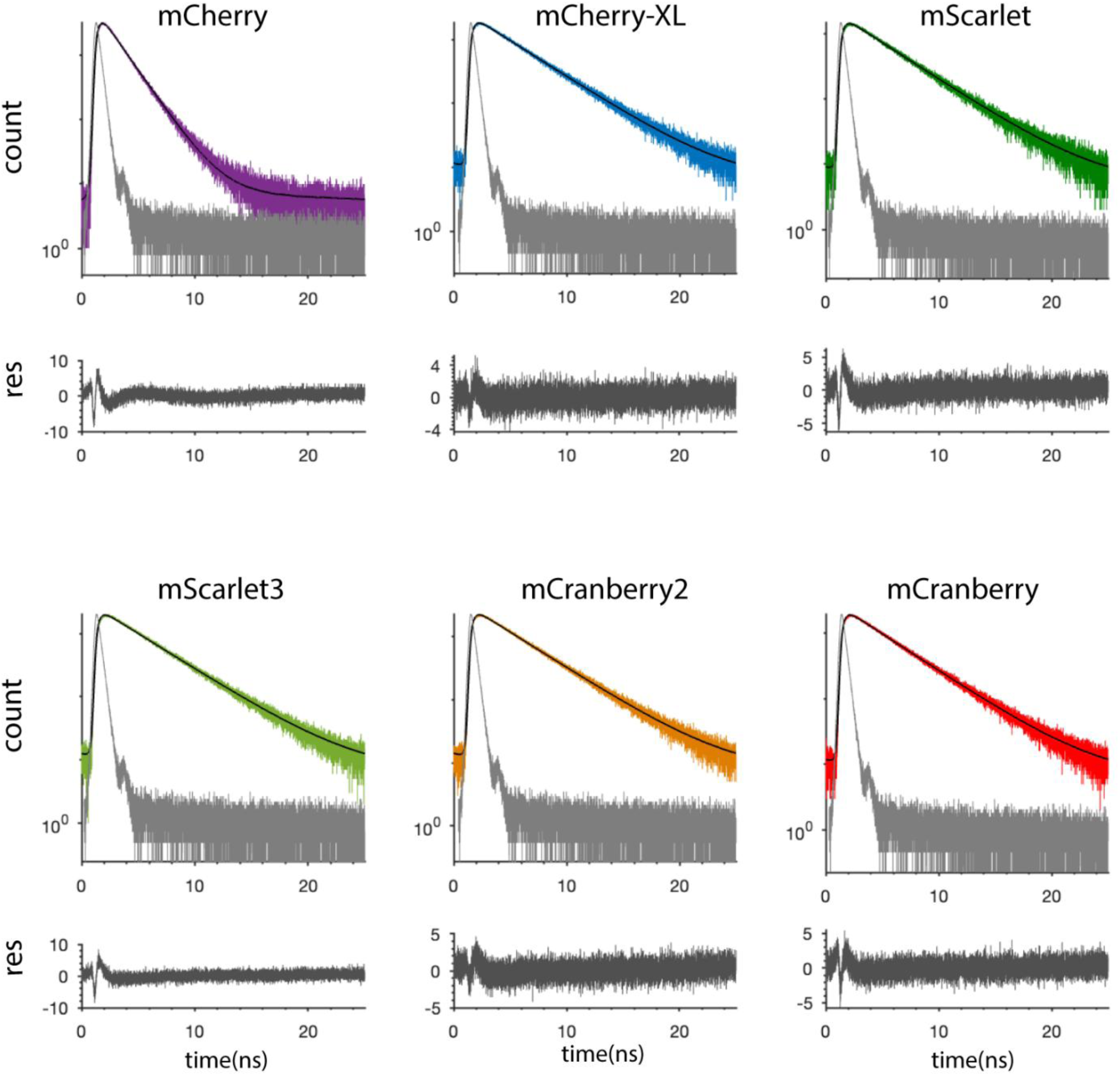
Fluorescence lifetime analysis of the purified protein mutants. Experimentally measured lifetime decays (colored curves) and their respective single-exponential fits (black curves) are presented for each protein. The instrument response function is indicated in gray, with the corresponding residuals displayed at the bottom. The fitted lifetime values are presented in Table 2, main text.

### 5. Changes in Non-radiative rate constant

**Extended Data Fig. 5.**
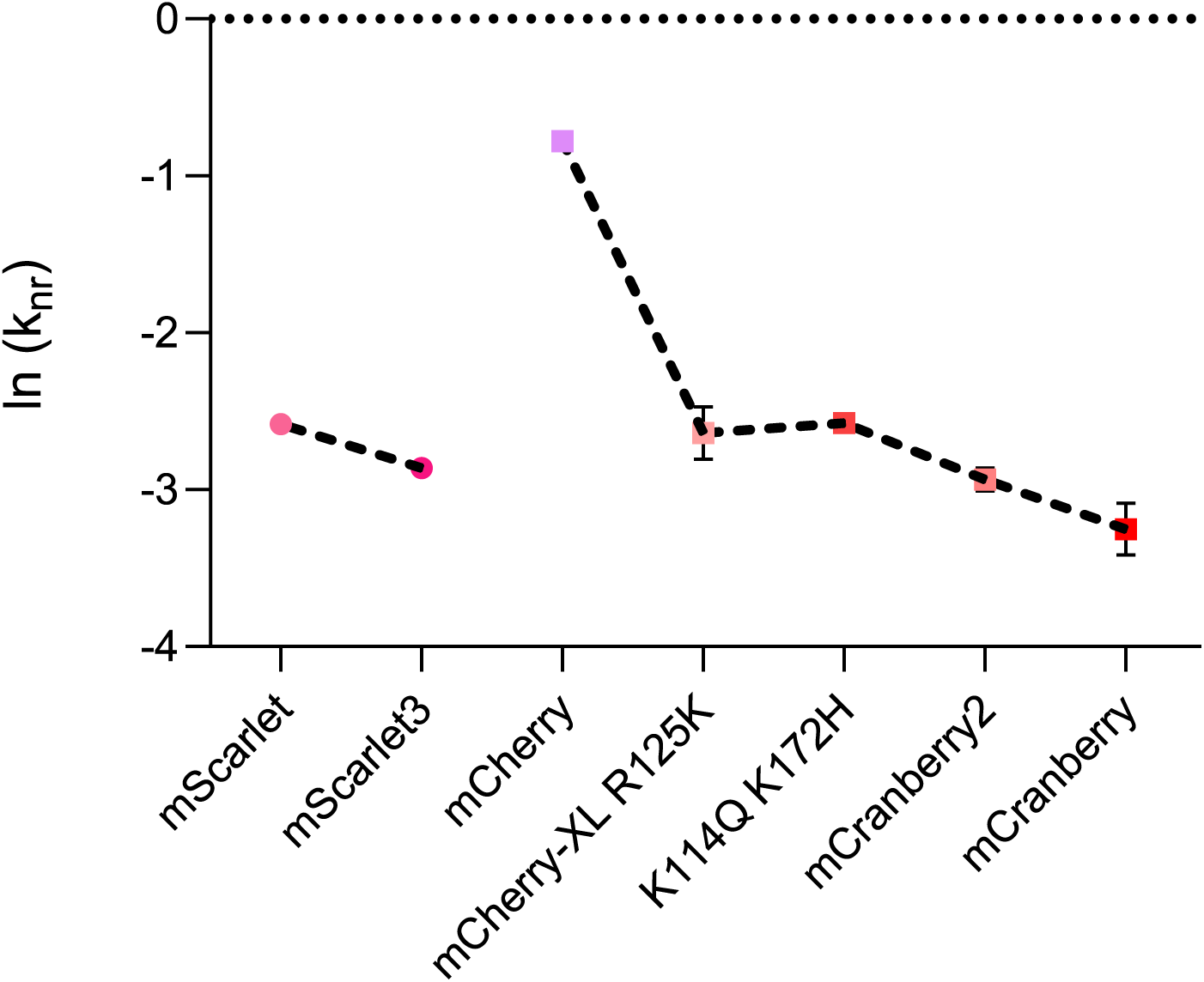
Non-radiative rate constants. Logarithmic non-radiative decay rates (ln *k*_nr_) across red fluorescent protein (RFP) variants was calculated from the quantum yield and lifetime values.

### 6. Confocal Photobleaching

**Extended Data Fig. 6.**
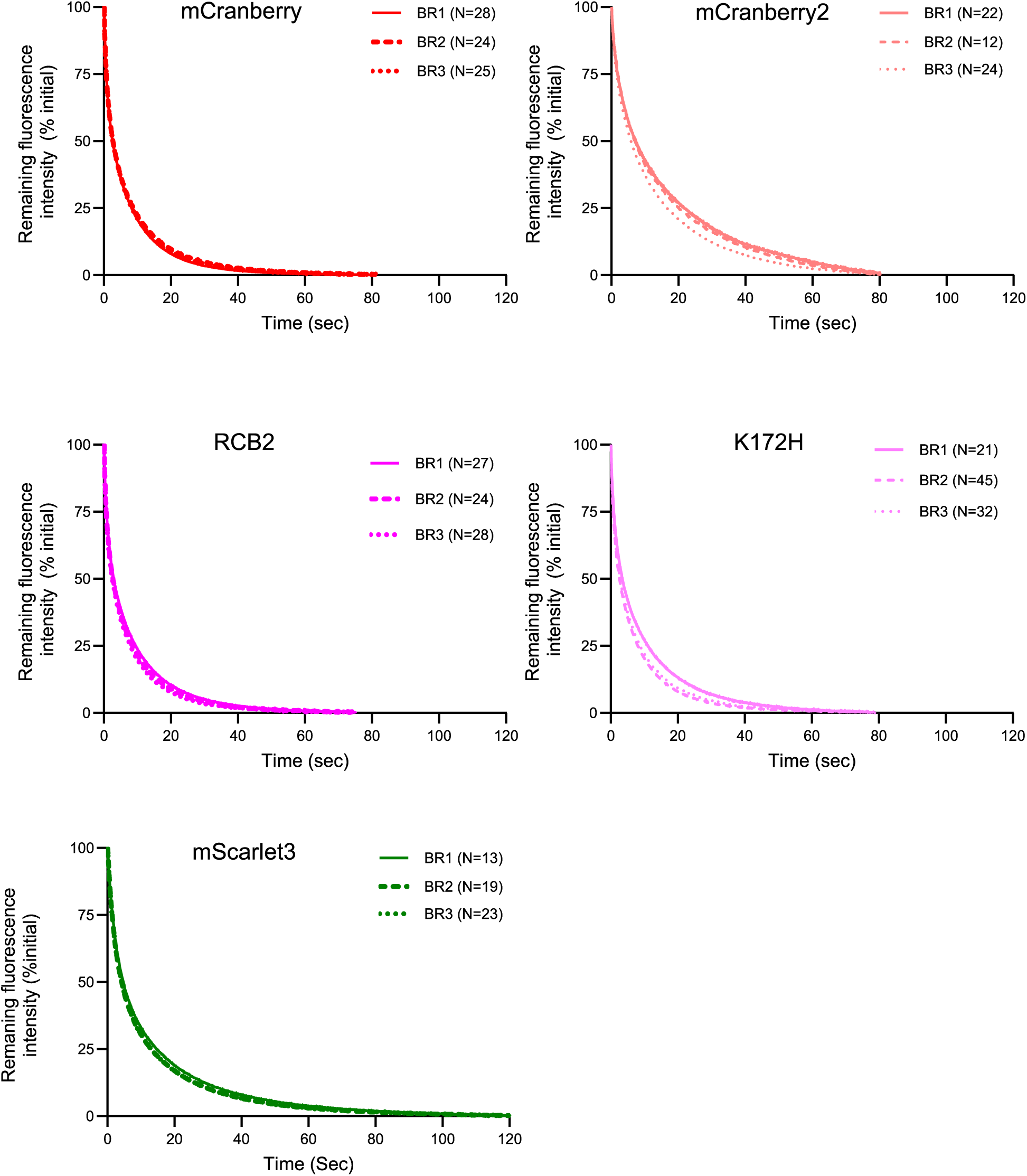
Photobleaching decay in U2OS cell. The fluorescence decay curves of the red fluorescent proteins across three independent biological replicates (BR), with the total number of analyzed cells (*N*) indicated for each replicate. As described in the Methods, the time axis was corrected to account for differences in extinction coefficient, fluorescence quantum yield, and excitation efficiency.

### 7. Brightness in Mammalian Cells

**Extended Data Fig. 7.**
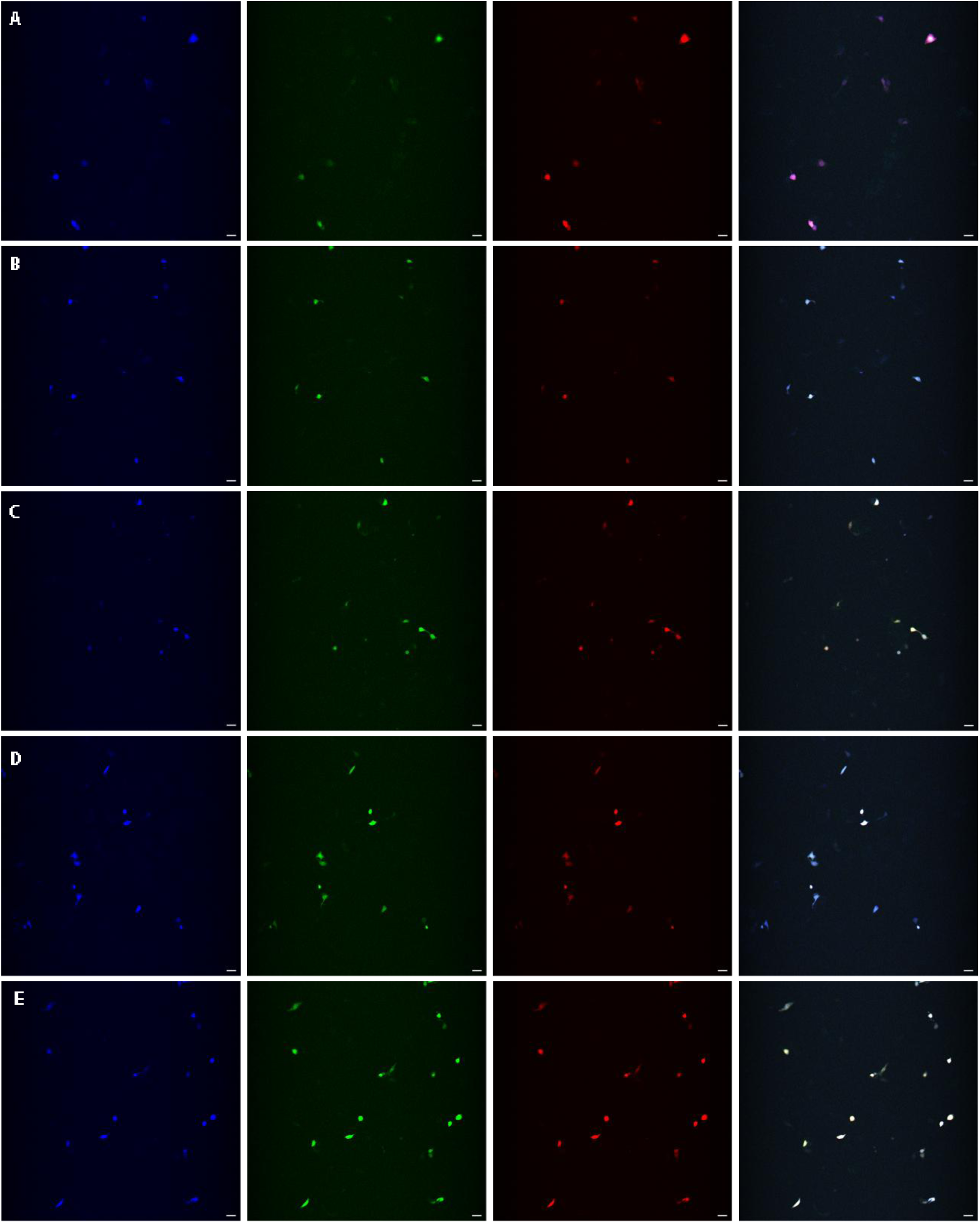
Cellular images of cells expressing mTurquise2 and RFPs. Representative confocal images of selected U2OS cells 48 hours post-transfection with pDRESS_mTurquoise2_spatial-linker-P2A_RFP. Columns from left to right display images acquired using distinct excitation and emission settings: first column (402 nm excitation, 425-475 nm emission), second column (487 nm excitation, 500-550 nm emission), third column (561 nm excitation, 570-620 nm emission), and fourth column represents merged of all channels. Rows represent different variants: (A) mScarlet3, (B) mCranberry, (C) mCranberry2, (D) RCB2, and (E) K172H. Scale bar: 50 μm. Fluorescence intensity is displayed with uniform brightness and contrast adjustments across all channels.

### 8. Red/Cyan ratio in mammalian cells

**Extended Data Fig. 8.**
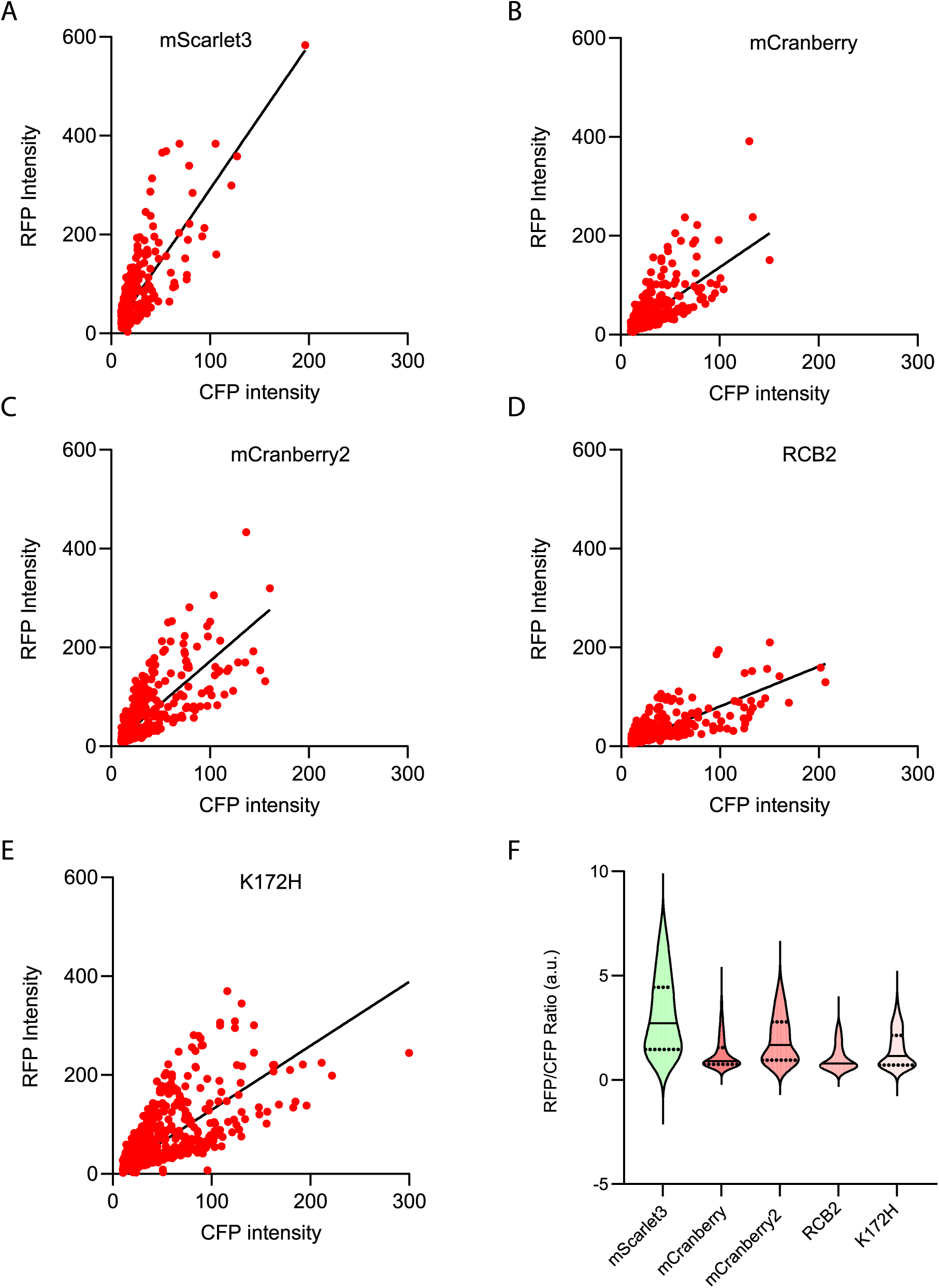
Red/ Cyan ratio in mammalian cells. **A-E,** Fluorescence intensity across the red and cyan channels was measured 48 h post-transfection. The red-to-cyan ratio and its distribution for each mutant are shown. Red fluorescence values were corrected for the spectral throughput of the microscope as described in the main text. Relative brightnesses of the RFPs (compared to mScarlet3) were calculated using the slopes of linear regression lines forced through the origin, with calculations provided in Source Data (Extended Data Fig. 7). **F,** Violin plots display the distribution of the red-to-cyan ratio across all cells analyzed, where solid black lines represent the median.

### 9. GFP Dead End Chromophore formations

**Extended Data Fig. 9.**
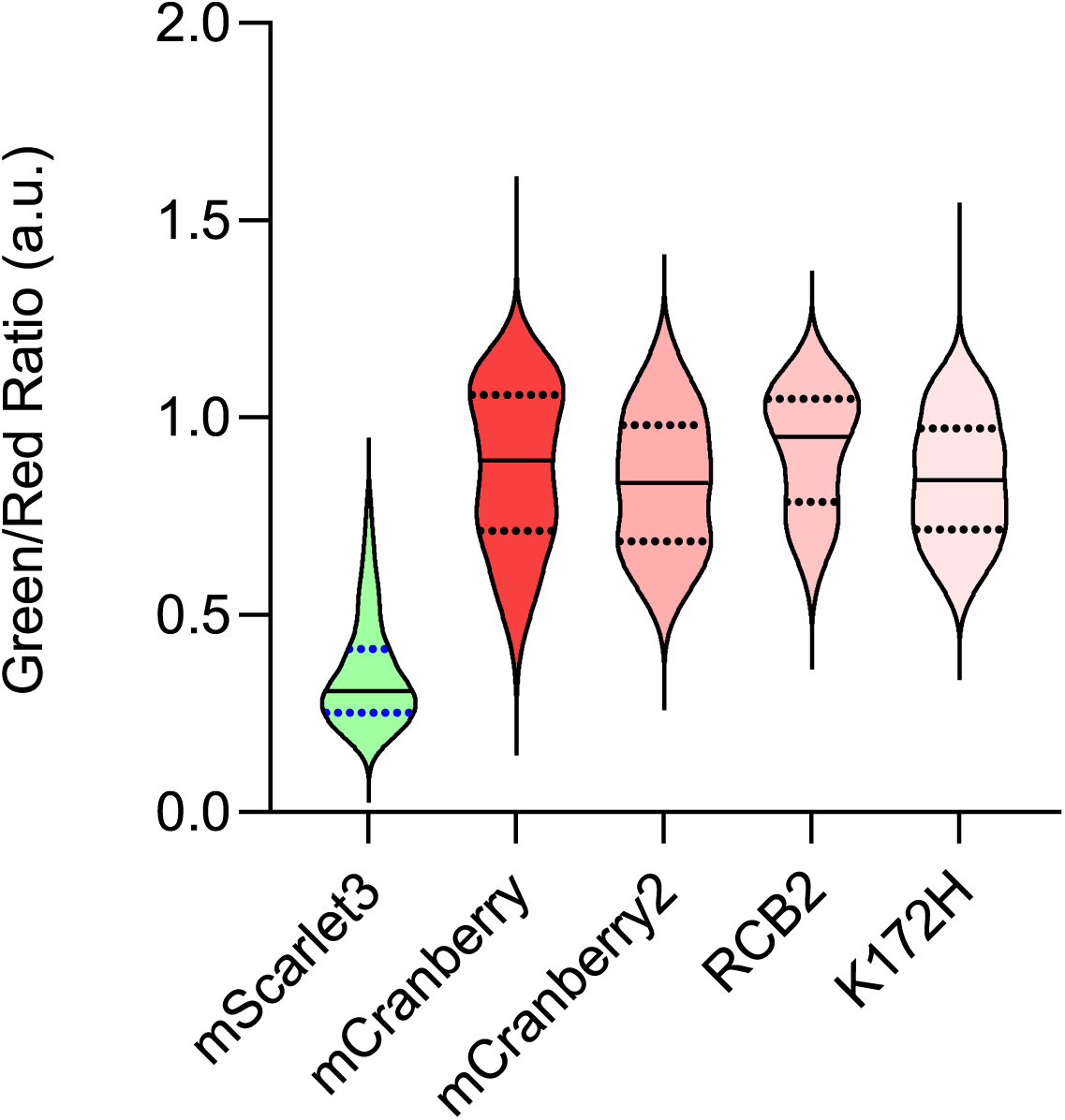
Ratiometric assessment of green dead-end chromophore formation. Green and red channel fluorescence intensities were recorded 48 hours post-transfection. The cellular distribution of green-to-red ratios is presented as violin plots (medians indicated by solid black lines). Higher green-to-red ratios reflect a greater proportion of dead-end green chromophore side-products. Compared to mScarlet3, the evaluated mCranberry mutants demonstrate significantly higher fractions of the green chromophore.

### 10. Cell viability assessment of CytERM-RFP

**Extended Data Fig. 10.**
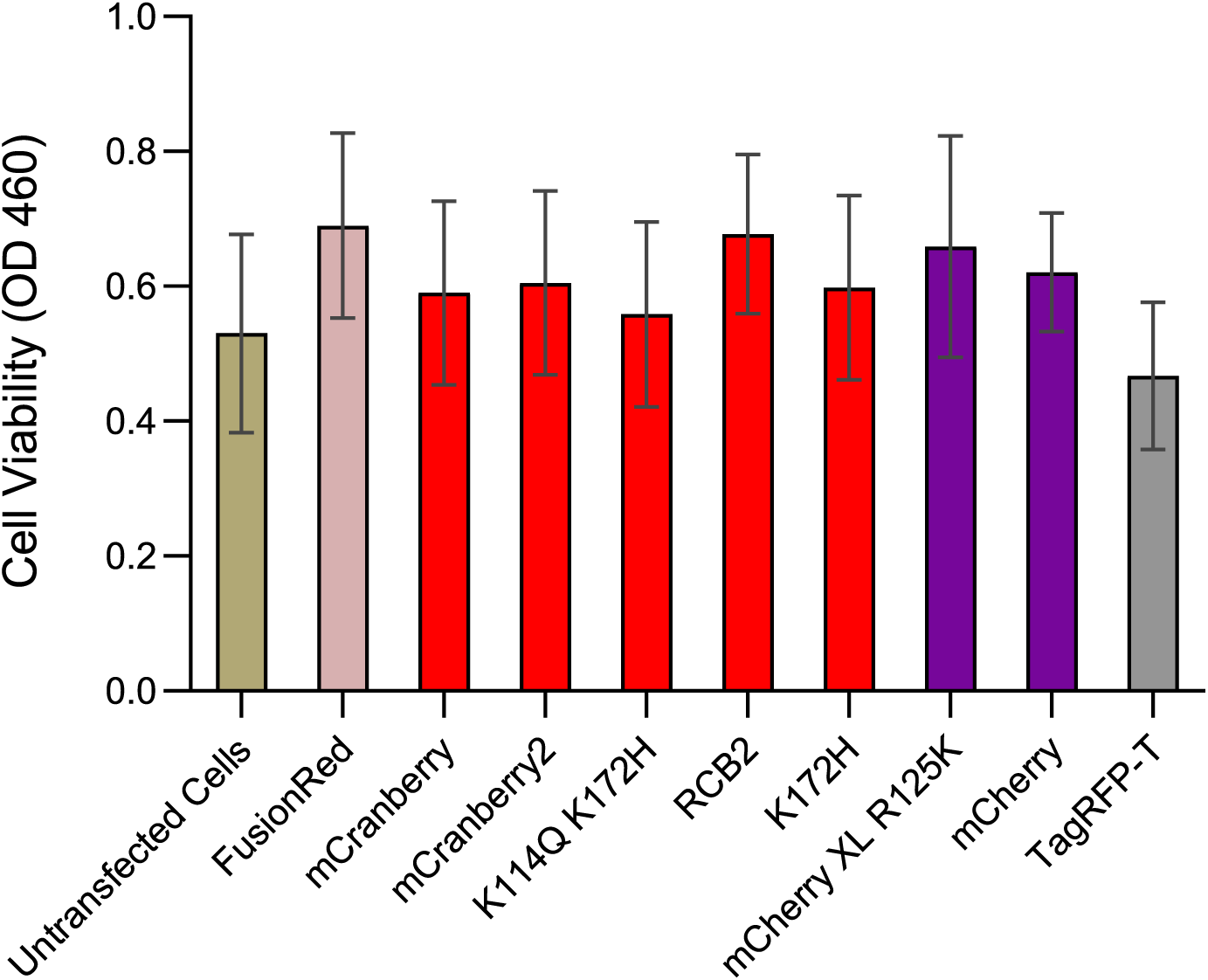
Evaluation of cellular viability following expression of CytERM-RFP fusion constructs for OSER analysis. Cellular cytotoxicity following 24-hour transfection of U2OS cells was assessed by measuring the reduction of WST-8 to water-soluble formazan, with absorbance read at 460 nm. Results are reported as mean absorbance at 460 nm (± s.d). across six independent transfections.

### 11. Ion Exchange Chromatography

**Extended Data Fig. 11.**
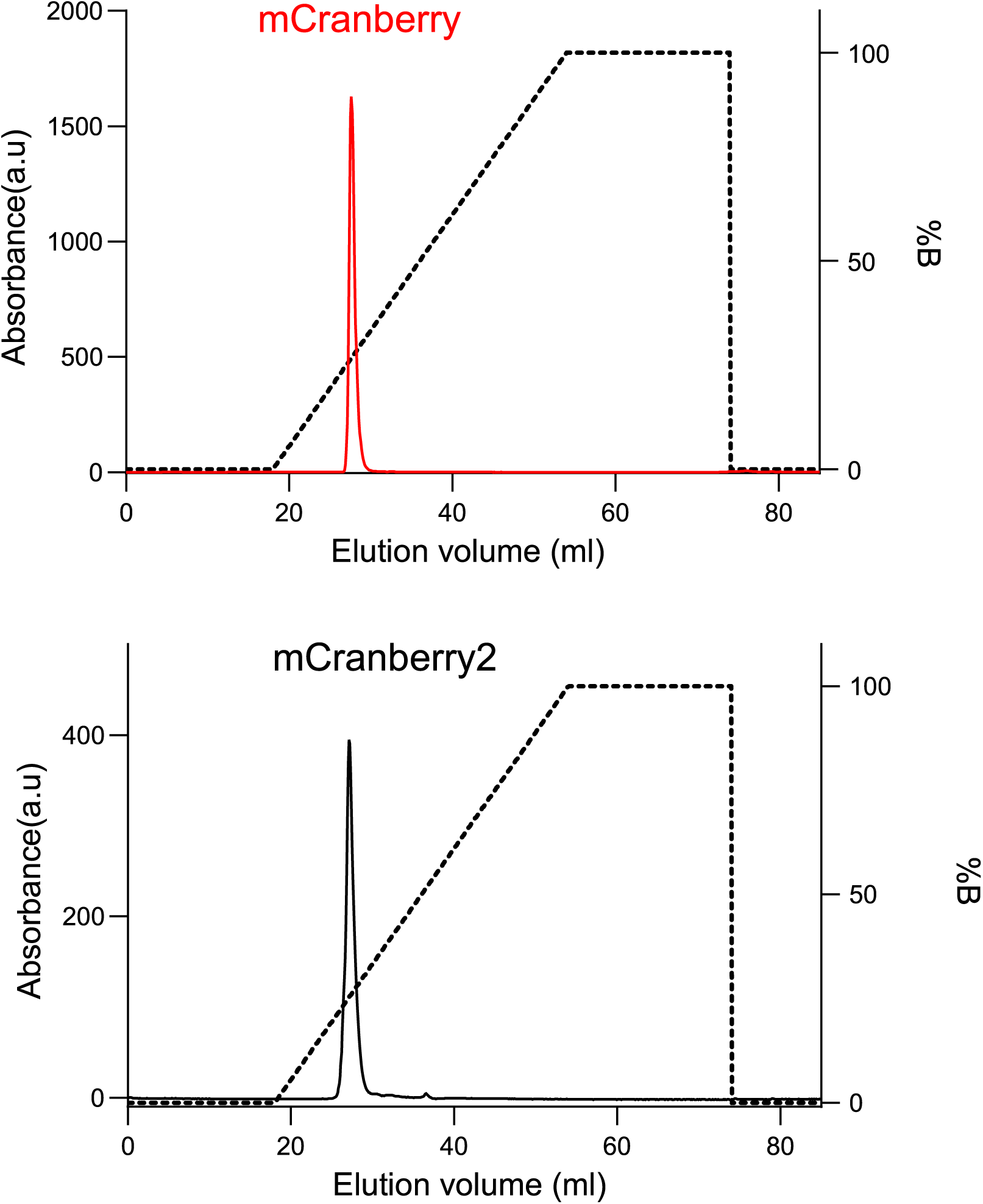
Ion-exchange chromatography of mCranberry mutants. Following His-tag removal, protein samples were applied to a Resource Q column on an FPLC system prior to crystallization screening. Elution was monitored by absorbance at 280 nm using a linear gradient (dashed line) of Buffer B (20 mM Tris-HCl, 1 M NaCl, pH 7.4).

