## Supplementary Material for "mCranberry: A rationally engineered monomeric red fluorescent protein with 85% quantum yield"

#### Table of contents

| **Tables** | **Content** |
|---|---|
| Table 1 | Spectroscopic properties of purified RFP and its variants. |
| Table 2 | Absorption, Excitation and Emission Spectra for the RFPs and Mutants |
| Table 3 | mCranberry and variants thereof |
| Table 4 | Protein Sequence IDs |
| Table 5 | DNA Sequence IDs |
| Table 6 | Crystallographic Data Summary |
| Table 7 | Hydrogen bond occupancies |
| Table 8 | Bridging hydrogen bond occupancies |
| Table 9 | All chromophore hydrogen bond occupancies |

| **Section** | **Title** |
|---|---|
| Sec. 1 | In Silico Mutagenesis |
| Sec. 2 | Program for OSER Analysis |
| Sec. 3 | Schematic of time-correlated single-photon counting setup |
| Sec. 4 | Protein sequence alignment |
| Sec. 5 | Computed per-residue RMSF |

#### Sec. 1. In Silico Mutagenesis

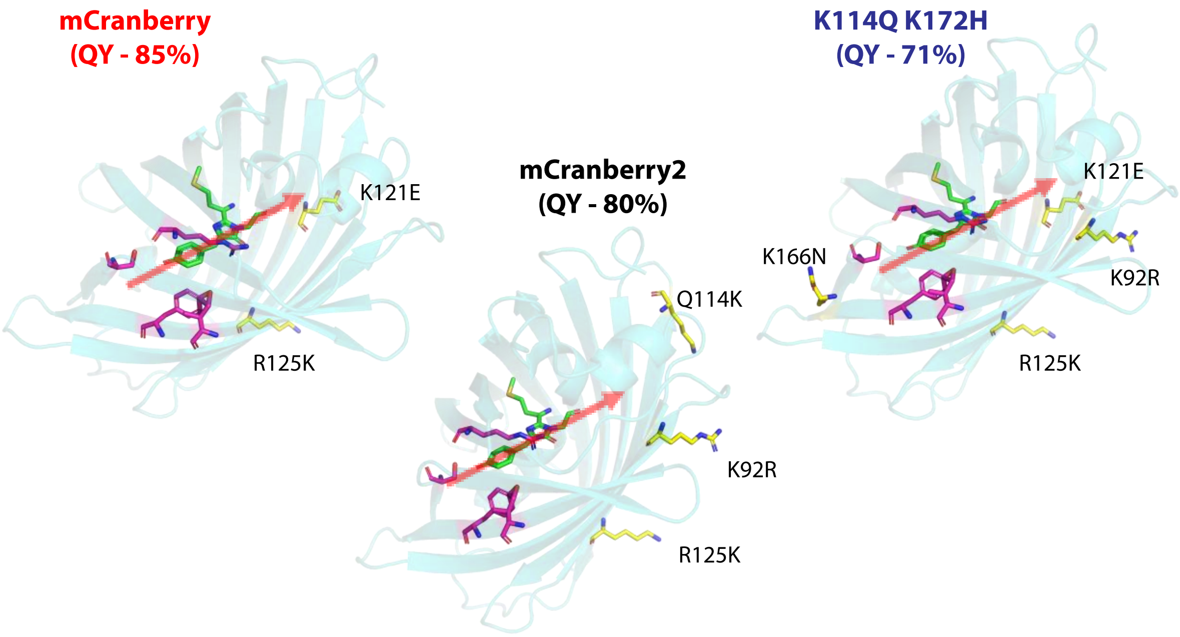

**Fig S1:** ***In silico*** **mutagenesis of the mCherry structure (PDB: 2H5Q)**[1] was performed using COOT (Crystallographic Object-Oriented Toolkit) to model the different mCranberry variants.[2] Surface and internal amino acid substitutions are highlighted in yellow and purple, respectively. Mutations are relative to mCherry and residue numbering follows PDB:2H5Q. The red arrow at the chromophore plane indicates the predicted orientation of the transition dipole moment.

#### Sec. 2. Program for OSER Analysis

To quantify the OSER score, we developed an automated image analysis pipeline using CellProfiler[3], adapted from a previously described protocol[4] with modifications (Supplementary Fig. S2A). The CellProfiler parameters used are shown in Supplementary Fig. S2B. In agreement with earlier reports[5], karmellae structures (nuclear-associated paired membranes induced by enzyme overexpression) were observed. These structures were excluded from whorl quantification within the analysis pipeline.

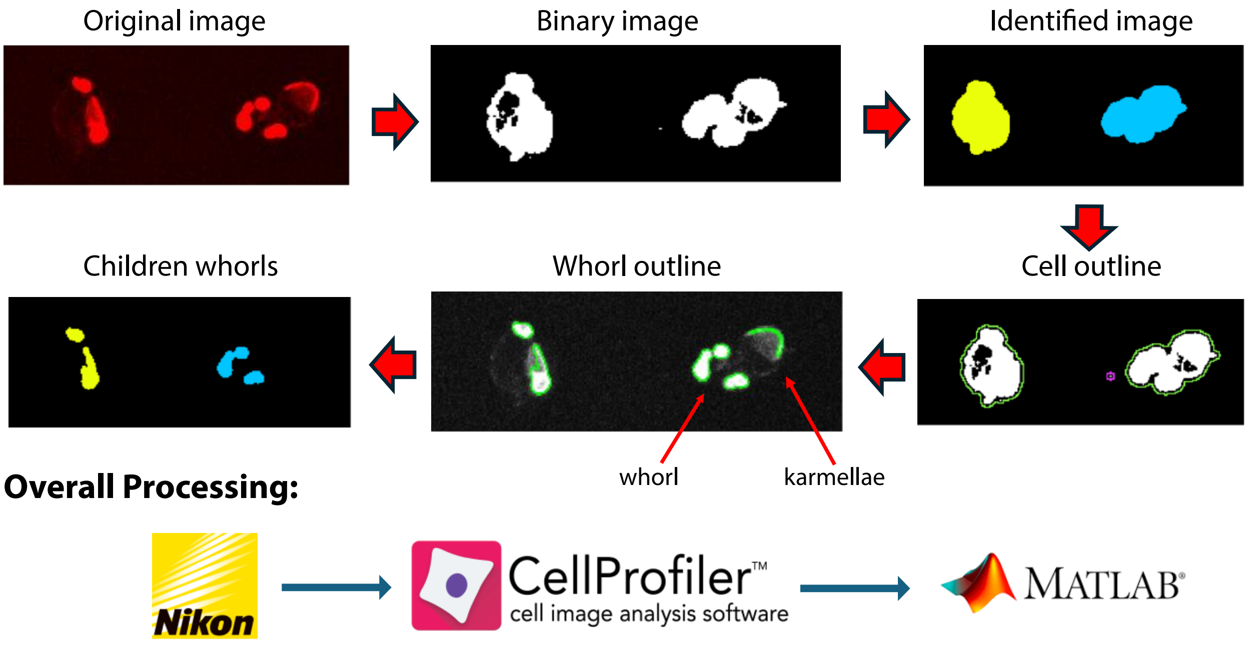

**Fig S2A: High-throughput OSER quantification**. Quantitative analysis of OSER structures in U2OS cells was performed using CellProfiler. Representative images depict TagRFP-T expressing U2OS cells forming whorl structures and karmellae; karmellae were excluded from quantification using the analysis pipeline. Specific parameter settings are detailed below in Supplementary Fig. S2B.

#### Module 1: Threshold

| Setting | Value |
|---|---|
| Select the input image | OSER (from NamesAndTypes) |
| Name the output image | ThreshCell |
| Threshold strategy | Global |
| Thresholding method | Minimum Cross-Entropy |
| Threshold smoothing scale | 1.3488 |
| Threshold correction factor | .4 |
| Lower and upper bounds on threshold | .008 / 1.0 |
| Log transform before thresholding? | ○ Yes ● No |

#### Module 2: IdentifyPrimaryObjects

| Setting | Value |
|---|---|
| Use advanced settings? | ● Yes ○ No |
| Select the input image | ThreshCell (from Threshold #05) |
| Name the primary objects to be identified | cell |
| Typical diameter of objects, in pixel units (Min,Max) | 45 / 225 |
| Discard objects outside the diameter range? | ● Yes ○ No |
| Discard objects touching the border of the image? | ○ Yes ● No |
| Threshold strategy | Global |
| Thresholding method | Otsu |
| Two-class or three-class thresholding? | Two classes |
| Threshold smoothing scale | 1.3488 |
| Threshold correction factor | 1.0 |
| Lower and upper bounds on threshold | 0.0 / 1.0 |
| Log transform before thresholding? | ○ Yes ● No |
| Method to distinguish clumped objects | Intensity |
| Method to draw dividing lines between clumped objects | Shape |
| Automatically calculate size of smoothing filter for declumping? | ● Yes ○ No |
| Automatically calculate minimum allowed distance betw | ● Yes ○ No |
| Speed up by using lower-resolution image to find local maxima? | ● Yes ○ No |
| Display accepted local maxima? | ○ Yes ● No |
| Fill holes in identified objects? | After both thresholding and declumping |
| Handling of objects if excessive number of objects identified | Continue |

#### Module 3: MeasureImageIntensity

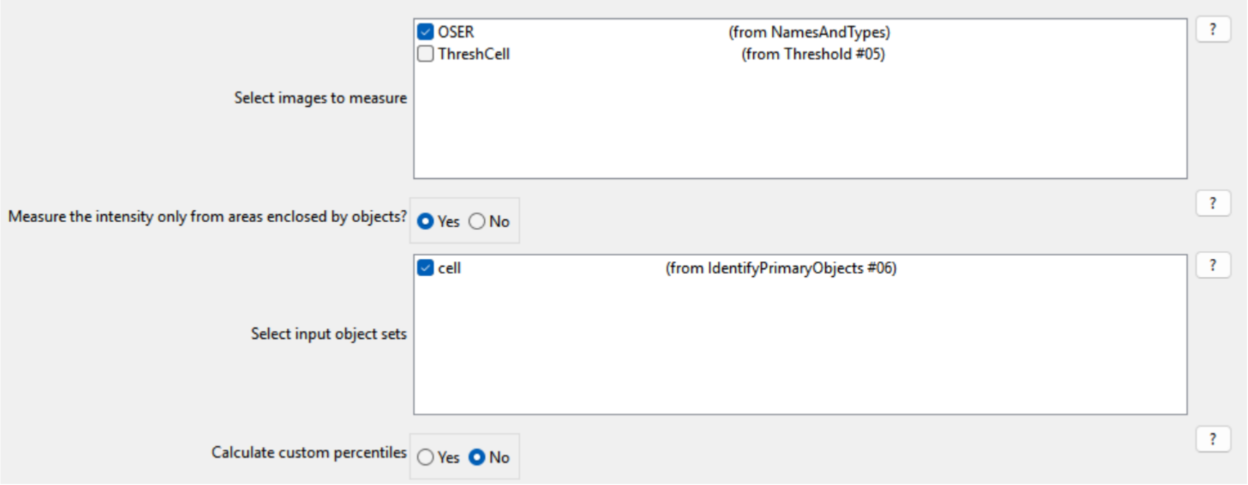

#### Module 4: CalculateMath

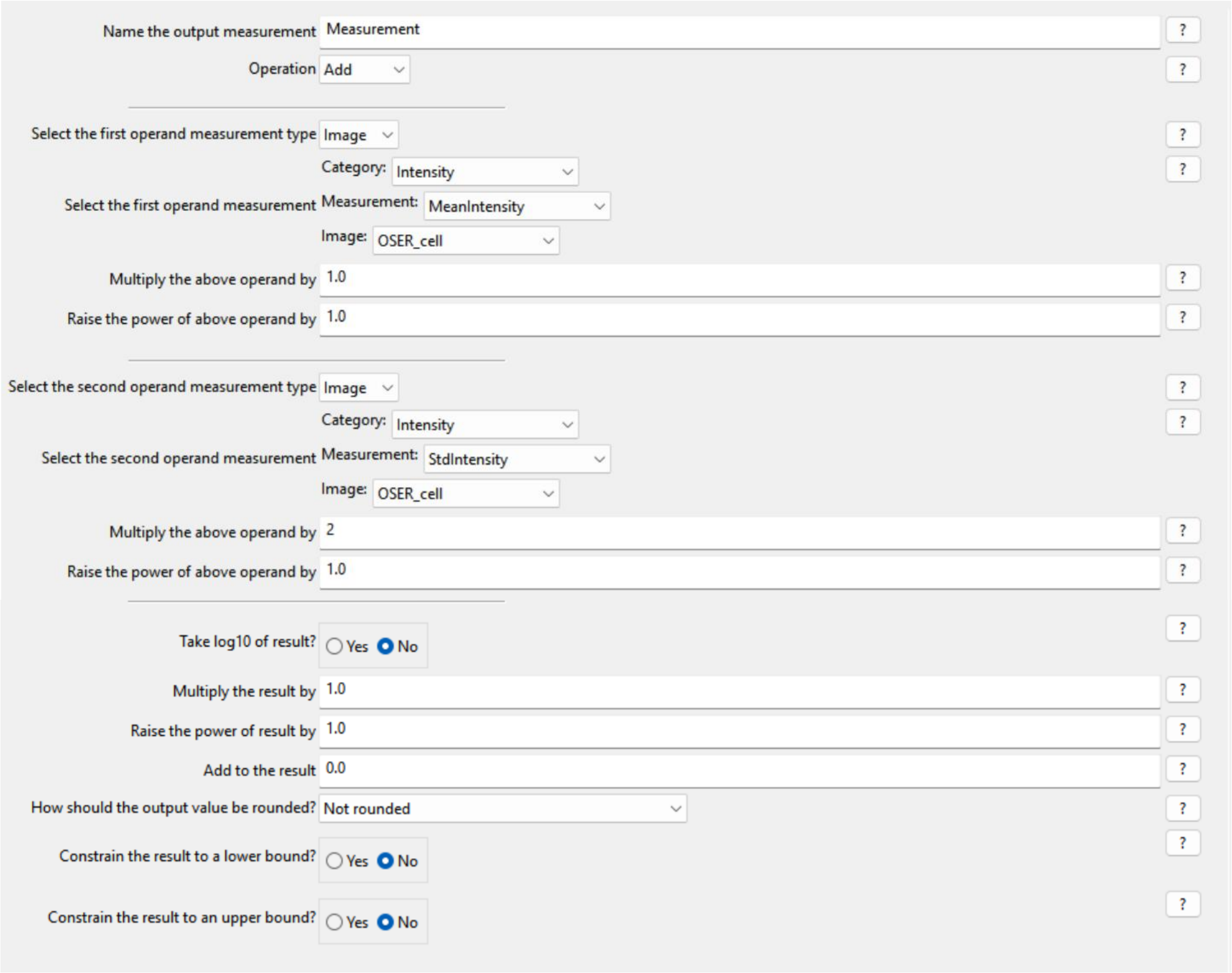

#### Module 5: Threshold

| Setting | Value |
|---|---|
| Select the input image | OSER (from NamesAndTypes) |
| Name the output image | ThreshWhorl |
| Threshold strategy | Global |
| Thresholding method | Otsu |
| Two-class or three-class thresholding? | Two classes |
| Threshold smoothing scale | 0 |
| Threshold correction factor | 1.5 |
| Lower and upper bounds on threshold | 0.005 / 1.0 |
| Log transform before thresholding? | Yes / **No** (selected) |

#### Module 6: IdentifyPrimaryObjects

| Setting | Value |
|---|---|
| Use advanced settings? | **Yes** (selected) / No |
| Select the input image | ThreshWhorl (from Threshold #09) |
| Name the primary objects to be identified | Whorl |
| Typical diameter of objects, in pixel units (Min,Max) | 10 / 30 |
| Discard objects outside the diameter range? | **Yes** (selected) / No |
| Discard objects touching the border of the image? | Yes / **No** (selected) |
| Threshold strategy | Global |
| Thresholding method | Otsu |
| Two-class or three-class thresholding? | Two classes |
| Threshold smoothing scale | 1.3488 |
| Threshold correction factor | 1.0 |
| Lower and upper bounds on threshold | 0.0 / 1.0 |
| Log transform before thresholding? | Yes / **No** (selected) |
| Method to draw dividing lines between clumped objects | None |
| Fill holes in identified objects? | After both thresholding and declumping |
| Handling of objects if excessive number of objects identified | Continue |

#### Module 7: MeasureObjectSizeShape

| Setting | Value |
|---|---|
| Select object sets to measure | [x] Whorl (from IdentifyPrimaryObjects #10)<br>[ ] cell (from IdentifyPrimaryObjects #06) |
| Calculate the Zernike features? | Yes / **No** (selected) |
| Calculate the advanced features? | Yes / **No** (selected) |

#### Module 8: FilterObjects

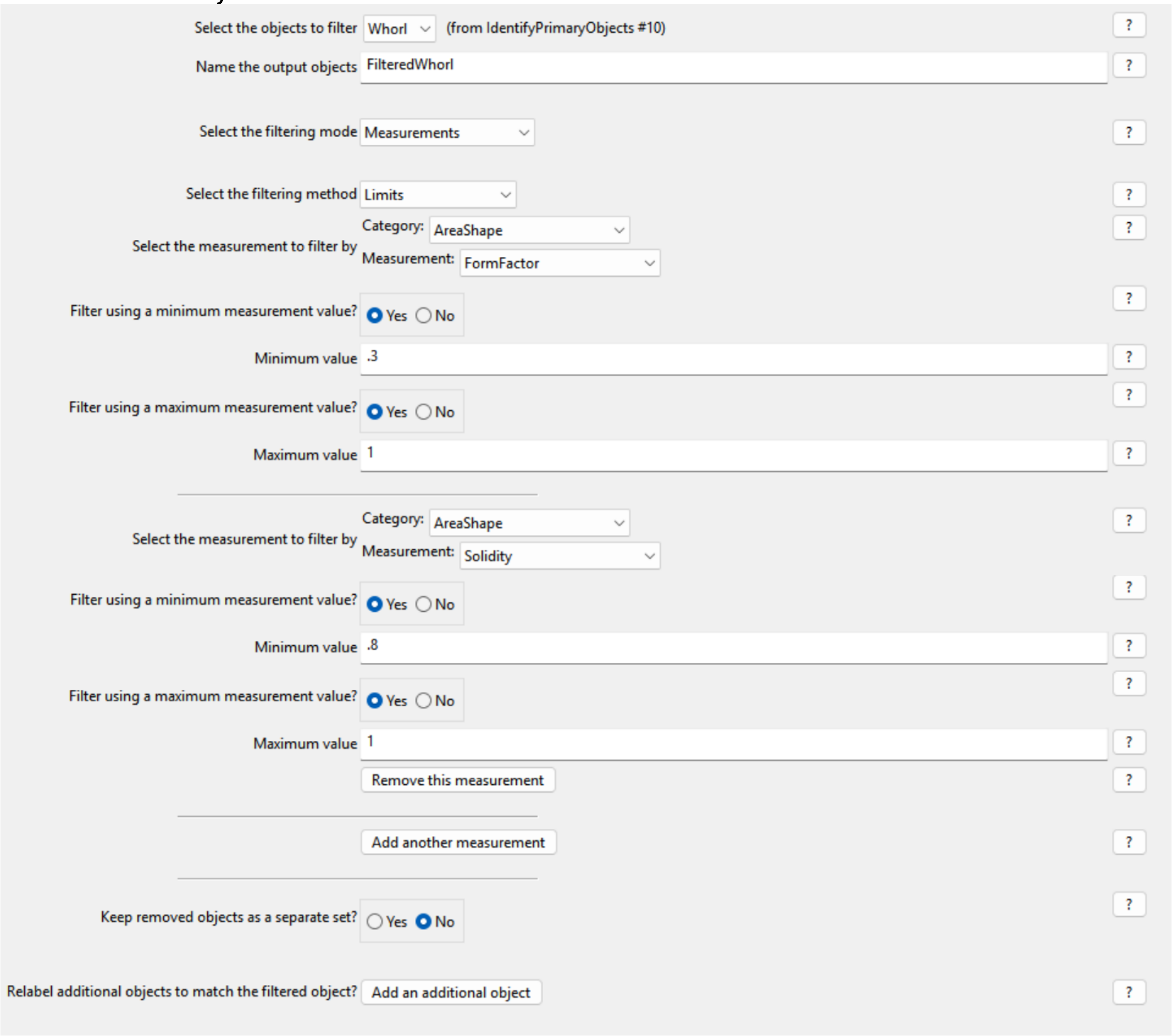

#### Module 9: RelateObjects

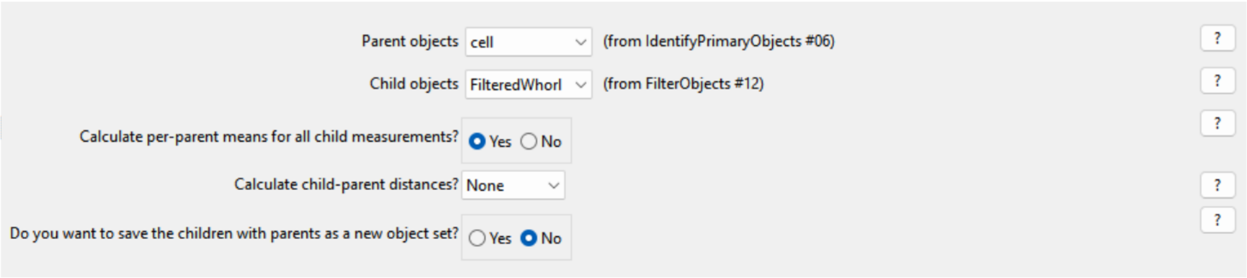

Module 10: MeasureImageIntensity

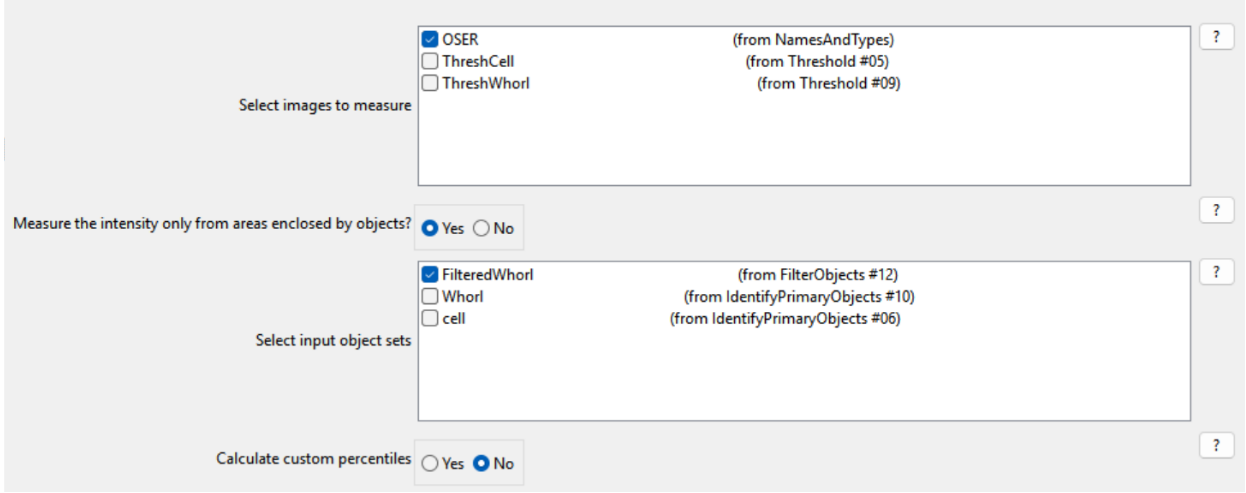

Module 11: ExportToSpreadsheet

**Fig S2B: CellProfiler parameters for OSER quantification**. For confocal acquisitions, images were captured at a scaling resolution of 0.31 µm/px prior to pipeline analysis.

In module 1, raw images were converted into binary images based on pixel intensity to effectively isolate bright cellular structures from the darker background. In module 2, individual cells were identified using specified size constraints (45-225 pixels, i.e. 14-70 µm) and intensity criteria, resulting in distinct cell outlines. In module 3, the fluorescence intensity distribution of each cell identified in the preceding module was measured. In module 4, to quantify heterogeneity of the cell population, we calculated the mean and standard deviation of their intensity distribution. In module 5, whorl structures were identified using a global threshold strategy with a two-class Otsu method. In module 6, structures were filtered by a size criterion of 10-30 pixels (3-9 µm). In module 7, the size and shape of whorl structures are measured. In module 8, karmellae are filtered out and excluded based on FormFactor and Solidity measurements within the AreaShape category, leaving only the whorl structures for final quantification. In module 9, the filtered whorls are related back to their parent cells to determine the number of whorls per cell. Module 10 measures the image intensity of the filtered whorls. Finally, module 11 exports the data to spreadsheets, and MATLAB is used to calculate the number of unique cells containing whorls.

#### Sec. 3. Schematic of time-correlated single-photon counting setup

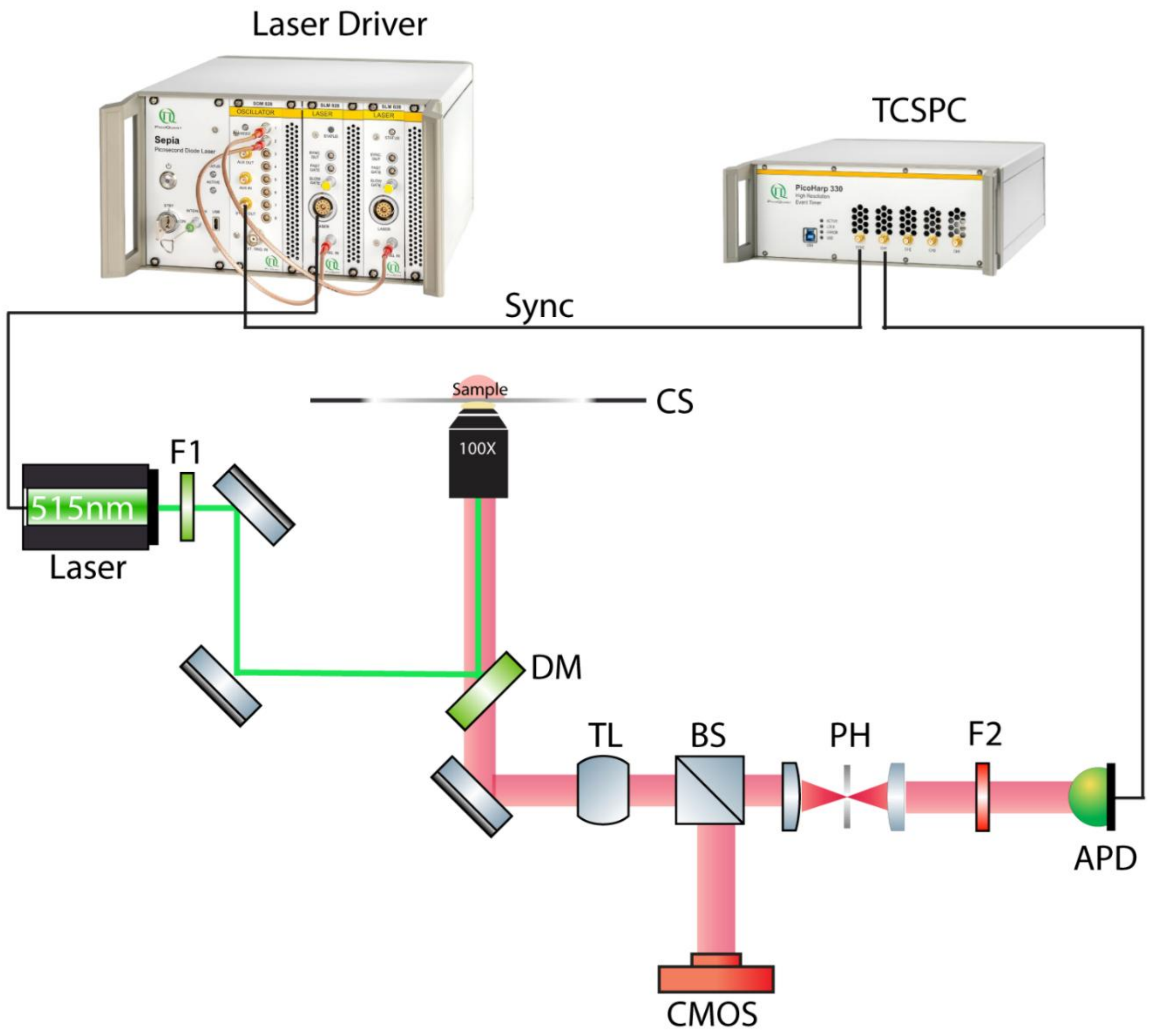

**Fig S3. Schematic of the homebuilt setup**. The sample is excited with a 515 nm solid-state pulsed laser (LDH-D-C-520, PicoQuant, Germany) which is driven by the PDL 828 "Sepia II" driver (PicoQuant, Germany). The laser is filtered with F1(FBH515-10) and directed by two mirrors onto the DM (ZT532/640/NIR_rpc), which reflects the laser to the sample via an objective (Olympus UplanSApo 100X oil immersion objective). The emitted light from the sample passes through the DM and is directed to the detection path with a mirror. The detection path has a beamsplitter (BS) which splits 10% of the light to the CMOS camera (CS165MU, Thorlabs) for precise focusing and the remaining 90% to the APD (SPCM-AQRH-14-TR, Excelitas) via a pinhole (50μm) followed by an emission filter (ET630/75 nm). The laser driver sends the sync-out signal to the TCSPC unit (Picoharp 330, PicoQuant, Germany) for precise time stamps.

#### Sec. 4. Protein sequence alignment

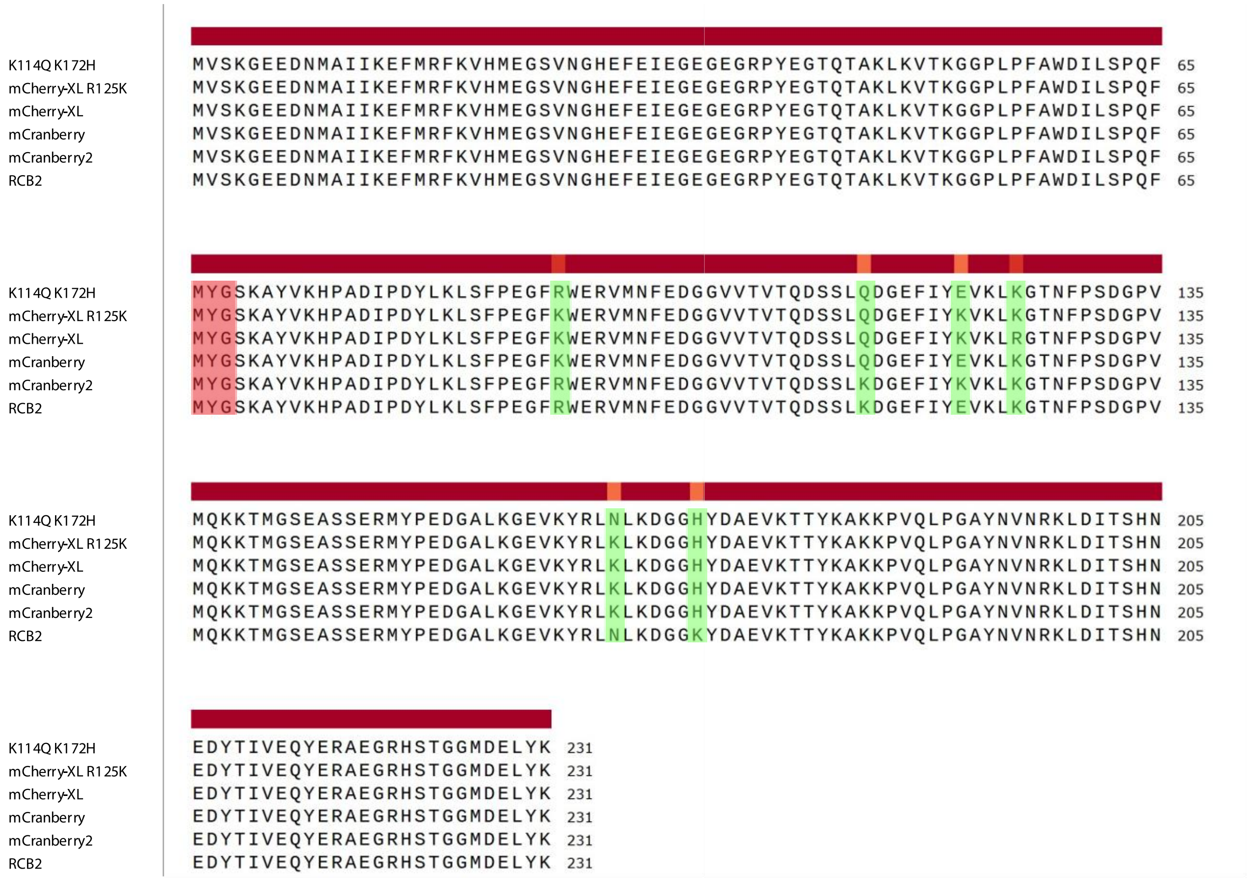

**Fig S4. Sequence alignment of key fluorescent proteins used in this study**. Numbering throughout the main text and supplementary information corresponds to the mCherry crystal structure (PDB: 2H5Q). Residues highlighted in red indicate the chromophore, and those in green mark sites mutated during the development of mCranberry variants.

#### Sec. 5. Computed per-residue RMSF

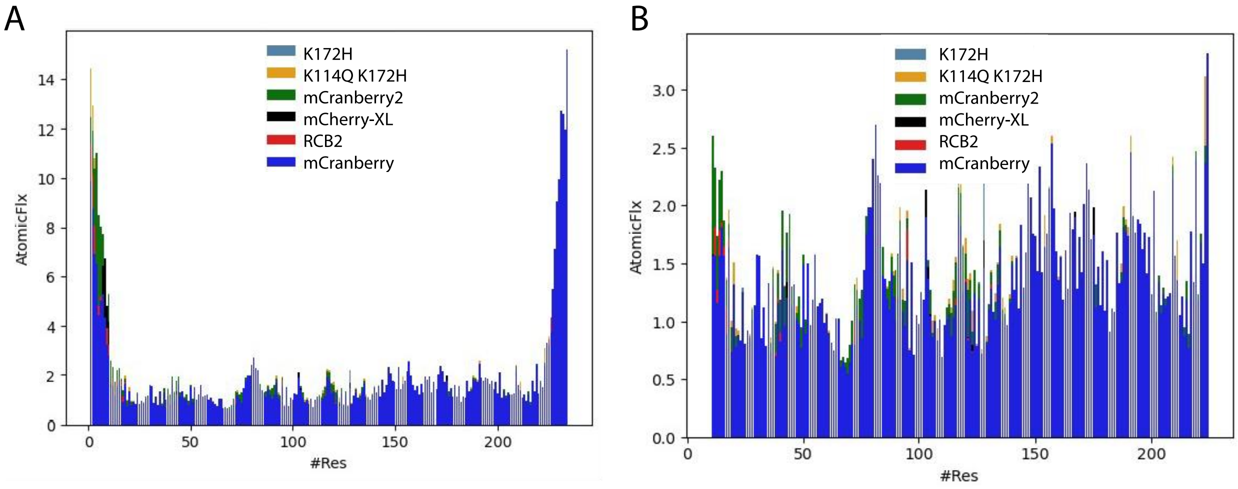

**Fig S5. Computed per-residue RMSF**. **A,** For all systems studied with the wild-type mCherry from previous work included as a comparison and computed per-residue RMSF is shown. **B,** Computed per-residue RMSF without N/C termini. While there are minor differences, particularly in the 120-140 range, most RMSF values for all systems lie below 2.5 Å except for the flexible N/C termini.

#### Table S1: Spectroscopic properties of purified RFP and its variants

| SEQ ID NO | Protein | Abs Max (nm) | Em Max (nm) | EC ($10^3$ $M^{-1}cm^{-1}$) (N) | QY (%) (N) | EC*QY |
|---|---|---|---|---|---|---|
| 1 | mCherry | 586 | 609 | 95.3 ± 3.1 (3) | 22 | 21.0 ± 0.7 |
| 3 | mCherry-XL R125K | 558 | 586 | 56.0 ± 4.2 (3) | 72 ± 5 (3) | 40.3 ± 4.1 |
| 5 | RCB2 | 557 | 585 | 61.4 ± 5.5 (7) | 77 ± 3 (6) | 47.4 ± 4.6 |
| 7 | mCranberry | 556 | 588 | 60.4 ± 1.9 (3) | 85 ± 3 (6) | 51.3 ± 2.0 |
| 9 | mCranberry2 | 558 | 586 | 63.5 ± 1.7 (2) | 80 ± 2 (3) | 50.8 ± 1.9 |
| 11 | K114Q K172H | 558 | 585 | 62.1 ± 1.1 (3) | 71 ± 2 (3) | 44.1 ± 1.5 |
| 13 | R92K K172H | 558 | 586 | 62.5 (1) | 82 (1) | 51.3 |
| 15 | E121K K172H | 558 | 587 | 62.1 ±1.1(3) | 73 (1) | 46.2 |
| 17 | N166K K172H | 556 | 585 | 61.4 ± 3.4 (3) | 75 ± 3 (3) | 46.1 ± 3.1 |
| 19 | R92K K114Q V135L K172H | 556 | 586 | 60.7 (1) | 78 (1) | 47.3 |
| 21 | K114Q E121K K172H | 557 | 586 | 60.9 (1) | 77 (1) | 46.9 |
| 23 | R92K N166K K172H | 556 | 586 | 59.0 (1) | 81 (1) | 47.8 |
| 25 | R92K E121K K172H | 558 | 585 | 63.3 (1) | 79 (1) | 50.0 |
| 27 | K114Q N166K K172H | 556 | 584 | 59.0 (1) | 80 (1) | 47.2 |
| 29 | R92K K114Q E121K K172H | 558 | 586 | 61.6 (1) | 83 ± 2 (3) | 51.1 ± 1.2 |
| 31 | R92K E121K N166K K172H | 557 | 586 | 61.7 (1) | 81 | 50.0 |
| 33 | K114Q E121K N166K K172H | 556 | 586 | 61.9 (1) | 82 ± 1 (3) | 50.8 ± 0.6 |
| 35 | R92K | 558 | 585 | 62.2 ± 4.6 (3) | 80 ± 1 (3) | 49.8 ± 3.7 |
| 37 | K114Q | 557 | 585 | 55.4 ± 5.1(3) | 77 ± 3 (3) | 42.7 ± 4.3 |
| 39 | E121K | 558 | 585 | 61.2 ± 1.9 (3) | 79 ± 2 (3) | 48.3 ± 1.9 |
| 41 | K125R | 558 | 586 | 62.2 ± 8.0 (3) | 80 ± 4 (3) | 49.8 ± 6.9 |
| 43 | N166K | 557 | 586 | 60.1 ± 1.5 (3) | 82 ± 2 (3) | 49.3 ± 1.7 |
| 45 | K172H | 558 | 587 | 59.9 ± 1.5 (3) | 83 ± 1 (3) | 49.7 ± 1.4 |
| 47 | R92K, K114Q, K125R, N166K, K172H | 557 | 584 | 85.0 (1) | x | x |
| 49 | R92K, K114Q, E121D, N166K, K172H | 556 | 558 | 87.0 (1) | x | x |
| 51 | mScarlet | 569 | 590 | 106.2 ± 9.4 (3) | 71 (1) | 75.4 ± 6.7 |
| 53 | mScarlet K122E | 569 | 591 | x | 73.0 (1) | x |
| 55 | mScarlet R126K | 569 | 591 | x | 66.0 (1) | x |
| 57 | mScarlet K122E R126K | 569 | 591 | x | 69 ± 0.02 (3) | x |
| 59 | mScarlet3 | 569 | 590 | 97 (1) | 78.0 (1) | 75.7 |
| 61 | mScarlet3 K121E | 570 | 590 | x | 79.0 (1) | x |
| 63 | mScarlet3 R125K | 569 | 590 | x | 64.0 (1) | x |
| 65 | mScarlet3 K121E R125K | 569 | 591 | x | 74.0 (1) | x |
| 67 | mScarlet3-H | 552 | 587 | x | 19.0 (1) | x |
| 69 | mScarlet3-H K121E | 551 | 588 | x | 21.0 (1) | x |
| 71 | mScarlet3-H R125K | 552 | 588 | x | 22.0 (1) | x |
| 73 | mScarlet3-H K121E R125K | 551 | 588 | x | 22.0 (1) | x |
| 75 | mScarlet3-S2 | 548 | 586 | x | 33.0 (1) | x |

| | | | | | | |
|---|---|---|---|---|---|---|
| 77 | mScarlet3-S2 K121E | 548 | 586 | x | 30.0 (1) | x |
| 79 | mScarlet3-S2 R125K | 548 | 586 | x | 29.0 (1) | x |
| 81 | mScarlet3-S2 K121E R125K | 547 | 586 | x | 34.0 (1) | x |
| | FusionRed | 575 | 596 | 85.8 ± 5.9(3) | 24 ± 2(3) | 20.6 ± 2.2 |
| | TagRFP-T | 556 | 586 | 85.4 ± 3.9 (3) | 50 ± 2 (3) | 42.7 ± 2.6 |

N= Number of biological replicates

#### Table S2: Absorption, Excitation and Emission Spectra for the RFPs and Mutants

**SEQ ID NO: 1**

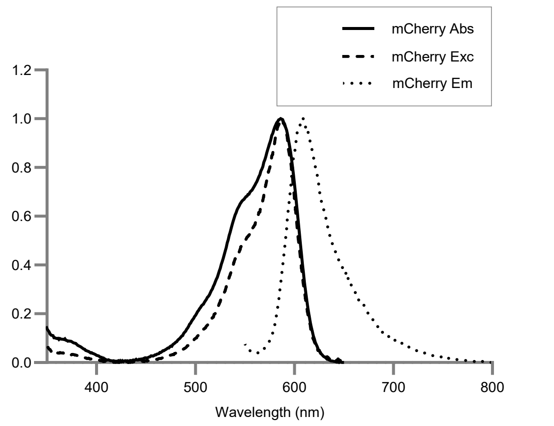

**SEQ ID NO: 3**

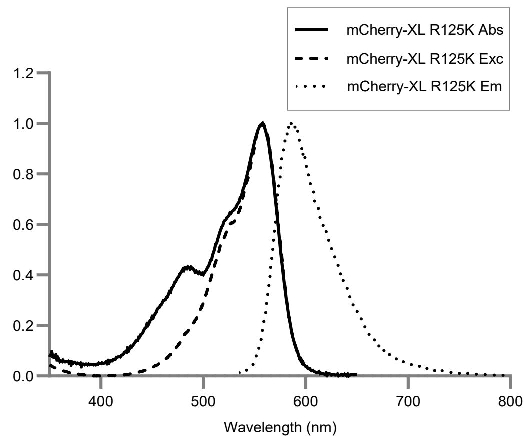

**SEQ ID NO: 5**

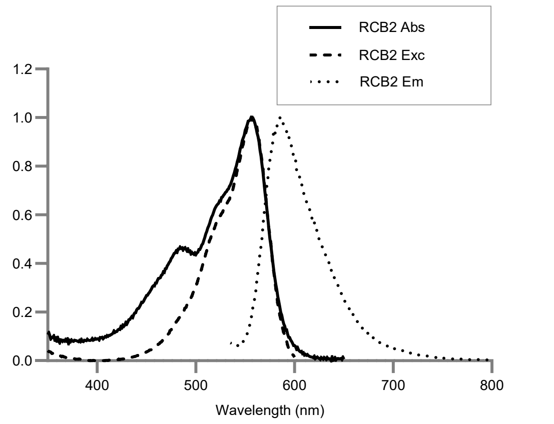

**SEQ ID NO: 7**

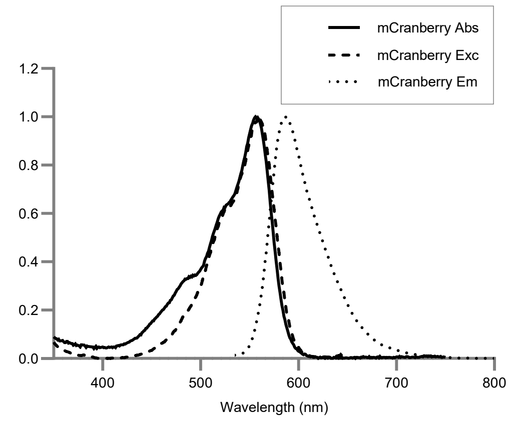

**SEQ ID NO: 9**

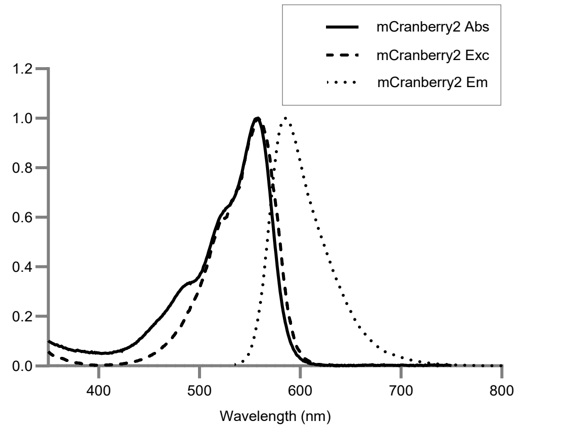

**SEQ ID NO: 11**

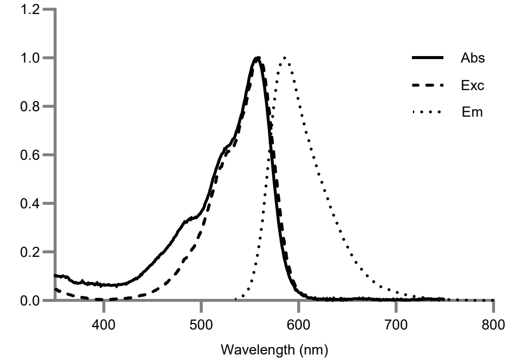

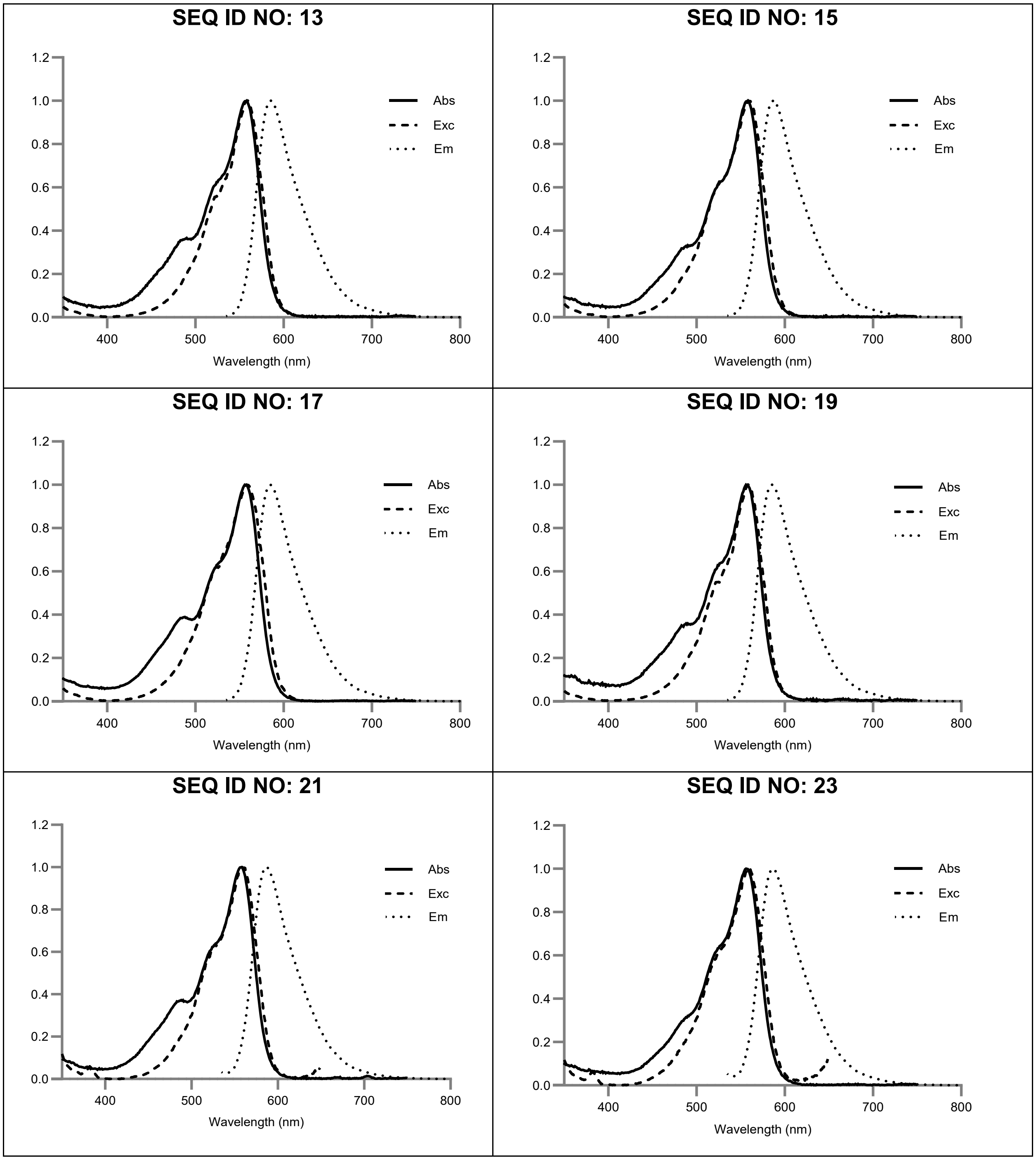
SEQ ID NO: 13
SEQ ID NO: 15
SEQ ID NO: 17
SEQ ID NO: 19
SEQ ID NO: 21
SEQ ID NO: 23
Abs
Exc
Em
1.2
1.0
0.8
0.6
0.4
0.2
0.0
400
500
600
700
800
Wavelength (nm)

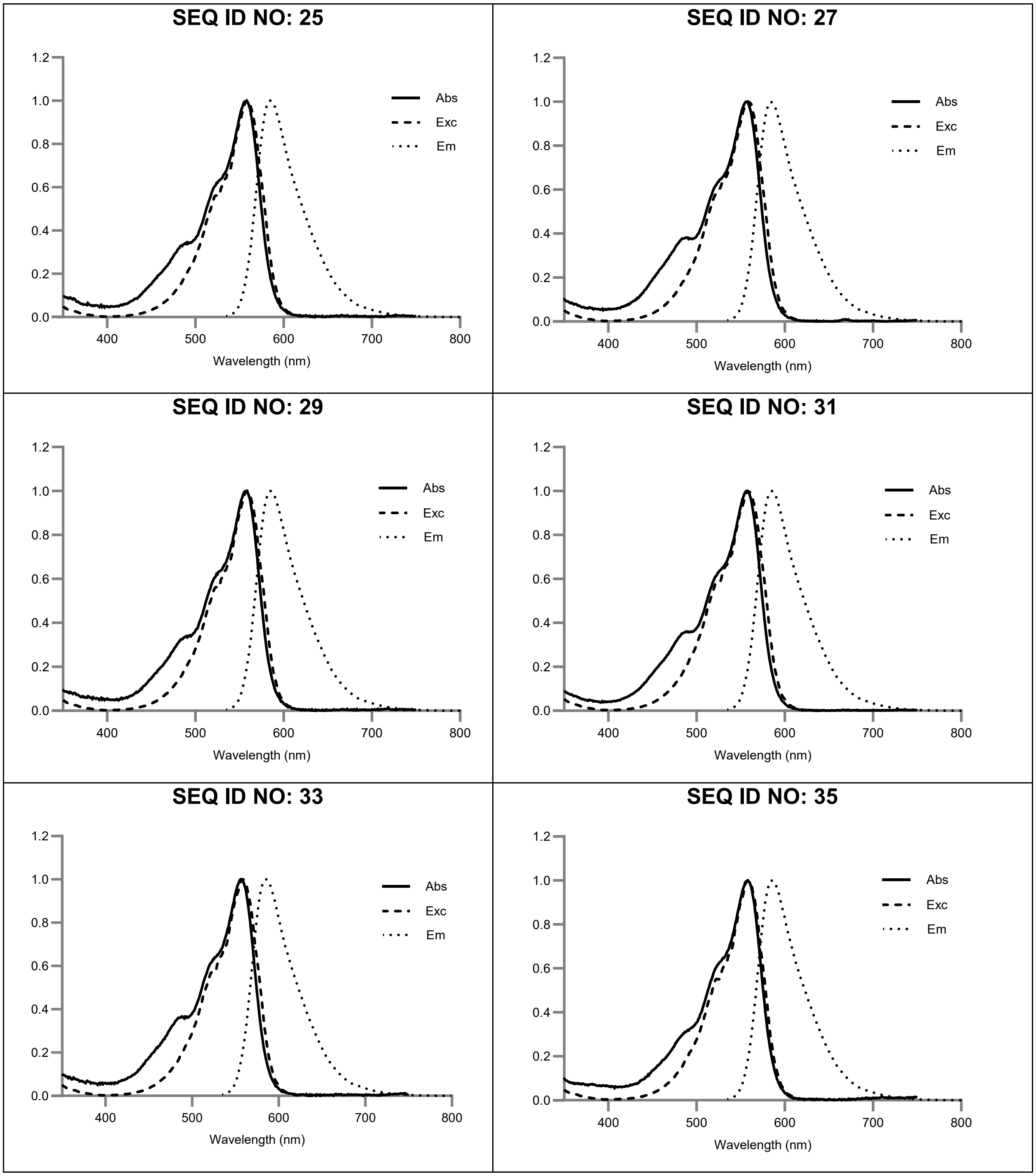
SEQ ID NO: 25
SEQ ID NO: 27
SEQ ID NO: 29
SEQ ID NO: 31
SEQ ID NO: 33
SEQ ID NO: 35
Abs
Exc
Em
Wavelength (nm)
0.0
0.2
0.4
0.6
0.8
1.0
1.2
400
500
600
700
800

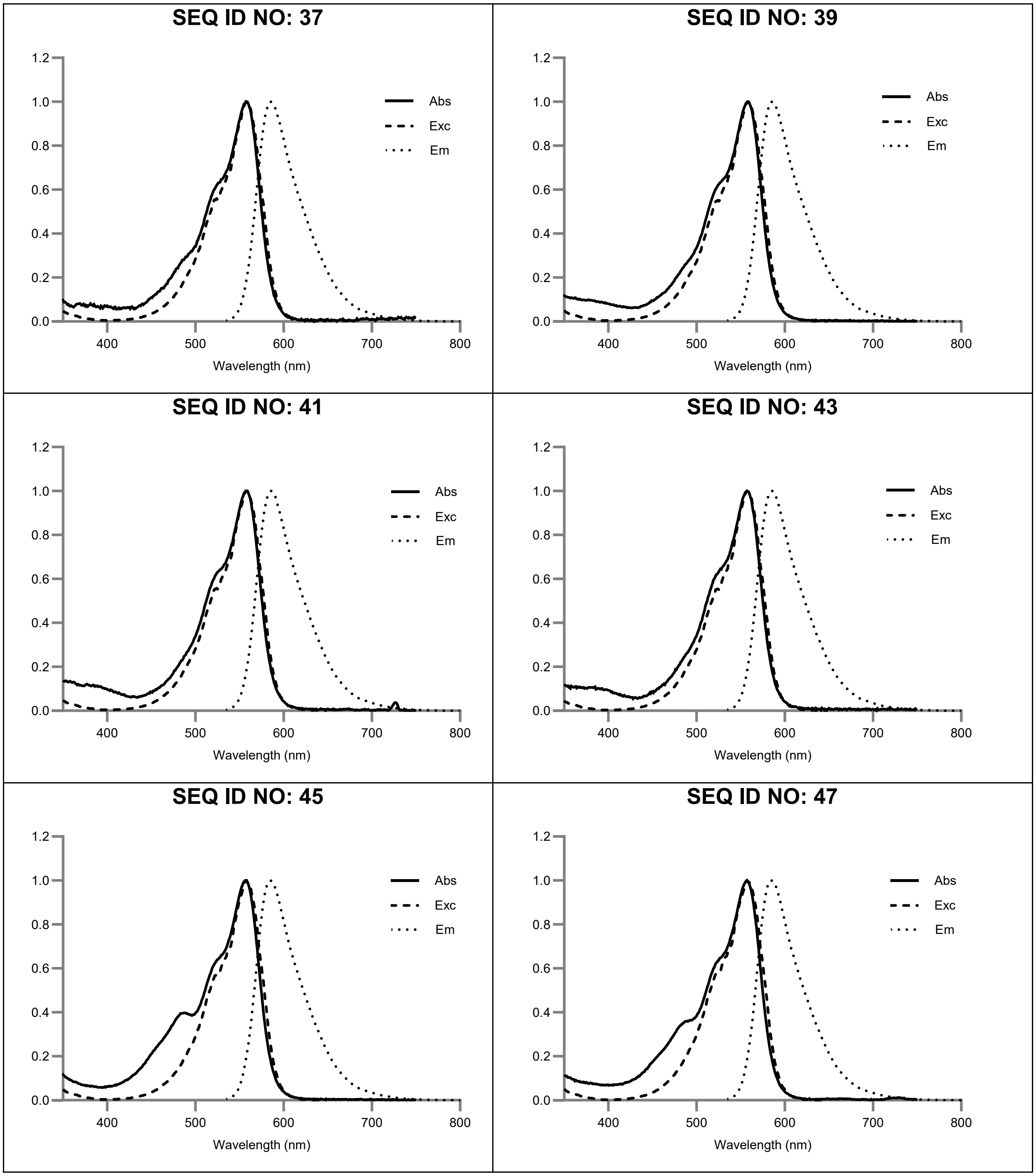
SEQ ID NO: 37
SEQ ID NO: 39
SEQ ID NO: 41
SEQ ID NO: 43
SEQ ID NO: 45
SEQ ID NO: 47
Abs
Exc
Em
1.2
1.0
0.8
0.6
0.4
0.2
0.0
400
500
600
700
800
Wavelength (nm)

**SEQ ID NO: 49**

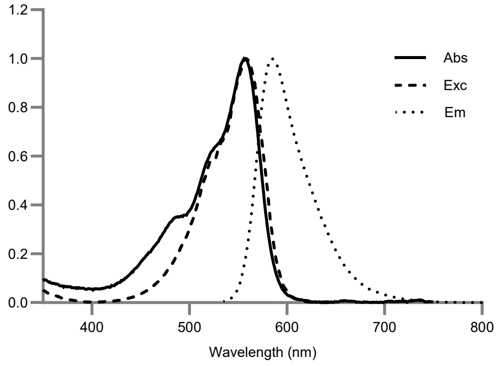

**SEQ ID NO: 51**

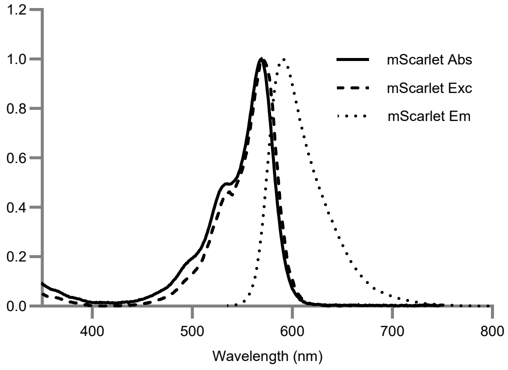

**SEQ ID NO: 53**

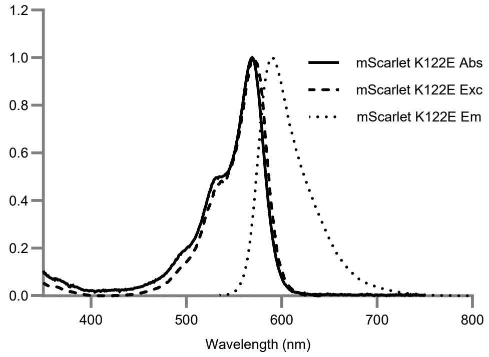

**SEQ ID NO: 55**

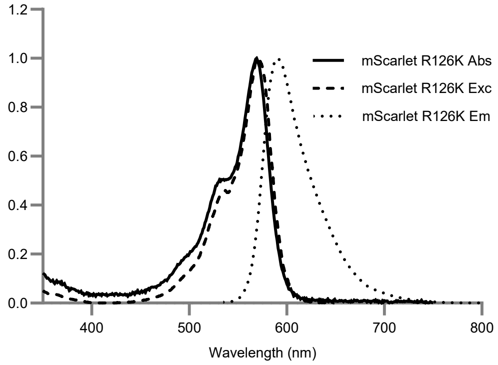

**SEQ ID NO: 57**

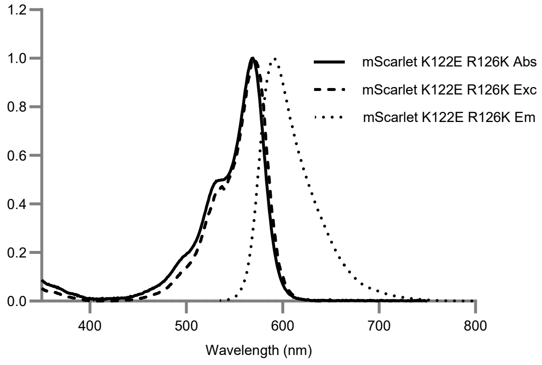

**SEQ ID NO: 59**

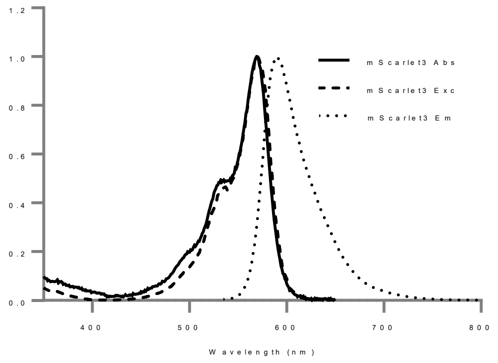

SEQ ID NO: 61

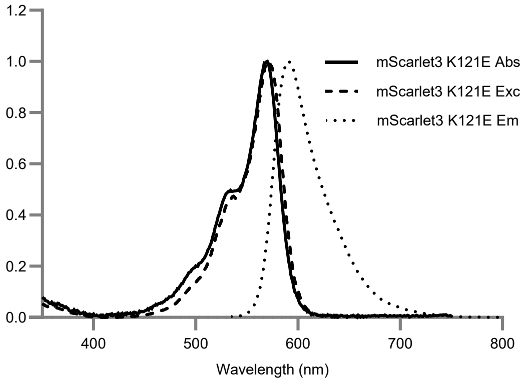

SEQ ID NO: 63

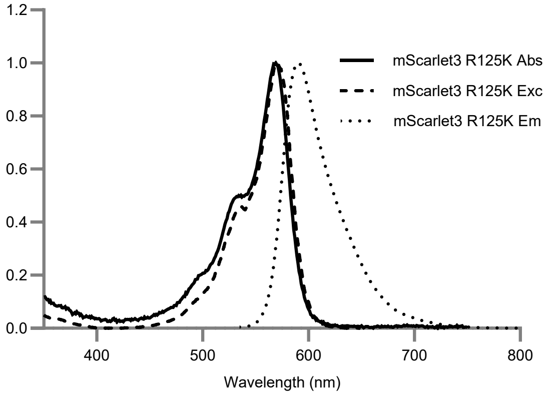

SEQ ID NO: 65

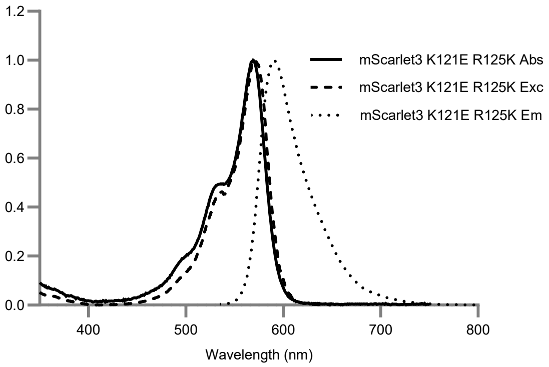

SEQ ID NO: 67

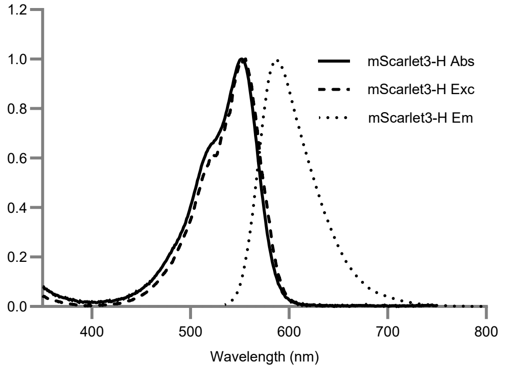

SEQ ID NO: 69

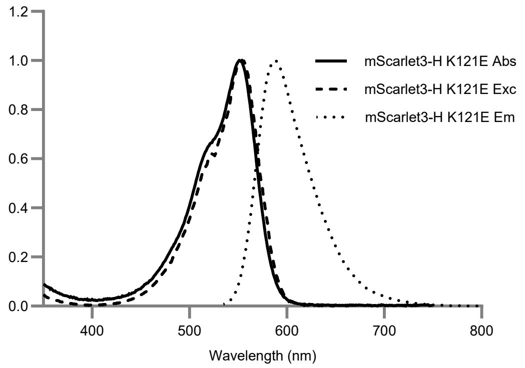

SEQ ID NO: 71

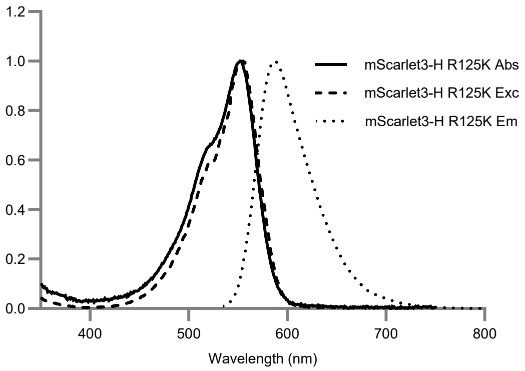

**SEQ ID NO: 73**

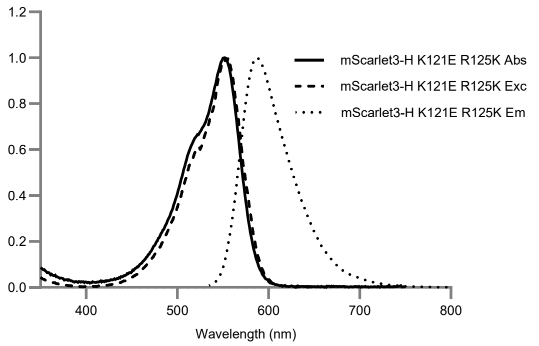

**SEQ ID NO: 75**

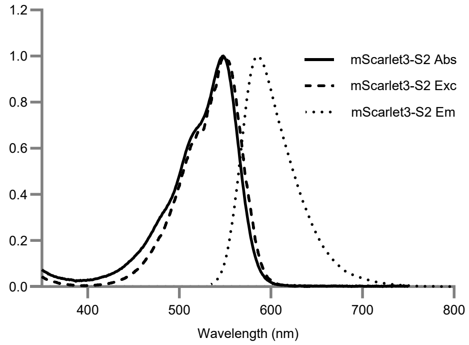

**SEQ ID NO: 77**

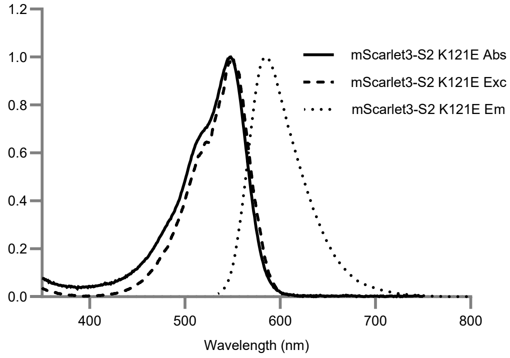

**SEQ ID NO: 79**

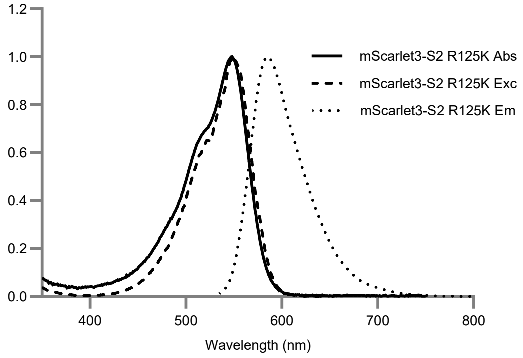

**SEQ ID NO: 81**

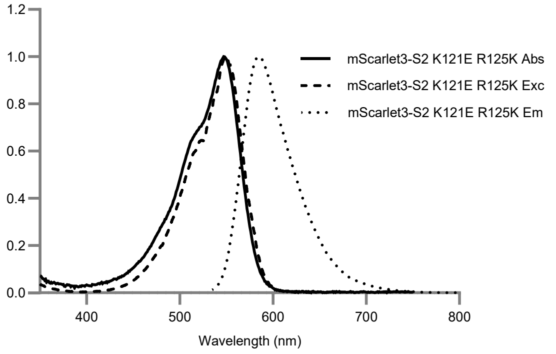

#### Table S3: mCranberry and variants thereof

The amino acid residue numbering for all mCherry and mCranberry series variants is initiated at position - 4 (Met), corresponding to the numbering convention established in PDB Accession No. 2H5Q (encompassed by SEQ ID NOs: 1–49). Conversely, amino acid residue numbering for the mScarlet series variants initiates at position 1 (Met), consistent with PDB Accession No. 5LK4 (encompassed by SEQ ID NOs: 51-57), whereas numbering for the mScarlet3 series variants initiates at residue position 1 (Met), consistent with PDB Accession No. 7ZCT (encompassed by SEQ ID NOs: 59–81). In the table below the amino acid substitution are with respect to following parent proteins: a-Seq ID 1-5, b- Seq ID 7-49, c- Seq ID 51-57, d-Seq ID 59-81.

| Mutants | AA Substitutions relative to mCherry[a], RCB2[b] mScarlet[c] and mScarlet3[d] | AA Sequence | DNA Sequence |
|---|---|---|---|
| mCherry | - | SEQ ID NO: 1 | SEQ ID NO: 2 |
| mCherry-XL R125K | R125K, W143S, I161V, Q163Y, I197R | SEQ ID NO: 3 | SEQ ID NO: 4 |
| RCB2 | K92R, Q114K, K121E, R125K, W143S, I161V, Q163Y, K166N, H172K, I197R | SEQ ID NO: 5 | SEQ ID NO: 6 |
| mCranberry | R92K, K114Q, N166K, K172H | SEQ ID NO: 7 | SEQ ID NO: 8 |
| mCranberry2 | E121K, N166K, K172H | SEQ ID NO: 9 | SEQ ID NO: 10 |
| K114Q K172H | K114Q, K172H | SEQ ID NO: 11 | SEQ ID NO: 12 |
| R92K K172H | R92K, K172H | SEQ ID NO: 13 | SEQ ID NO: 14 |
| E121K K172H | E121K, K172H | SEQ ID NO: 15 | SEQ ID NO: 16 |
| N166K K172H | N166K, K172H | SEQ ID NO: 17 | SEQ ID NO: 18 |
| R92K K114Q V135L K172H | R92K, K114Q, V135L, K172H | SEQ ID NO: 19 | SEQ ID NO: 20 |
| K114Q E121K K172H | K114Q, E121K, K172H | SEQ ID NO: 21 | SEQ ID NO: 22 |
| R92K N166K K172H | R92K, N166K, K172H | SEQ ID NO: 23 | SEQ ID NO: 24 |
| R92K E121K K172H | R92K, E121K, K172H | SEQ ID NO: 25 | SEQ ID NO: 26 |
| K114Q N166K K172H | K114Q, N166K, K172H | SEQ ID NO: 27 | SEQ ID NO: 28 |

| | | | |
|---|---|---|---|
| R92K K114Q E121K K172H | R92K, K114Q, E121K, K172H | SEQ ID NO: 29 | SEQ ID NO: 30 |
| R92K E121K N166K K172H | R92K, E121K, N166K, K172H | SEQ ID NO: 31 | SEQ ID NO: 32 |
| K114Q E121K N166K K172H | K114Q, E121K, N166K, K172H | SEQ ID NO: 33 | SEQ ID NO: 34 |
| R92K | R92K | SEQ ID NO: 35 | SEQ ID NO: 36 |
| K114Q | K114Q | SEQ ID NO: 37 | SEQ ID NO: 38 |
| E121K | E121K | SEQ ID NO: 39 | SEQ ID NO: 40 |
| K125R | K125R | SEQ ID NO: 41 | SEQ ID NO: 42 |
| N166K | N166K | SEQ ID NO: 43 | SEQ ID NO: 44 |
| K172H | K172H | SEQ ID NO: 45 | SEQ ID NO: 46 |
| R92K K114Q K125R N166K K172H | R92K, K114Q, K125R, N166K, K172H | SEQ ID NO: 47 | SEQ ID NO: 48 |
| R92K K114Q E121D N166K K172H | R92K, K114Q, E121D, N166K, K172H | SEQ ID NO: 49 | SEQ ID NO: 50 |
| mScarlet | - | SEQ ID NO: 51 | SEQ ID NO: 52 |
| mScarlet K122E | K122E | SEQ ID NO: 53 | SEQ ID NO: 54 |
| mScarlet R126K | R126K | SEQ ID NO: 55 | SEQ ID NO: 56 |
| mScarlet K122E R126K | K122E R126K | SEQ ID NO: 57 | SEQ ID NO: 58 |
| mScarlet3 | - | SEQ ID NO: 59 | SEQ ID NO: 60 |
| mScarlet3 K121E | K121E | SEQ ID NO: 61 | SEQ ID NO: 62 |
| mScarlet3 R125K | R125K | SEQ ID NO: 63 | SEQ ID NO: 64 |
| mScarlet3 K121E R125K | K121E R125K | SEQ ID NO: 65 | SEQ ID NO: 66 |
| mScarlet3-H | M164H | SEQ ID NO: 67 | SEQ ID NO: 68 |
| mScarlet3-H K121E | K121E, M164H | SEQ ID NO: 69 | SEQ ID NO: 70 |
| mScarlet3-H R125K | R125K, M164H | SEQ ID NO: 71 | SEQ ID NO: 72 |

| | | | |
|---|---|---|---|
| mScarlet3-H K121E R125K | K121E, R125K, M164H | SEQ ID NO: 73 | SEQ ID NO: 74 |
| mScarlet3-S2 | M67Q, M164H | SEQ ID NO: 75 | SEQ ID NO: 76 |
| mScarlet3-S2 K121E | M67Q, K121E, M164H | SEQ ID NO: 77 | SEQ ID NO: 78 |
| mScarlet3-S2 R125K | M67Q, R125K, M164H | SEQ ID NO: 79 | SEQ ID NO: 80 |
| mScarlet3-S2 K121E R125K | M67Q, K121E, R125K, M164H | SEQ ID NO: 81 | SEQ ID NO: 82 |

#### Table S4: Protein Sequence IDs

| Seq ID NO | Sequence |
|---|---|
| SEQ ID NO: 1 | MVSKGEEDNMAIIKEFMRFKVHMEGSVNGHEFEIEGEGEGRPYEGTQTAKLKVTK<br>GGPLPFAWDILSPQFMYGSKAYVKHPADIPDYLKLSFPEGFKWERVMNFEDGGVV<br>TVTQDSSLQDGEFIYKVKLRGTNFPSDGPVMQKKTMGWEASSERMYPEDGALKG<br>EIKQRLKLKDGGHYDAEVKTTYKAKKPVQLPGAYNVNIKLDITSHNEDYTIVEQYER<br>AEGRHSTGGMDELYK |
| SEQ ID NO: 3 | MVSKGEEDNMAIIKEFMRFKVHMEGSVNGHEFEIEGEGEGRPYEGTQTAKLKVTK<br>GGPLPFAWDILSPQFMYGSKAYVKHPADIPDYLKLSFPEGFKWERVMNFEDGGVV<br>TVTQDSSLQDGEFIYKVKLKGTNFPSDGPVMQKKTMGSEASSERMYPEDGALKG<br>EVKYRLKLKDGGHYDAEVKTTYKAKKPVQLPGAYNVNRKLDITSHNEDYTIVEQYE<br>RAEGRHSTGGMDELYK |
| SEQ ID NO: 5 | MVSKGEEDNMAIIKEFMRFKVHMEGSVNGHEFEIEGEGEGRPYEGTQTAKLKVTK<br>GGPLPFAWDILSPQFMYGSKAYVKHPADIPDYLKLSFPEGFRWERVMNFEDGGVV<br>TVTQDSSLKDGEFIYEVKLKGTNFPSDGPVMQKKTMGSEASSERMYPEDGALKGE<br>VKYRLNLKDGGKYDAEVKTTYKAKKPVQLPGAYNVNRKLDITSHNEDYTIVEQYER<br>AEGRHSTGGMDELYK |
| SEQ ID NO: 7 | MVSKGEEDNMAIIKEFMRFKVHMEGSVNGHEFEIEGEGEGRPYEGTQTAKLKVTK<br>GGPLPFAWDILSPQFMYGSKAYVKHPADIPDYLKLSFPEGFKWERVMNFEDGGVV<br>TVTQDSSLQDGEFIYEVKLKGTNFPSDGPVMQKKTMGSEASSERMYPEDGALKG<br>EVKYRLKLKDGGHYDAEVKTTYKAKKPVQLPGAYNVNRKLDITSHNEDYTIVEQYE<br>RAEGRHSTGGMDELYK |
| SEQ ID NO: 9 | MVSKGEEDNMAIIKEFMRFKVHMEGSVNGHEFEIEGEGEGRPYEGTQTAKLKVTK<br>GGPLPFAWDILSPQFMYGSKAYVKHPADIPDYLKLSFPEGFRWERVMNFEDGGVV<br>TVTQDSSLKDGEFIYKVKLKGTNFPSDGPVMQKKTMGSEASSERMYPEDGALKGE<br>VKYRLKLKDGGHYDAEVKTTYKAKKPVQLPGAYNVNRKLDITSHNEDYTIVEQYER<br>AEGRHSTGGMDELYK |
| SEQ ID NO: 11 | MVSKGEEDNMAIIKEFMRFKVHMEGSVNGHEFEIEGEGEGRPYEGTQTAKLKVTK<br>GGPLPFAWDILSPQFMYGSKAYVKHPADIPDYLKLSFPEGFRWERVMNFEDGGVV<br>TVTQDSSLQDGEFIYEVKLKGTNFPSDGPVMQKKTMGSEASSERMYPEDGALKG<br>EVKYRLNLKDGGHYDAEVKTTYKAKKPVQLPGAYNVNRKLDITSHNEDYTIVEQYE<br>RAEGRHSTGGMDELYK |
| SEQ ID NO: 13 | MVSKGEEDNMAIIKEFMRFKVHMEGSVNGHEFEIEGEGEGRPYEGTQTAKLKVTK<br>GGPLPFAWDILSPQFMYGSKAYVKHPADIPDYLKLSFPEGFKWERVMNFEDGGVV<br>TVTQDSSLKDGEFIYEVKLKGTNFPSDGPVMQKKTMGSEASSERMYPEDGALKGE<br>VKYRLNLKDGGHYDAEVKTTYKAKKPVQLPGAYNVNRKLDITSHNEDYTIVEQYER<br>AEGRHSTGGMDELYK |
| SEQ ID NO: 15 | MVSKGEEDNMAIIKEFMRFKVHMEGSVNGHEFEIEGEGEGRPYEGTQTAKLKVTK<br>GGPLPFAWDILSPQFMYGSKAYVKHPADIPDYLKLSFPEGFRWERVMNFEDGGVV<br>TVTQDSSLKDGEFIYKVKLKGTNFPSDGPVMQKKTMGSEASSERMYPEDGALKGE<br>VKYRLNLKDGGHYDAEVKTTYKAKKPVQLPGAYNVNRKLDITSHNEDYTIVEQYER<br>AEGRHSTGGMDELYK |
| SEQ ID NO: 17 | MVSKGEEDNMAIIKEFMRFKVHMEGSVNGHEFEIEGEGEGRPYEGTQTAKLKVTK<br>GGPLPFAWDILSPQFMYGSKAYVKHPADIPDYLKLSFPEGFRWERVMNFEDGGVV<br>TVTQDSSLKDGEFIYEVKLKGTNFPSDGPVMQKKTMGSEASSERMYPEDGALKGE<br>VKYRLKLKDGGHYDAEVKTTYKAKKPVQLPGAYNVNRKLDITSHNEDYTIVEQYER<br>AEGRHSTGGMDELYK |
| SEQ ID NO: 19 | MVSKGEEDNMAIIKEFMRFKVHMEGSVNGHEFEIEGEGEGRPYEGTQTAKLKVTK<br>GGPLPFAWDILSPQFMYGSKAYVKHPADIPDYLKLSFPEGFKWERVMNFEDGGVV<br>TVTQDSSLQDGEFIYEVKLKGTNFPSDGPLMQKKTMGSEASSERMYPEDGALKGE<br>VKYRLNLKDGGHYDAEVKTTYKAKKPVQLPGAYNVNRKLDITSHNEDYTIVEQYER<br>AEGRHSTGGMDELYK |

| | |
|---|---|
| SEQ ID NO: 21 | MVSKGEEDNMAIIKEFMRFKVHMEGSVNGHEFEIEGEGEGRPYEGTQTAKLKVTKGGPLPFAWDILSPQFMYGSKAYVKHPADIPDYLKLSFPEGFRWERVMNFEDGGVVTVTQDSSLQDGEFIYKVKLKGTNFPSDGPVMQKKTMGSEASSERMYPEDGALKGEVKYRLNLKDGGHYDAEVKTTYKAKKPVQLPGAYNVNRKLDITSHNEDYTIVEQYERAEGRHSTGGMDELYK |
| SEQ ID NO: 23 | MVSKGEEDNMAIIKEFMRFKVHMEGSVNGHEFEIEGEGEGRPYEGTQTAKLKVTKGGPLPFAWDILSPQFMYGSKAYVKHPADIPDYLKLSFPEGFKWERVMNFEDGGVVTVTQDSSLKDGEFIYEVKLKGTNFPSDGPVMQKKTMGSEASSERMYPEDGALKGEVKYRLKLKDGGHYDAEVKTTYKAKKPVQLPGAYNVNRKLDITSHNEDYTIVEQYERAEGRHSTGGMDELYK |
| SEQ ID NO: 25 | MVSKGEEDNMAIIKEFMRFKVHMEGSVNGHEFEIEGEGEGRPYEGTQTAKLKVTKGGPLPFAWDILSPQFMYGSKAYVKHPADIPDYLKLSFPEGFKWERVMNFEDGGVVTVTQDSSLKDGEFIYKVKLKGTNFPSDGPVMQKKTMGSEASSERMYPEDGALKGEVKYRLNLKDGGHYDAEVKTTYKAKKPVQLPGAYNVNRKLDITSHNEDYTIVEQYERAEGRHSTGGMDELYK |
| SEQ ID NO: 27 | MVSKGEEDNMAIIKEFMRFKVHMEGSVNGHEFEIEGEGEGRPYEGTQTAKLKVTKGGPLPFAWDILSPQFMYGSKAYVKHPADIPDYLKLSFPEGFRWERVMNFEDGGVVTVTQDSSLQDGEFIYEVKLKGTNFPSDGPVMQKKTMGSEASSERMYPEDGALKGEVKYRLKLKDGGHYDAEVKTTYKAKKPVQLPGAYNVNRKLDITSHNEDYTIVEQYERAEGRHSTGGMDELYK |
| SEQ ID NO: 29 | MVSKGEEDNMAIIKEFMRFKVHMEGSVNGHEFEIEGEGEGRPYEGTQTAKLKVTKGGPLPFAWDILSPQFMYGSKAYVKHPADIPDYLKLSFPEGFKWERVMNFEDGGVVTVTQDSSLQDGEFIYKVKLKGTNFPSDGPVMQKKTMGSEASSERMYPEDGALKGEVKYRLNLKDGGHYDAEVKTTYKAKKPVQLPGAYNVNRKLDITSHNEDYTIVEQYERAEGRHSTGGMDELYK |
| SEQ ID NO: 31 | MVSKGEEDNMAIIKEFMRFKVHMEGSVNGHEFEIEGEGEGRPYEGTQTAKLKVTKGGPLPFAWDILSPQFMYGSKAYVKHPADIPDYLKLSFPEGFKWERVMNFEDGGVVTVTQDSSLKDGEFIYKVKLKGTNFPSDGPVMQKKTMGSEASSERMYPEDGALKGEVKYRLKLKDGGHYDAEVKTTYKAKKPVQLPGAYNVNRKLDITSHNEDYTIVEQYERAEGRHSTGGMDELYK |
| SEQ ID NO: 33 | MVSKGEEDNMAIIKEFMRFKVHMEGSVNGHEFEIEGEGEGRPYEGTQTAKLKVTKGGPLPFAWDILSPQFMYGSKAYVKHPADIPDYLKLSFPEGFRWERVMNFEDGGVVTVTQDSSLQDGEFIYKVKLKGTNFPSDGPVMQKKTMGSEASSERMYPEDGALKGEVKYRLKLKDGGHYDAEVKTTYKAKKPVQLPGAYNVNRKLDITSHNEDYTIVEQYERAEGRHSTGGMDELYK |
| SEQ ID NO: 35 | MVSKGEEDNMAIIKEFMRFKVHMEGSVNGHEFEIEGEGEGRPYEGTQTAKLKVTKGGPLPFAWDILSPQFMYGSKAYVKHPADIPDYLKLSFPEGFKWERVMNFEDGGVVTVTQDSSLKDGEFIYEVKLKGTNFPSDGPVMQKKTMGSEASSERMYPEDGALKGEVKYRLNLKDGGKYDAEVKTTYKAKKPVQLPGAYNVNRKLDITSHNEDYTIVEQYERAEGRHSTGGMDELYK |
| SEQ ID NO: 37 | MVSKGEEDNMAIIKEFMRFKVHMEGSVNGHEFEIEGEGEGRPYEGTQTAKLKVTKGGPLPFAWDILSPQFMYGSKAYVKHPADIPDYLKLSFPEGFRWERVMNFEDGGVVTVTQDSSLQDGEFIYEVKLKGTNFPSDGPVMQKKTMGSEASSERMYPEDGALKGEVKYRLNLKDGGKYDAEVKTTYKAKKPVQLPGAYNVNRKLDITSHNEDYTIVEQYERAEGRHSTGGMDELYK |
| SEQ ID NO: 39 | MVSKGEEDNMAIIKEFMRFKVHMEGSVNGHEFEIEGEGEGRPYEGTQTAKLKVTKGGPLPFAWDILSPQFMYGSKAYVKHPADIPDYLKLSFPEGFRWERVMNFEDGGVVTVTQDSSLKDGEFIYKVKLKGTNFPSDGPVMQKKTMGSEASSERMYPEDGALKGEVKYRLNLKDGGKYDAEVKTTYKAKKPVQLPGAYNVNRKLDITSHNEDYTIVEQYERAEGRHSTGGMDELYK |
| SEQ ID NO: 41 | MVSKGEEDNMAIIKEFMRFKVHMEGSVNGHEFEIEGEGEGRPYEGTQTAKLKVTKGGPLPFAWDILSPQFMYGSKAYVKHPADIPDYLKLSFPEGFRWERVMNFEDGGVVTVTQDSSLKDGEFIYEVKLRGTNFPSDGPVMQKKTMGSEASSERMYPEDGALKGEVKYRLNLKDGGKYDAEVKTTYKAKKPVQLPGAYNVNRKLDITSHNEDYTIVEQYERAEGRHSTGGMDELYK |

| | |
|---|---|
| SEQ ID NO: 43 | MVSKGEEDNMAIIKEFMRFKVHMEGSVNGHEFEIEGEGEGRPYEGTQTAKLKVTKGGPLPFAWDILSPQFMYGSKAYVKHPADIPDYLKLSFPEGFRWERVMNFEDGGVVTVTQDSSLKDGEFIYEVKLKGTNFPSDGPVMQKKTMGSEASSERMYPEDGALKGEVKYRLKLKDGGKYDAEVKTTYKAKKPVQLPGAYNVNRKLDITSHNEDYTIVEQYERAEGRHSTGGMDELYK |
| SEQ ID NO: 45 | MVSKGEEDNMAIIKEFMRFKVHMEGSVNGHEFEIEGEGEGRPYEGTQTAKLKVTKGGPLPFAWDILSPQFMYGSKAYVKHPADIPDYLKLSFPEGFRWERVMNFEDGGVVTVTQDSSLKDGEFIYEVKLKGTNFPSDGPVMQKKTMGSEASSERMYPEDGALKGEVKYRLNLKDGGHYDAEVKTTYKAKKPVQLPGAYNVNRKLDITSHNEDYTIVEQYERAEGRHSTGGMDELYK |
| SEQ ID NO: 47 | MVSKGEEDNMAIIKEFMRFKVHMEGSVNGHEFEIEGEGEGRPYEGTQTAKLKVTKGGPLPFAWDILSPQFMYGSKAYVKHPADIPDYLKLSFPEGFKWERVMNFEDGGVVTVTQDSSLQDGEFIYEVKLRGTNFPSDGPVMQKKTMGSEASSERMYPEDGALKGEVKYRLKLKDGGHYDAEVKTTYKAKKPVQLPGAYNVNRKLDITSHNEDYTIVEQYERAEGRHSTGGMDELYK |
| SEQ ID NO: 49 | MVSKGEEDNMAIIKEFMRFKVHMEGSVNGHEFEIEGEGEGRPYEGTQTAKLKVTKGGPLPFAWDILSPQFMYGSKAYVKHPADIPDYLKLSFPEGFKWERVMNFEDGGVVTVTQDSSLQDGEFIYDVKLKGTNFPSDGPVMQKKTMGSEASSERMYPEDGALKGEVKYRLKLKDGGHYDAEVKTTYKAKKPVQLPGAYNVNRKLDITSHNEDYTIVEQYERAEGRHSTGGMDELYK |
| SEQ ID NO: 51 | MVSKGEAVIKEFMRFKVHMEGSMNGHEFEIEGEGEGRPYEGTQTAKLKVTKGGPLPFSWDILSPQFMYGSRAFTKHPADIPDYYKQSFPEGFKWERVMNFEDGGAVTVTQDTSLEDGTLIYKVKLRGTNFPPDGPVMQKKTMGWEASTERLYPEDGVLKGDIKMALRLKDGGRYLADFKTTYKAKKPVQMPGAYNVDRKLDITSHNEDYTVVEQYERSEGRHSTGGMDELYK |
| SEQ ID NO: 53 | MVSKGEAVIKEFMRFKVHMEGSMNGHEFEIEGEGEGRPYEGTQTAKLKVTKGGPLPFSWDILSPQFMYGSRAFTKHPADIPDYYKQSFPEGFKWERVMNFEDGGAVTVTQDTSLEDGTLIYEVKLRGTNFPPDGPVMQKKTMGWEASTERLYPEDGVLKGDIKMALRLKDGGRYLADFKTTYKAKKPVQMPGAYNVDRKLDITSHNEDYTVVEQYERSEGRHSTGGMDELYK |
| SEQ ID NO: 55 | MVSKGEAVIKEFMRFKVHMEGSMNGHEFEIEGEGEGRPYEGTQTAKLKVTKGGPLPFSWDILSPQFMYGSRAFTKHPADIPDYYKQSFPEGFKWERVMNFEDGGAVTVTQDTSLEDGTLIYKVKLKGTNFPPDGPVMQKKTMGWEASTERLYPEDGVLKGDIKMALRLKDGGRYLADFKTTYKAKKPVQMPGAYNVDRKLDITSHNEDYTVVEQYERSEGRHSTGGMDELYK |
| SEQ ID NO: 57 | MVSKGEAVIKEFMRFKVHMEGSMNGHEFEIEGEGEGRPYEGTQTAKLKVTKGGPLPFSWDILSPQFMYGSRAFTKHPADIPDYYKQSFPEGFKWERVMNFEDGGAVTVTQDTSLEDGTLIYEVKLKGTNFPPDGPVMQKKTMGWEASTERLYPEDGVLKGDIKMALRLKDGGRYLADFKTTYKAKKPVQMPGAYNVDRKLDITSHNEDYTVVEQYERSEGRHSTGGMDELYK |
| SEQ ID NO: 59 | MDSTEAVIKEFMRFKVHMEGSMNGHEFEIEGEGEGRPYEGTQTAKLRVTKGGPLPFSWDILSPQFMYGSRAFTKHPADIPDYWKQSFPEGFKWERVMNFEDGGAVSVAQDTSLEDGTLIYKVKLRGTNFPPDGPVMQKKTMGWEASTERLYPEDVVLKGDIKMALRLKDGGRYLADFKTTYRAKKPVQMPGAFNIDRKLDITSHNEDYTVVEQYERSVARHSTGGSGGS |
| SEQ ID NO: 61 | MVSKGEAVIKEFMRFKVHMEGSMNGHEFEIEGEGEGRPYEGTQTAKLRVTKGGPLPFSWDILSPQFMYGSRAFTKHPADIPDYWKQSFPEGFKWERVMNFEDGGAVSVAQDTSLEDGTLIYEVKLRGTNFPPDGPVMQKKTMGWEASTERLYPEDVVLKGDIKMALRLKDGGRYLADFKTTYRAKKPVQMPGAFNIDRKLDITSHNEDYTVVEQYERSVARHSTGGMDELYK |
| SEQ ID NO: 63 | MDSTEAVIKEFMRFKVHMEGSMNGHEFEIEGEGEGRPYEGTQTAKLRVTKGGPLPFSWDILSPQFMYGSRAFTKHPADIPDYWKQSFPEGFKWERVMNFEDGGAVSVAQDTSLEDGTLIYKVKLKGTNFPPDGPVMQKKTMGWEASTERLYPEDVVLKGDIKMALRLKDGGRYLADFKTTYRAKKPVQMPGAFNIDRKLDITSHNEDYTVVEQYERSVARHSTGGSGGS |

| | |
|---|---|
| SEQ ID NO: 65 | MVSKGEAVIKEFMRFKVHMEGSMNGHEFEIEGEGEGRPYEGTQTAKLRVTKGGPL PFSWDILSPQFMYGSRAFTKHPADIPDYWKQSFPEGFKWERVMNFEDGGAVSVA QDTSLEDGTLIYEVKLKGTNFPPDGPVMQKKTMGWEASTERLYPEDVVLKGDIKM ALRLKDGGRYLADFKTTYRAKKPVQMPGAFNIDRKLDITSHNEDYTVVEQYERSVA RHSTGGMDELYK |
| SEQ ID NO: 67 | MVSKGEAVIKEFMRFKVHMEGSMNGHEFEIEGEGEGRPYEGTQTAKLRVTKGGPL PFSWDILSPQFMYGSRAFTKHPADIPDYWKQSFPEGFKWERVMNFEDGGAVSVA QDTSLEDGTLIYKVKLRGTNFPPDGPVMQKKTMGWEASTERLYPEDVVLKGDIKH ALRLKDGGRYLADFKTTYRAKKPVQMPGAFNIDRKLDITSHNEDYTVVEQYERSVA RHSTGGMDELYK |
| SEQ ID NO: 69 | MVSKGEAVIKEFMRFKVHMEGSMNGHEFEIEGEGEGRPYEGTQTAKLRVTKGGPL PFSWDILSPQFMYGSRAFTKHPADIPDYWKQSFPEGFKWERVMNFEDGGAVSVA QDTSLEDGTLIYEVKLRGTNFPPDGPVMQKKTMGWEASTERLYPEDVVLKGDIKH ALRLKDGGRYLADFKTTYRAKKPVQMPGAFNIDRKLDITSHNEDYTVVEQYERSVA RHSTGGMDELYK |
| SEQ ID NO: 71 | MVSKGEAVIKEFMRFKVHMEGSMNGHEFEIEGEGEGRPYEGTQTAKLRVTKGGPL PFSWDILSPQFMYGSRAFTKHPADIPDYWKQSFPEGFKWERVMNFEDGGAVSVA QDTSLEDGTLIYKVKLKGTNFPPDGPVMQKKTMGWEASTERLYPEDVVLKGDIKH ALRLKDGGRYLADFKTTYRAKKPVQMPGAFNIDRKLDITSHNEDYTVVEQYERSVA RHSTGGMDELYK |
| SEQ ID NO: 73 | MVSKGEAVIKEFMRFKVHMEGSMNGHEFEIEGEGEGRPYEGTQTAKLRVTKGGPL PFSWDILSPQFMYGSRAFTKHPADIPDYWKQSFPEGFKWERVMNFEDGGAVSVA QDTSLEDGTLIYEVKLKGTNFPPDGPVMQKKTMGWEASTERLYPEDVVLKGDIKH ALRLKDGGRYLADFKTTYRAKKPVQMPGAFNIDRKLDITSHNEDYTVVEQYERSVA RHSTGGMDELYK |
| SEQ ID NO: 75 | MVSKGEAVIKEFMRFKVHMEGSMNGHEFEIEGEGEGRPYEGTQTAKLRVTKGGPL PFSWDILSPQFQYGSRAFTKHPADIPDYWKQSFPEGFKWERVMNFEDGGAVSVA QDTSLEDGTLIYKVKLRGTNFPPDGPVMQKKTMGWEASTERLYPEDVVLKGDIKH ALRLKDGGRYLADFKTTYRAKKPVQMPGAFNIDRKLDITSHNEDYTVVEQYERSVA RHSTGGMDELYK |
| SEQ ID NO: 77 | MVSKGEAVIKEFMRFKVHMEGSMNGHEFEIEGEGEGRPYEGTQTAKLRVTKGGPL PFSWDILSPQFQYGSRAFTKHPADIPDYWKQSFPEGFKWERVMNFEDGGAVSVA QDTSLEDGTLIYEVKLRGTNFPPDGPVMQKKTMGWEASTERLYPEDVVLKGDIKH ALRLKDGGRYLADFKTTYRAKKPVQMPGAFNIDRKLDITSHNEDYTVVEQYERSVA RHSTGGMDELYK |
| SEQ ID NO: 79 | MVSKGEAVIKEFMRFKVHMEGSMNGHEFEIEGEGEGRPYEGTQTAKLRVTKGGPL PFSWDILSPQFQYGSRAFTKHPADIPDYWKQSFPEGFKWERVMNFEDGGAVSVA QDTSLEDGTLIYKVKLKGTNFPPDGPVMQKKTMGWEASTERLYPEDVVLKGDIKH ALRLKDGGRYLADFKTTYRAKKPVQMPGAFNIDRKLDITSHNEDYTVVEQYERSVA RHSTGGMDELYK |
| SEQ ID NO: 81 | MVSKGEAVIKEFMRFKVHMEGSMNGHEFEIEGEGEGRPYEGTQTAKLRVTKGGPL PFSWDILSPQFQYGSRAFTKHPADIPDYWKQSFPEGFKWERVMNFEDGGAVSVA QDTSLEDGTLIYEVKLKGTNFPPDGPVMQKKTMGWEASTERLYPEDVVLKGDIKH ALRLKDGGRYLADFKTTYRAKKPVQMPGAFNIDRKLDITSHNEDYTVVEQYERSVA RHSTGGMDELYK |

#### Table S5: DNA Sequence IDs

| Seq ID NO | Sequence |
|---|---|
| SEQ ID NO: 2 | ATGGTGAGCAAGGGCGAAGAGGATAACATGGCCATCATCAAGGAGTTCATGCG<br>CTTCAAGGTGCACATGGAGGGCTCCGTGAACGGCCACGAGTTCGAGATCGAG<br>GGCGAGGGCGAGGGCCGCCCCTACGAGGGCACCCAGACCGCCAAGCTGAAG<br>GTGACCAAGGGTGGCCCCCTGCCCTTCGCCTGGGACATCCTGTCCCCTCAGTT<br>CATGTACGGCTCCAAGGCCTACGTGAAGCACCCCGCCGACATCCCCGACTACT<br>TGAAGCTGTCCTTCCCCGAGGGCTTCAAGTGGGAGCGCGTGATGAACTTCGAG<br>GACGGCGGCGTGGTGACCGTGACCCAGGACTCCTCCCTGCAGGACGGCGAG<br>TTCATCTACAAGGTGAAGCTGCGCGGCACCAACTTCCCCTCCGACGGCCCCGT<br>AATGCAGAAGAAGACCATGGGCTGGGAGGCCTCCTCCGAGCGGATGTACCCC<br>GAGGACGGCGCCCTGAAGGGCGAGATCAAGCAGAGGCTGAAGCTGAAGGAC<br>GGCGGCCACTACGACGCTGAGGTCAAGACCACCTACAAGGCCAAGAAGCCCG<br>TGCAGCTGCCCGGCGCCTACAACGTCAACATCAAGTTGGACATCACCTCCCAC<br>AACGAGGACTACACCATCGTGGAACAGTACGAACGCGCCGAGGGCCGCCACT<br>CCACCGGCGGCATGGACGAGCTGTACAAG |
| SEQ ID NO: 4 | ATGGTGAGCAAGGGCGAGGAGGATAACATGGCCATCATCAAGGAGTTCATGCG<br>CTTCAAGGTGCACATGGAGGGCTCCGTGAACGGCCACGAGTTCGAGATCGAG<br>GGCGAGGGCGAGGGCCGCCCCTACGAGGGCACCCAGACCGCCAAGCTGAAG<br>GTGACCAAGGGTGGCCCCCTGCCCTTCGCCTGGGACATCCTGTCCCCTCAGTT<br>CATGTACGGCTCCAAGGCCTACGTGAAGCACCCCGCCGACATCCCCGACTACT<br>TGAAGCTGTCATTCCCCGAGGGCTTCAAGTGGGAGCGCGTGATGAACTTCGAG<br>GACGGCGGCGTGGTGACCGTGACCCAGGACTCCTCCCTGCAGGACGGCGAG<br>TTCATCTACAAGGTGAAGCTGAAGGGCACCAACTTTCCCTCCGACGGCCCCGT<br>AATGCAGAAGAAGACCATGGGCAGTGAGGCCTCCTCCGAGCGGATGTACCCCG<br>AGGACGGCGCCCTGAAGGGCGAGGTTAAGTATAGGCTGAAGCTGAAGGACGG<br>CGGCCACTACGACGCTGAGGTCAAGACCACCTACAAGGCCAAGAAGCCCGTG<br>CAGCTGCCCGGCGCCTACAACGTCAACCGTAAGTTGGACATCACCTCCCACAA<br>CGAGGACTACACCATCGTGGAACAGTACGAACGCGCCGAGGGCCGCCACTCC<br>ACCGGCGGCATGGACGAGCTGTACAAG |
| SEQ ID NO: 6 | ATGGTGAGCAAGGGCGAGGAGGATAACATGGCCATCATCAAGGAGTTCATGCG<br>CTTCAAGGTGCACATGGAGGGCTCCGTGAACGGCCACGAGTTCGAGATCGAG<br>GGCGAGGGCGAGGGCCGCCCCTACGAGGGCACCCAGACCGCCAAGCTGAAG<br>GTGACCAAGGGTGGCCCCCTGCCCTTCGCCTGGGACATCCTGTCCCCTCAGTT<br>CATGTACGGCTCCAAGGCCTACGTGAAGCACCCCGCCGACATCCCCGACTACT<br>TGAAGCTGTCATTCCCCGAGGGCTTCAGGTGGGAGCGCGTGATGAATTTCGAG<br>GACGGCGGCGTGGTGACCGTGACCCAGGACTCCTCCCTGAAGGACGGCGAG<br>TTCATCTACGAAGTGAAGCTGAAGGGCACCAACTTTCCCTCCGACGGCCCCGT<br>AATGCAGAAGAAGACCATGGGCAGTGAGGCCTCCTCCGAGCGGATGTACCCCG<br>AGGACGGCGCCCTGAAGGGCGAGGTTAAATATAGGCTGAATCTGAAGGACGGC<br>GGCAAATACGACGCTGAGGTCAAGACCACCTACAAGGCCAAGAAGCCCGTGCA<br>GCTGCCCGGCGCCTACAACGTCAACCGTAAGTTGGACATCACCTCCCACAACG<br>AGGACTACACCATCGTGGAACAGTACGAACGCGCCGAGGGCCGCCACTCCAC<br>CGGCGGCATGGACGAGCTGTACAAG |
| SEQ ID NO: 8 | ATGGTGAGCAAGGGCGAGGAGGATAACATGGCCATCATCAAGGAGTTCATGCG<br>CTTCAAGGTGCACATGGAGGGCTCCGTGAACGGCCACGAGTTCGAGATCGAG<br>GGCGAGGGCGAGGGCCGCCCCTACGAGGGCACCCAGACCGCCAAGCTGAAG<br>GTGACCAAGGGTGGCCCCCTGCCCTTCGCCTGGGACATCCTGTCCCCTCAGTT<br>CATGTACGGCTCCAAGGCCTACGTGAAGCACCCCGCCGACATCCCCGACTACT<br>TGAAGCTGTCATTCCCCGAGGGCTTCAAGTGGGAGCGCGTGATGAATTTCGAG<br>GACGGCGGCGTGGTGACCGTGACCCAGGACTCCTCCCTGCAGGACGGCGAG<br>TTCATCTACGAAGTGAAGCTGAAGGGCACCAACTTTCCCTCCGACGGCCCCGT |

| | |
|---|---|
| | AATGCAGAAGAAGACCATGGGCAGTGAGGCCTCCTCCGAGCGGATGTACCCCG<br>AGGACGGCGCCCTGAAGGGCGAGGTTAAATATAGGCTGAAGCTGAAGGACGG<br>CGGCCACTACGACGCTGAGGTCAAGACCACCTACAAGGCCAAGAAGCCCGTG<br>CAGCTGCCCGGCGCCTACAACGTCAACCGTAAGTTGGACATCACCTCCCACAA<br>CGAGGACTACACCATCGTGGAACAGTACGAACGCGCCGAGGGCCGCCACTCC<br>ACCGGCGGCATGGACGAGCTGTACAAG |
| SEQ ID NO: 10 | ATGGTGAGCAAGGGCGAGGAGGATAACATGGCCATCATCAAGGAGTTCATGCG<br>CTTCAAGGTGCACATGGAGGGCTCCGTGAACGGCCACGAGTTCGAGATCGAG<br>GGCGAGGGCGAGGGCCGCCCCTACGAGGGCACCCAGACCGCCAAGCTGAAG<br>GTGACCAAGGGTGGCCCCCTGCCCTTCGCCTGGGACATCCTGTCCCCTCAGTT<br>CATGTACGGCTCCAAGGCCTACGTGAAGCACCCCGCCGACATCCCCGACTACT<br>TGAAGCTGTCATTCCCCGAGGGCTTCAGGTGGGAGCGCGTGATGAATTTCGAG<br>GACGGCGGCGTGGTGACCGTGACCCAGGACTCCTCCCTGAAGGACGGCGAG<br>TTCATCTACAAAGTGAAGCTGAAGGGCACCAACTTTCCCTCCGACGGCCCCGT<br>AATGCAGAAGAAGACCATGGGCAGTGAGGCCTCCTCCGAGCGGATGTACCCCG<br>AGGACGGCGCCCTGAAGGGCGAGGTTAAATATAGGCTGAAGCTGAAGGACGG<br>CGGCCACTACGACGCTGAGGTCAAGACCACCTACAAGGCCAAGAAGCCCGTG<br>CAGCTGCCCGGCGCCTACAACGTCAACCGTAAGTTGGACATCACCTCCCACAA<br>CGAGGACTACACCATCGTGGAACAGTACGAACGCGCCGAGGGCCGCCACTCC<br>ACCGGCGGCATGGACGAGCTGTACAAG |
| SEQ ID NO: 12 | ATGGTGAGCAAGGGCGAGGAGGATAACATGGCCATCATCAAGGAGTTCATGCG<br>CTTCAAGGTGCACATGGAGGGCTCCGTGAACGGCCACGAGTTCGAGATCGAG<br>GGCGAGGGCGAGGGCCGCCCCTACGAGGGCACCCAGACCGCCAAGCTGAAG<br>GTGACCAAGGGTGGCCCCCTGCCCTTCGCCTGGGACATCCTGTCCCCTCAGTT<br>CATGTACGGCTCCAAGGCCTACGTGAAGCACCCCGCCGACATCCCCGACTACT<br>TGAAGCTGTCATTCCCCGAGGGCTTCAGGTGGGAGCGCGTGATGAATTTCGAG<br>GACGGCGGCGTGGTGACCGTGACCCAGGACTCCTCCCTGCAGGACGGCGAG<br>TTCATCTACGAAGTGAAGCTGAAGGGCACCAACTTTCCCTCCGACGGCCCCGT<br>AATGCAGAAGAAGACCATGGGCAGTGAGGCCTCCTCCGAGCGGATGTACCCCG<br>AGGACGGCGCCCTGAAGGGCGAGGTTAAATATAGGCTGAATCTGAAGGACGGC<br>GGCCACTACGACGCTGAGGTCAAGACCACCTACAAGGCCAAGAAGCCCGTGC<br>AGCTGCCCGGCGCCTACAACGTCAACCGTAAGTTGGACATCACCTCCCACAAC<br>GAGGACTACACCATCGTGGAACAGTACGAACGCGCCGAGGGCCGCCACTCCA<br>CCGGCGGCATGGACGAGCTGTACAAG |
| SEQ ID NO: 14 | ATGGTGAGCAAGGGCGAGGAGGATAACATGGCCATCATCAAGGAGTTCATGCG<br>CTTCAAGGTGCACATGGAGGGCTCCGTGAACGGCCACGAGTTCGAGATCGAG<br>GGCGAGGGCGAGGGCCGCCCCTACGAGGGCACCCAGACCGCCAAGCTGAAG<br>GTGACCAAGGGTGGCCCCCTGCCCTTCGCCTGGGACATCCTGTCCCCTCAGTT<br>CATGTACGGCTCCAAGGCCTACGTGAAGCACCCCGCCGACATCCCCGACTACT<br>TGAAGCTGTCATTCCCCGAGGGCTTCAAGTGGGAGCGCGTGATGAATTTCGAG<br>GACGGCGGCGTGGTGACCGTGACCCAGGACTCCTCCCTGAAGGACGGCGAG<br>TTCATCTACGAAGTGAAGCTGAAGGGCACCAACTTTCCCTCCGACGGCCCCGT<br>AATGCAGAAGAAGACCATGGGCAGTGAGGCCTCCTCCGAGCGGATGTACCCCG<br>AGGACGGCGCCCTGAAGGGCGAGGTTAAATATAGGCTGAATCTGAAGGACGGC<br>GGCCACTACGACGCTGAGGTCAAGACCACCTACAAGGCCAAGAAGCCCGTGC<br>AGCTGCCCGGCGCCTACAACGTCAACCGTAAGTTGGACATCACCTCCCACAAC<br>GAGGACTACACCATCGTGGAACAGTACGAACGCGCCGAGGGCCGCCACTCCA<br>CCGGCGGCATGGACGAGCTGTACAAG |
| SEQ ID NO: 16 | ATGGTGAGCAAGGGCGAGGAGGATAACATGGCCATCATCAAGGAGTTCATGCG<br>CTTCAAGGTGCACATGGAGGGCTCCGTGAACGGCCACGAGTTCGAGATCGAG<br>GGCGAGGGCGAGGGCCGCCCCTACGAGGGCACCCAGACCGCCAAGCTGAAG<br>GTGACCAAGGGTGGCCCCCTGCCCTTCGCCTGGGACATCCTGTCCCCTCAGTT<br>CATGTACGGCTCCAAGGCCTACGTGAAGCACCCCGCCGACATCCCCGACTACT<br>TGAAGCTGTCATTCCCCGAGGGCTTCAGGTGGGAGCGCGTGATGAATTTCGAG<br>GACGGCGGCGTGGTGACCGTGACCCAGGACTCCTCCCTGCAGGACGGCGAG<br>TTCATCTACGAAGTGAAGCTGAAGGGCACCAACTTTCCCTCCGACGGCCCCGT |

| | |
|---|---|
| | AATGCAGAAGAAGACCATGGGCAGTGAGGCCTCCTCCGAGCGGATGTACCCCG<br>AGGACGGCGCCCTGAAGGGCGAGGTTAAATATAGGCTGAATCTGAAGGACGGC<br>GGCCACTACGACGCTGAGGTCAAGACCACCTACAAGGCCAAGAAGCCCGTGC<br>AGCTGCCCGGCGCCTACAACGTCAACCGTAAGTTGGACATCACCTCCCACAAC<br>GAGGACTACACCATCGTGGAACAGTACGAACGCGCCGAGGGCCGCCACTCCA<br>CCGGCGGCATGGACGAGCTGTACAAG |
| SEQ ID NO: 18 | ATGGTGAGCAAGGGCGAGGAGGATAACATGGCCATCATCAAGGAGTTCATGCG<br>CTTCAAGGTGCACATGGAGGGCTCCGTGAACGGCCACGAGTTCGAGATCGAG<br>GGCGAGGGCGAGGGCCGCCCCTACGAGGGCACCCAGACCGCCAAGCTGAAG<br>GTGACCAAGGGTGGCCCCCTGCCCTTCGCCTGGGACATCCTGTCCCCTCAGTT<br>CATGTACGGCTCCAAGGCCTACGTGAAGCACCCCGCCGACATCCCCGACTACT<br>TGAAGCTGTCATTCCCCGAGGGCTTCAGGTGGGAGCGCGTGATGAATTTCGAG<br>GACGGCGGCGTGGTGACCGTGACCCAGGACTCCTCCCTGAAGGACGGCGAG<br>TTCATCTACGAAGTGAAGCTGAAGGGCACCAACTTTCCCTCCGACGGCCCCGT<br>AATGCAGAAGAAGACCATGGGCAGTGAGGCCTCCTCCGAGCGGATGTACCCCG<br>AGGACGGCGCCCTGAAGGGCGAGGTTAAATATAGGCTGAAGCTGAAGGACGG<br>CGGCCACTACGACGCTGAGGTCAAGACCACCTACAAGGCCAAGAAGCCCGTG<br>CAGCTGCCCGGCGCCTACAACGTCAACCGTAAGTTGGACATCACCTCCCACAA<br>CGAGGACTACACCATCGTGGAACAGTACGAACGCGCCGAGGGCCGCCACTCC<br>ACCGGCGGCATGGACGAGCTGTACAAG |
| SEQ ID NO: 20 | ATGGTGAGCAAGGGCGAGGAGGATAACATGGCCATCATCAAGGAGTTCATGCG<br>CTTCAAGGTGCACATGGAGGGCTCCGTGAACGGCCACGAGTTCGAGATCGAG<br>GGCGAGGGCGAGGGCCGCCCCTACGAGGGCACCCAGACCGCCAAGCTGAAG<br>GTGACCAAGGGTGGCCCCCTGCCCTTCGCCTGGGACATCCTGTCCCCTCAGTT<br>CATGTACGGCTCCAAGGCCTACGTGAAGCACCCCGCCGACATCCCCGACTACT<br>TGAAGCTGTCATTCCCCGAGGGCTTCAAGTGGGAGCGCGTGATGAATTTCGAG<br>GACGGCGGCGTGGTGACCGTGACCCAGGACTCCTCCCTGCAGGACGGCGAG<br>TTCATCTACGAAGTGAAGCTGAAGGGCACCAACTTTCCCTCCGACGGCCCCTTA<br>ATGCAGAAGAAGACCATGGGCAGTGAGGCCTCCTCCGAGCGGATGTACCCCGA<br>GGACGGCGCCCTGAAGGGCGAGGTTAAATATAGGCTGAATCTGAAGGACGGCG<br>GCCACTACGACGCTGAGGTCAAGACCACCTACAAGGCCAAGAAGCCCGTGCA<br>GCTGCCCGGCGCCTACAACGTCAACCGTAAGTTGGACATCACCTCCCACAACG<br>AGGACTACACCATCGTGGAACAGTACGAACGCGCCGAGGGCCGCCACTCCAC<br>CGGCGGCATGGACGAGCTGTACAAG |
| SEQ ID NO: 22 | ATGGTGAGCAAGGGCGAGGAGGATAACATGGCCATCATCAAGGAGTTCATGCG<br>CTTCAAGGTGCACATGGAGGGCTCCGTGAACGGCCACGAGTTCGAGATCGAG<br>GGCGAGGGCGAGGGCCGCCCCTACGAGGGCACCCAGACCGCCAAGCTGAAG<br>GTGACCAAGGGTGGCCCCCTGCCCTTCGCCTGGGACATCCTGTCCCCTCAGTT<br>CATGTACGGCTCCAAGGCCTACGTGAAGCACCCCGCCGACATCCCCGACTACT<br>TGAAGCTGTCATTCCCCGAGGGCTTCAGGTGGGAGCGCGTGATGAATTTCGAG<br>GACGGCGGCGTGGTGACCGTGACCCAGGACTCCTCCCTGCAGGACGGCGAG<br>TTCATCTACAAAGTGAAGCTGAAGGGCACCAACTTTCCCTCCGACGGCCCCGT<br>AATGCAGAAGAAGACCATGGGCAGTGAGGCCTCCTCCGAGCGGATGTACCCCG<br>AGGACGGCGCCCTGAAGGGCGAGGTTAAATATAGGCTGAATCTGAAGGACGGC<br>GGCCACTACGACGCTGAGGTCAAGACCACCTACAAGGCCAAGAAGCCCGTGC<br>AGCTGCCCGGCGCCTACAACGTCAACCGTAAGTTGGACATCACCTCCCACAAC<br>GAGGACTACACCATCGTGGAACAGTACGAACGCGCCGAGGGCCGCCACTCCA<br>CCGGCGGCATGGACGAGCTGTACAAG |
| SEQ ID NO: 24 | ATGGTGAGCAAGGGCGAGGAGGATAACATGGCCATCATCAAGGAGTTCATGCG<br>CTTCAAGGTGCACATGGAGGGCTCCGTGAACGGCCACGAGTTCGAGATCGAG<br>GGCGAGGGCGAGGGCCGCCCCTACGAGGGCACCCAGACCGCCAAGCTGAAG<br>GTGACCAAGGGTGGCCCCCTGCCCTTCGCCTGGGACATCCTGTCCCCTCAGTT<br>CATGTACGGCTCCAAGGCCTACGTGAAGCACCCCGCCGACATCCCCGACTACT<br>TGAAGCTGTCATTCCCCGAGGGCTTCAAGTGGGAGCGCGTGATGAATTTCGAG<br>GACGGCGGCGTGGTGACCGTGACCCAGGACTCCTCCCTGAAGGACGGCGAG<br>TTCATCTACGAAGTGAAGCTGAAGGGCACCAACTTTCCCTCCGACGGCCCCGT |

| | |
|---|---|
| | AATGCAGAAGAAGACCATGGGCAGTGAGGCCTCCTCCGAGCGGATGTACCCCG AGGACGGCGCCCTGAAGGGCGAGGTTAAATATAGGCTGAAGCTGAAGGACGG CGGCCACTACGACGCTGAGGTCAAGACCACCTACAAGGCCAAGAAGCCCGTG CAGCTGCCCGGCGCCTACAACGTCAACCGTAAGTTGGACATCACCTCCCACAA CGAGGACTACACCATCGTGGAACAGTACGAACGCGCCGAGGGCCGCCACTCC ACCGGCGGCATGGACGAGCTGTACAAG |
| SEQ ID NO: 26 | ATGGTGAGCAAGGGCGAGGAGGATAACATGGCCATCATCAAGGAGTTCATGCG CTTCAAGGTGCACATGGAGGGCTCCGTGAACGGCCACGAGTTCGAGATCGAG GGCGAGGGCGAGGGCCGCCCCTACGAGGGCACCCAGACCGCCAAGCTGAAG GTGACCAAGGGTGGCCCCCTGCCCTTCGCCTGGGACATCCTGTCCCCTCAGTT CATGTACGGCTCCAAGGCCTACGTGAAGCACCCCGCCGACATCCCCGACTACT TGAAGCTGTCATTCCCCGAGGGCTTCAAGTGGGAGCGCGTGATGAATTTCGAG GACGGCGGCGTGGTGACCGTGACCCAGGACTCCTCCCTGAAGGACGGCGAG TTCATCTACAAAGTGAAGCTGAAGGGCACCAACTTTCCCTCCGACGGCCCCGT AATGCAGAAGAAGACCATGGGCAGTGAGGCCTCCTCCGAGCGGATGTACCCCG AGGACGGCGCCCTGAAGGGCGAGGTTAAATATAGGCTGAATCTGAAGGACGGC GGCCACTACGACGCTGAGGTCAAGACCACCTACAAGGCCAAGAAGCCCGTGC AGCTGCCCGGCGCCTACAACGTCAACCGTAAGTTGGACATCACCTCCCACAAC GAGGACTACACCATCGTGGAACAGTACGAACGCGCCGAGGGCCGCCACTCCA CCGGCGGCATGGACGAGCTGTACAAG |
| SEQ ID NO: 28 | ATGGTGAGCAAGGGCGAGGAGGATAACATGGCCATCATCAAGGAGTTCATGCG CTTCAAGGTGCACATGGAGGGCTCCGTGAACGGCCACGAGTTCGAGATCGAG GGCGAGGGCGAGGGCCGCCCCTACGAGGGCACCCAGACCGCCAAGCTGAAG GTGACCAAGGGTGGCCCCCTGCCCTTCGCCTGGGACATCCTGTCCCCTCAGTT CATGTACGGCTCCAAGGCCTACGTGAAGCACCCCGCCGACATCCCCGACTACT TGAAGCTGTCATTCCCCGAGGGCTTCAGGTGGGAGCGCGTGATGAATTTCGAG GACGGCGGCGTGGTGACCGTGACCCAGGACTCCTCCCTGCAGGACGGCGAG TTCATCTACGAAGTGAAGCTGAAGGGCACCAACTTTCCCTCCGACGGCCCCGT AATGCAGAAGAAGACCATGGGCAGTGAGGCCTCCTCCGAGCGGATGTACCCCG AGGACGGCGCCCTGAAGGGCGAGGTTAAATATAGGCTGAAGCTGAAGGACGG CGGCCACTACGACGCTGAGGTCAAGACCACCTACAAGGCCAAGAAGCCCGTG CAGCTGCCCGGCGCCTACAACGTCAACCGTAAGTTGGACATCACCTCCCACAA CGAGGACTACACCATCGTGGAACAGTACGAACGCGCCGAGGGCCGCCACTCC ACCGGCGGCATGGACGAGCTGTACAAG |
| SEQ ID NO: 30 | ATGGTGAGCAAGGGCGAGGAGGATAACATGGCCATCATCAAGGAGTTCATGCG CTTCAAGGTGCACATGGAGGGCTCCGTGAACGGCCACGAGTTCGAGATCGAG GGCGAGGGCGAGGGCCGCCCCTACGAGGGCACCCAGACCGCCAAGCTGAAG GTGACCAAGGGTGGCCCCCTGCCCTTCGCCTGGGACATCCTGTCCCCTCAGTT CATGTACGGCTCCAAGGCCTACGTGAAGCACCCCGCCGACATCCCCGACTACT TGAAGCTGTCATTCCCCGAGGGCTTCAAGTGGGAGCGCGTGATGAATTTCGAG GACGGCGGCGTGGTGACCGTGACCCAGGACTCCTCCCTGCAGGACGGCGAG TTCATCTACAAAGTGAAGCTGAAGGGCACCAACTTTCCCTCCGACGGCCCCGT AATGCAGAAGAAGACCATGGGCAGTGAGGCCTCCTCCGAGCGGATGTACCCCG AGGACGGCGCCCTGAAGGGCGAGGTTAAATATAGGCTGAATCTGAAGGACGGC GGCCACTACGACGCTGAGGTCAAGACCACCTACAAGGCCAAGAAGCCCGTGC AGCTGCCCGGCGCCTACAACGTCAACCGTAAGTTGGACATCACCTCCCACAAC GAGGACTACACCATCGTGGAACAGTACGAACGCGCCGAGGGCCGCCACTCCA CCGGCGGCATGGACGAGCTGTACAAG |
| SEQ ID NO: 32 | ATGGTGAGCAAGGGCGAGGAGGATAACATGGCCATCATCAAGGAGTTCATGCG CTTCAAGGTGCACATGGAGGGCTCCGTGAACGGCCACGAGTTCGAGATCGAG GGCGAGGGCGAGGGCCGCCCCTACGAGGGCACCCAGACCGCCAAGCTGAAG GTGACCAAGGGTGGCCCCCTGCCCTTCGCCTGGGACATCCTGTCCCCTCAGTT CATGTACGGCTCCAAGGCCTACGTGAAGCACCCCGCCGACATCCCCGACTACT TGAAGCTGTCATTCCCCGAGGGCTTCAAGTGGGAGCGCGTGATGAATTTCGAG GACGGCGGCGTGGTGACCGTGACCCAGGACTCCTCCCTGAAGGACGGCGAG TTCATCTACAAAGTGAAGCTGAAGGGCACCAACTTTCCCTCCGACGGCCCCGT |

| | |
|---|---|
| | AATGCAGAAGAAGACCATGGGCAGTGAGGCCTCCTCCGAGCGGATGTACCCCG<br>AGGACGGCGCCCTGAAGGGCGAGGTTAAATATAGGCTGAAGCTGAAGGACGG<br>CGGCCACTACGACGCTGAGGTCAAGACCACCTACAAGGCCAAGAAGCCCGTG<br>CAGCTGCCCGGCGCCTACAACGTCAACCGTAAGTTGGACATCACCTCCCACAA<br>CGAGGACTACACCATCGTGGAACAGTACGAACGCGCCGAGGGCCGCCACTCC<br>ACCGGCGGCATGGACGAGCTGTACAAG |
| SEQ ID NO: 34 | ATGGTGAGCAAGGGCGAGGAGGATAACATGGCCATCATCAAGGAGTTCATGCG<br>CTTCAAGGTGCACATGGAGGGCTCCGTGAACGGCCACGAGTTCGAGATCGAG<br>GGCGAGGGCGAGGGCCGCCCCTACGAGGGCACCCAGACCGCCAAGCTGAAG<br>GTGACCAAGGGTGGCCCCCTGCCCTTCGCCTGGGACATCCTGTCCCCTCAGTT<br>CATGTACGGCTCCAAGGCCTACGTGAAGCACCCCGCCGACATCCCCGACTACT<br>TGAAGCTGTCATTCCCCGAGGGCTTCAGGTGGGAGCGCGTGATGAATTTCGAG<br>GACGGCGGCGTGGTGACCGTGACCCAGGACTCCTCCCTGCAGGACGGCGAG<br>TTCATCTACAAAGTGAAGCTGAAGGGCACCAACTTTCCCTCCGACGGCCCCGT<br>AATGCAGAAGAAGACCATGGGCAGTGAGGCCTCCTCCGAGCGGATGTACCCCG<br>AGGACGGCGCCCTGAAGGGCGAGGTTAAATATAGGCTGAAGCTGAAGGACGG<br>CGGCCACTACGACGCTGAGGTCAAGACCACCTACAAGGCCAAGAAGCCCGTG<br>CAGCTGCCCGGCGCCTACAACGTCAACCGTAAGTTGGACATCACCTCCCACAA<br>CGAGGACTACACCATCGTGGAACAGTACGAACGCGCCGAGGGCCGCCACTCC<br>ACCGGCGGCATGGACGAGCTGTACAAG |
| SEQ ID NO: 36 | ATGGTGAGCAAGGGCGAGGAGGATAACATGGCCATCATCAAGGAGTTCATGCG<br>CTTCAAGGTGCACATGGAGGGCTCCGTGAACGGCCACGAGTTCGAGATCGAG<br>GGCGAGGGCGAGGGCCGCCCCTACGAGGGCACCCAGACCGCCAAGCTGAAG<br>GTGACCAAGGGTGGCCCCCTGCCCTTCGCCTGGGACATCCTGTCCCCTCAGTT<br>CATGTACGGCTCCAAGGCCTACGTGAAGCACCCCGCCGACATCCCCGACTACT<br>TGAAGCTGTCATTCCCCGAGGGCTTCAAGTGGGAGCGCGTGATGAATTTCGAG<br>GACGGCGGCGTGGTGACCGTGACCCAGGACTCCTCCCTGAAGGACGGCGAG<br>TTCATCTACGAAGTGAAGCTGAAGGGCACCAACTTTCCCTCCGACGGCCCCGT<br>AATGCAGAAGAAGACCATGGGCAGTGAGGCCTCCTCCGAGCGGATGTACCCCG<br>AGGACGGCGCCCTGAAGGGCGAGGTTAAATATAGGCTGAATCTGAAGGACGGC<br>GGCAAATACGACGCTGAGGTCAAGACCACCTACAAGGCCAAGAAGCCCGTGCA<br>GCTGCCCGGCGCCTACAACGTCAACCGTAAGTTGGACATCACCTCCCACAACG<br>AGGACTACACCATCGTGGAACAGTACGAACGCGCCGAGGGCCGCCACTCCAC<br>CGGCGGCATGGACGAGCTGTACAAG |
| SEQ ID NO: 38 | ATGGTGAGCAAGGGCGAGGAGGATAACATGGCCATCATCAAGGAGTTCATGCG<br>CTTCAAGGTGCACATGGAGGGCTCCGTGAACGGCCACGAGTTCGAGATCGAG<br>GGCGAGGGCGAGGGCCGCCCCTACGAGGGCACCCAGACCGCCAAGCTGAAG<br>GTGACCAAGGGTGGCCCCCTGCCCTTCGCCTGGGACATCCTGTCCCCTCAGTT<br>CATGTACGGCTCCAAGGCCTACGTGAAGCACCCCGCCGACATCCCCGACTACT<br>TGAAGCTGTCATTCCCCGAGGGCTTCAGGTGGGAGCGCGTGATGAATTTCGAG<br>GACGGCGGCGTGGTGACCGTGACCCAGGACTCCTCCCTGCAGGACGGCGAG<br>TTCATCTACGAAGTGAAGCTGAAGGGCACCAACTTTCCCTCCGACGGCCCCGT<br>AATGCAGAAGAAGACCATGGGCAGTGAGGCCTCCTCCGAGCGGATGTACCCCG<br>AGGACGGCGCCCTGAAGGGCGAGGTTAAATATAGGCTGAATCTGAAGGACGGC<br>GGCAAATACGACGCTGAGGTCAAGACCACCTACAAGGCCAAGAAGCCCGTGCA<br>GCTGCCCGGCGCCTACAACGTCAACCGTAAGTTGGACATCACCTCCCACAACG<br>AGGACTACACCATCGTGGAACAGTACGAACGCGCCGAGGGCCGCCACTCCAC<br>CGGCGGCATGGACGAGCTGTACAAG |
| SEQ ID NO: 40 | ATGGTGAGCAAGGGCGAGGAGGATAACATGGCCATCATCAAGGAGTTCATGCG<br>CTTCAAGGTGCACATGGAGGGCTCCGTGAACGGCCACGAGTTCGAGATCGAG<br>GGCGAGGGCGAGGGCCGCCCCTACGAGGGCACCCAGACCGCCAAGCTGAAG<br>GTGACCAAGGGTGGCCCCCTGCCCTTCGCCTGGGACATCCTGTCCCCTCAGTT<br>CATGTACGGCTCCAAGGCCTACGTGAAGCACCCCGCCGACATCCCCGACTACT<br>TGAAGTTGTCATTCCCCGAGGGCTTCAGGTGGGAGCGCGTGATGAATTTCGAG<br>GACGGCGGCGTGGTGACCGTGACCCAGGACTCCTCCCTGAAGGACGGCGAG<br>TTCATCTACAAAGTGAAGCTGAAGGGCACCAACTTTCCCTCCGACGGCCCCGT |

| | |
|---|---|
| | AATGCAGAAGAAGACCATGGGCAGTGAGGCCTCCTCCGAGCGGATGTACCCCG AGGACGGCGCCCTGAAGGGCGAGGTTAAATATAGGCTGAATCTGAAGGACGGC GGCAAATACGACGCTGAGGTCAAGACCACCTACAAGGCCAAGAAGCCCGTGCA GCTGCCCGGCGCCTACAACGTCAACCGTAAGTTGGACATCACCTCCCACAACG AGGACTACACCATCGTGGAACAGTACGAACGCGCCGAGGGCCGCCACTCCAC CGGCGGCATGGACGAGCTGTACAAG |
| SEQ ID NO: 42 | ATGGTGAGCAAGGGCGAGGAGGATAACATGGCCATCATCAAGGAGTTCATGCG CTTCAAGGTGCACATGGAGGGCTCCGTGAACGGCCACGAGTTCGAGATCGAG GGCGAGGGCGAGGGCCGCCCCTACGAGGGCACCCAGACCGCCAAGCTGAAG GTGACCAAGGGTGGCCCCCTGCCCTTCGCCTGGGACATCCTGTCCCCTCAGTT CATGTACGGCTCCAAGGCCTACGTGAAGCACCCCGCCGACATCCCCGACTACT TGAAGCTGTCATTCCCCGAGGGCTTCAGGTGGGAGCGCGTGATGAATTTCGAG GACGGCGGCGTGGTGACCGTGACCCAGGACTCCTCCCTGAAGGACGGCGAG TTCATCTACGAAGTGAAGCTGAGGGGCACCAACTTTCCCTCCGACGGCCCCGT AATGCAGAAGAAGACCATGGGCAGTGAGGCCTCCTCCGAGCGGATGTACCCCG AGGACGGCGCCCTGAAGGGCGAGGTTAAATATAGGCTGAATCTGAAGGACGGC GGCAAATACGACGCTGAGGTCAAGACCACCTACAAGGCCAAGAAGCCCGTGCA GCTGCCCGGCGCCTACAACGTCAACCGTAAGTTGGACATCACCTCCCACAACG AGGACTACACCATCGTGGAACAGTACGAACGCGCCGAGGGCCGCCACTCCAC CGGCGGCATGGACGAGCTGTACAAG |
| SEQ ID NO: 44 | ATGGTGAGCAAGGGCGAGGAGGATAACATGGCCATCATCAAGGAGTTCATGCG CTTCAAGGTGCACATGGAGGGCTCCGTGAACGGCCACGAGTTCGAGATCGAG GGCGAGGGCGAGGGCCGCCCCTACGAGGGCACCCAGACCGCCAAGCTGAAG GTGACCAAGGGTGGCCCCCTGCCCTTCGCCTGGGACATCCTGTCCCCTCAGTT CATGTACGGCTCCAAGGCCTACGTGAAGCACCCCGCCGACATCCCCGACTACT TGAAGCTGTCATTCCCCGAGGGCTTCAGGTGGGAGCGCGTGATGAATTTCGAG GACGGCGGCGTGGTGACCGTGACCCAGGACTCCTCCCTGAAGGACGGCGAG TTCATCTACGAAGTGAAGCTGAAGGGCACCAACTTTCCCTCCGACGGCCCCGT AATGCAGAAGAAGACCATGGGCAGTGAGGCCTCCTCCGAGCGGATGTACCCCG AGGACGGCGCCCTGAAGGGCGAGGTTAAATATAGGCTGAAGCTGAAGGACGG CGGCAAATACGACGCTGAGGTCAAGACCACCTACAAGGCCAAGAAGCCCGTGC AGCTGCCCGGCGCCTACAACGTCAACCGTAAGTTGGACATCACCTCCCACAAC GAGGACTACACCATCGTGGAACAGTACGAACGCGCCGAGGGCCGCCACTCCA CCGGCGGCATGGACGAGCTGTACAAG |
| SEQ ID NO: 46 | ATGGTGAGCAAGGGCGAGGAGGATAACATGGCCATCATCAAGGAGTTCATGCG CTTCAAGGTGCACATGGAGGGCTCCGTGAACGGCCACGAGTTCGAGATCGAG GGCGAGGGCGAGGGCCGCCCCTACGAGGGCACCCAGACCGCCAAGCTGAAG GTGACCAAGGGTGGCCCCCTGCCCTTCGCCTGGGACATCCTGTCCCCTCAGTT CATGTACGGCTCCAAGGCCTACGTGAAGCACCCCGCCGACATCCCCGACTACT TGAAGCTGTCATTCCCCGAGGGCTTCAGGTGGGAGCGCGTGATGAATTTCGAG GACGGCGGCGTGGTGACCGTGACCCAGGACTCCTCCCTGAAGGACGGCGAG TTCATCTACGAAGTGAAGCTGAAGGGCACCAACTTTCCCTCCGACGGCCCCGT AATGCAGAAGAAGACCATGGGCAGTGAGGCCTCCTCCGAGCGGATGTACCCCG AGGACGGCGCCCTGAAGGGCGAGGTTAAATATAGGCTGAATCTGAAGGACGGC GGCCACTACGACGCTGAGGTCAAGACCACCTACAAGGCCAAGAAGCCCGTGC AGCTGCCCGGCGCCTACAACGTCAACCGTAAGTTGGACATCACCTCCCACAAC GAGGACTACACCATCGTGGAACAGTACGAACGCGCCGAGGGCCGCCACTCCA CCGGCGGCATGGACGAGCTGTACAAG |
| SEQ ID NO: 48 | ATGGTGAGCAAGGGCGAGGAGGATAACATGGCCATCATCAAGGAGTTCATGCG CTTCAAGGTGCACATGGAGGGCTCCGTGAACGGCCACGAGTTCGAGATCGAG GGCGAGGGCGAGGGCCGCCCCTACGAGGGCACCCAGACCGCCAAGCTGAAG GTGACCAAGGGTGGCCCCCTGCCCTTCGCCTGGGACATCCTGTCCCCTCAGTT CATGTACGGCTCCAAGGCCTACGTGAAGCACCCCGCCGACATCCCCGACTACT TGAAGCTGTCATTCCCCGAGGGCTTCAAGTGGGAGCGCGTGATGAATTTCGAG GACGGCGGCGTGGTGACCGTGACCCAGGACTCCTCCCTGCAGGACGGCGAG TTCATCTACGAAGTGAAGCTGCGCGGCACCAACTTTCCCTCCGACGGCCCCGT |

| | |
|---|---|
| | AATGCAGAAGAAGACCATGGGCAGTGAGGCCTCCTCCGAGCGGATGTACCCCG AGGACGGCGCCCTGAAGGGCGAGGTTAAATATAGGCTGAAGCTGAAGGACGG CGGCCACTACGACGCTGAGGTCAAGACCACCTACAAGGCCAAGAAGCCCGTG CAGCTGCCCGGCGCCTACAACGTCAACCGTAAGTTGGACATCACCTCCCACAA CGAGGACTACACCATCGTGGAACAGTACGAACGCGCCGAGGGCCGCCACTCC ACCGGCGGCATGGACGAGCTGTACAAG |
| SEQ ID NO: 50 | ATGGTGAGCAAGGGCGAGGAGGATAACATGGCCATCATCAAGGAGTTCATGCG CTTCAAGGTGCACATGGAGGGCTCCGTGAACGGCCACGAGTTCGAGATCGAG GGCGAGGGCGAGGGCCGCCCCTACGAGGGCACCCAGACCGCCAAGCTGAAG GTGACCAAGGGTGGCCCCCTGCCCTTCGCCTGGGACATCCTGTCCCCTCAGTT CATGTACGGCTCCAAGGCCTACGTGAAGCACCCCGCCGACATCCCCGACTACT TGAAGCTGTCATTCCCCGAGGGCTTCAAGTGGGAGCGCGTGATGAATTTCGAG GACGGCGGCGTGGTGACCGTGACCCAGGACTCCTCCCTGCAGGACGGCGAG TTCATCTACGATGTGAAGCTGAAGGGCACCAACTTTCCCTCCGACGGCCCCGTA ATGCAGAAGAAGACCATGGGCAGTGAGGCCTCCTCCGAGCGGATGTACCCCGA GGACGGCGCCCTGAAGGGCGAGGTTAAATATAGGCTGAAGCTGAAGGACGGC GGCCACTACGACGCTGAGGTCAAGACCACCTACAAGGCCAAGAAGCCCGTGC AGCTGCCCGGCGCCTACAACGTCAACCGTAAGTTGGACATCACCTCCCACAAC GAGGACTACACCATCGTGGAACAGTACGAACGCGCCGAGGGCCGCCACTCCA CCGGCGGCATGGACGAGCTGTACAAG |
| SEQ ID NO: 52 | ATGGTGAGCAAGGGCGAGGCAGTGATCAAGGAGTTCATGCGGTTCAAGGTGCA CATGGAGGGCTCCATGAACGGCCACGAGTTCGAGATCGAGGGCGAGGGCGAG GGCCGCCCCTACGAGGGCACCCAGACCGCCAAGCTGAAGGTGACCAAGGGTG GCCCCCTGCCCTTCTCCTGGGACATCCTGTCCCCTCAGTTCATGTACGGCTCC AGGGCCTTCACCAAGCACCCCGCCGACATCCCCGACTACTATAAGCAGTCCTT CCCCGAGGGCTTCAAGTGGGAGCGCGTGATGAACTTCGAGGACGGCGGCGCC GTGACCGTGACCCAGGACACCTCCCTGGAGGACGGCACCCTGATCTACAAGG TGAAGCTCCGCGGCACCAACTTCCCTCCTGACGGCCCCGTAATGCAGAAGAAG ACAATGGGCTGGGAAGCGTCCACCGAGCGGTTGTACCCCGAGGACGGCGTGC TGAAGGGCGACATTAAGATGGCCCTGCGCCTGAAGGACGGCGGCCGCTACCT GGCGGACTTCAAGACCACCTACAAGGCCAAGAAGCCCGTGCAGATGCCCGGC GCCTACAACGTCGACCGCAAGTTGGACATCACCTCCCACAACGAGGACTACAC CGTGGTGGAACAGTACGAACGCTCCGAGGGCCGCCACTCCACCGGCGGCATG GACGAGCTGTACAAG |
| SEQ ID NO: 54 | ATGGTGAGCAAGGGCGAGGCAGTGATCAAGGAGTTCATGCGGTTCAAGGTGCA CATGGAGGGCTCCATGAACGGCCACGAGTTCGAGATCGAGGGCGAGGGCGAG GGCCGCCCCTACGAGGGCACCCAGACCGCCAAGCTGAAGGTGACCAAGGGTG GCCCCCTGCCCTTCTCCTGGGACATCCTGTCCCCTCAGTTCATGTACGGCTCC AGGGCCTTCACCAAGCACCCCGCCGACATCCCCGACTACTATAAGCAGTCCTT CCCCGAGGGCTTCAAGTGGGAGCGCGTGATGAACTTCGAGGACGGCGGCGCC GTGACCGTGACCCAGGACACCTCCCTGGAGGACGGCACCCTGATCTACGAAG TGAAGCTCCGCGGCACCAACTTCCCTCCTGACGGCCCCGTAATGCAGAAGAAG ACAATGGGCTGGGAAGCGTCCACCGAGCGGTTGTACCCCGAGGACGGCGTGC TGAAGGGCGACATTAAGATGGCCCTGCGCCTGAAGGACGGCGGCCGCTACCT GGCGGACTTCAAGACCACCTACAAGGCCAAGAAGCCCGTGCAGATGCCCGGC GCCTACAACGTCGACCGCAAGTTGGACATCACCTCCCACAACGAGGACTACAC CGTGGTGGAACAGTACGAACGCTCCGAGGGCCGCCACTCCACCGGCGGCATG GACGAGCTGTACAAG |
| SEQ ID NO: 56 | ATGGTGAGCAAGGGCGAGGCAGTGATCAAGGAGTTCATGCGGTTCAAGGTGCA CATGGAGGGCTCCATGAACGGCCACGAGTTCGAGATCGAGGGCGAGGGCGAG GGCCGCCCCTACGAGGGCACCCAGACCGCCAAGCTGAAGGTGACCAAGGGTG GCCCCCTGCCCTTCTCCTGGGACATCCTGTCCCCTCAGTTCATGTACGGCTCC AGGGCCTTCACCAAGCACCCCGCCGACATCCCCGACTACTATAAGCAGTCCTT CCCCGAGGGCTTCAAGTGGGAGCGCGTGATGAACTTCGAGGACGGCGGCGCC GTGACCGTGACCCAGGACACCTCCCTGGAGGACGGCACCCTGATCTACAAGG TGAAGCTCAAAGGCACCAACTTCCCTCCTGACGGCCCCGTAATGCAGAAGAAG |

| | |
|---|---|
| | ACAATGGGCTGGGAAGCGTCCACCGAGCGGTTGTACCCCGAGGACGGCGTGC TGAAGGGCGACATTAAGATGGCCCTGCGCCTGAAGGACGGCGGCCGCTACCT GGCGGACTTCAAGACCACCTACAAGGCCAAGAAGCCCGTGCAGATGCCCGGC GCCTACAACGTCGACCGCAAGTTGGACATCACCTCCCACAACGAGGACTACAC CGTGGTGGAACAGTACGAACGCTCCGAGGGCCGCCACTCCACCGGCGGCATG GACGAGCTGTACAAG |
| SEQ ID NO: 58 | ATGGTGAGCAAGGGCGAGGCAGTGATCAAGGAGTTCATGCGGTTCAAGGTGCA CATGGAGGGCTCCATGAACGGCCACGAGTTCGAGATCGAGGGCGAGGGCGAG GGCCGCCCCTACGAGGGCACCCAGACCGCCAAGCTGAAGGTGACCAAGGGTG GCCCCCTGCCCTTCTCCTGGGACATCCTGTCCCCTCAGTTCATGTACGGCTCC AGGGCCTTCACCAAGCACCCCGCCGACATCCCCGACTACTATAAGCAGTCCTT CCCCGAGGGCTTCAAGTGGGAGCGCGTGATGAACTTCGAGGACGGCGGCGCC GTGACCGTGACCCAGGACACCTCCCTGGAGGACGGCACCCTGATCTACGAAG TGAAGCTCAAAGGCACCAACTTCCCTCCTGACGGCCCCGTAATGCAGAAGAAG ACAATGGGCTGGGAAGCGTCCACCGAGCGGTTGTACCCCGAGGACGGCGTGC TGAAGGGCGACATTAAGATGGCCCTGCGCCTGAAGGACGGCGGCCGCTACCT GGCGGACTTCAAGACCACCTACAAGGCCAAGAAGCCCGTGCAGATGCCCGGC GCCTACAACGTCGACCGCAAGTTGGACATCACCTCCCACAACGAGGACTACAC CGTGGTGGAACAGTACGAACGCTCCGAGGGCCGCCACTCCACCGGCGGCATG GACGAGCTGTACAAG |
| SEQ ID NO: 60 | ATGGATAGCACCGAGGCAGTGATCAAGGAGTTCATGCGGTTCAAGGTGCACAT GGAGGGCTCCATGAACGGCCACGAGTTCGAGATCGAGGGCGAGGGCGAGGG CCGCCCCTACGAGGGCACCCAGACCGCCAAGCTGAGGGTGACCAAGGGTGG CCCCCTGCCCTTCTCCTGGGACATCCTGTCCCCTCAGTTCATGTACGGCTCCA GGGCCTTCACGAAGCACCCCGCCGACATCCCCGACTACTGGAAGCAGTCCTTC CCCGAGGGCTTCAAGTGGGAGCGCGTGATGAACTTCGAGGACGGCGGCGCC GTGTCCGTGGCCCAGGACACCTCCCTGGAGGACGGCACCCTGATCTACAAGG TGAAGCTCCGCGGCACCAACTTCCCTCCTGACGGCCCCGTAATGCAGAAGAAG ACAATGGGCTGGGAAGCATCCACCGAGCGGTTGTACCCCGAGGACGTCGTGC TGAAGGGCGACATTAAGATGGCCCTGCGCCTGAAGGACGGCGGCCGCTACCT GGCGGACTTCAAGACCACCTACAGGGCCAAGAAGCCCGTGCAGATGCCCGGC GCCTTCAACATCGACCGCAAGTTGGACATCACATCCCACAACGAGGACTACACC GTGGTGGAACAGTACGAACGCTCCGTGGCCCGCCACTCCACCGGCGGCTCCG GTGGCTCC |
| SEQ ID NO: 62 | ATGGTGAGCAAGGGCGAGGCAGTGATCAAGGAGTTCATGCGGTTCAAGGTGCA CATGGAGGGCTCCATGAACGGCCACGAGTTCGAGATCGAGGGCGAGGGCGAG GGCCGCCCCTACGAGGGCACCCAGACCGCCAAGCTGAGGGTGACCAAGGGT GGCCCCCTGCCCTTCTCCTGGGACATCCTGTCCCCTCAGTTCATGTACGGCTC CAGGGCCTTCACGAAGCACCCCGCCGACATCCCCGACTACTGGAAGCAGTCCT TCCCCGAGGGCTTCAAGTGGGAGCGCGTGATGAACTTCGAGGACGGCGGCGC CGTGTCCGTGGCCCAGGACACCTCCCTGGAGGACGGCACCCTGATCTACGAA GTGAAGCTCCGCGGCACCAACTTCCCTCCTGACGGCCCCGTAATGCAGAAGAA GACAATGGGCTGGGAAGCATCCACCGAGCGGTTGTACCCCGAGGACGTCGTG CTGAAGGGCGACATTAAGATGGCCCTGCGCCTGAAGGACGGCGGCCGCTACC TGGCGGACTTCAAGACCACCTACAGGGCCAAGAAGCCCGTGCAGATGCCCGG CGCCTTCAACATCGACCGCAAGTTGGACATCACATCCCACAACGAGGACTACAC CGTGGTGGAACAGTACGAACGCTCCGTGGCCCGCCACTCCACCGGCGGCATG GACGAGCTGTACAAG |
| SEQ ID NO: 64 | ATGGATAGCACCGAGGCAGTGATCAAGGAGTTCATGCGGTTCAAGGTGCACAT GGAGGGCTCCATGAACGGCCACGAGTTCGAGATCGAGGGCGAGGGCGAGGG CCGCCCCTACGAGGGCACCCAGACCGCCAAGCTGAGGGTGACCAAGGGTGG CCCCCTGCCCTTCTCCTGGGACATCCTGTCCCCTCAGTTCATGTACGGCTCCA GGGCCTTCACGAAGCACCCCGCCGACATCCCCGACTACTGGAAGCAGTCCTTC CCCGAGGGCTTCAAGTGGGAGCGCGTGATGAACTTCGAGGACGGCGGCGCC GTGTCCGTGGCCCAGGACACCTCCCTGGAGGACGGCACCCTGATCTACAAGG TGAAGCTCAAAGGCACCAACTTCCCTCCTGACGGCCCCGTAATGCAGAAGAAG |

| | |
|---|---|
| | ACAATGGGCTGGGAAGCATCCACCGAGCGGTTGTACCCCGAGGACGTCGTGC<br>TGAAGGGCGACATTAAGATGGCCCTGCGCCTGAAGGACGGCGGCCGCTACCT<br>GGCGGACTTCAAGACCACCTACAGGGCCAAGAAGCCCGTGCAGATGCCCGGC<br>GCCTTCAACATCGACCGCAAGTTGGACATCACATCCCACAACGAGGACTACACC<br>GTGGTGGAACAGTACGAACGCTCCGTGGCCCGCCACTCCACCGGCGGCTCCG<br>GTGGCTCC |
| SEQ ID NO: 66 | ATGGTGAGCAAGGGCGAGGCAGTGATCAAGGAGTTCATGCGGTTCAAGGTGCA<br>CATGGAGGGCTCCATGAACGGCCACGAGTTCGAGATCGAGGGCGAGGGCGAG<br>GGCCGCCCCTACGAGGGCACCCAGACCGCCAAGCTGAGGGTGACCAAGGGT<br>GGCCCCCTGCCCTTCTCCTGGGACATCCTGTCCCCTCAGTTCATGTACGGCTC<br>CAGGGCCTTCACGAAGCACCCCGCCGACATCCCCGACTACTGGAAGCAGTCCT<br>TCCCCGAGGGCTTCAAGTGGGAGCGCGTGATGAACTTCGAGGACGGCGGCGC<br>CGTGTCCGTGGCCCAGGACACCTCCCTGGAGGACGGCACCCTGATCTACGAA<br>GTGAAGCTCAAAGGCACCAACTTCCCTCCTGACGGCCCCGTAATGCAGAAGAA<br>GACAATGGGCTGGGAAGCATCCACCGAGCGGTTGTACCCCGAGGACGTCGTG<br>CTGAAGGGCGACATTAAGATGGCCCTGCGCCTGAAGGACGGCGGCCGCTACC<br>TGGCGGACTTCAAGACCACCTACAGGGCCAAGAAGCCCGTGCAGATGCCCGG<br>CGCCTTCAACATCGACCGCAAGTTGGACATCACATCCCACAACGAGGACTACAC<br>CGTGGTGGAACAGTACGAACGCTCCGTGGCCCGCCACTCCACCGGCGGCATG<br>GACGAGCTGTACAAG |
| SEQ ID NO: 68 | ATGGTGAGCAAGGGCGAGGCAGTGATCAAGGAGTTCATGCGGTTCAAGGTGCA<br>CATGGAGGGCTCCATGAACGGCCACGAGTTCGAGATCGAGGGCGAGGGCGAG<br>GGCCGCCCCTACGAGGGCACCCAGACCGCCAAGCTGAGGGTGACCAAGGGT<br>GGCCCCCTGCCCTTCTCCTGGGACATCCTGTCCCCTCAGTTCATGTACGGCTC<br>CAGGGCCTTCACGAAGCACCCCGCCGACATCCCCGACTACTGGAAGCAGTCCT<br>TCCCCGAGGGCTTCAAGTGGGAGCGCGTGATGAACTTCGAGGACGGCGGCGC<br>CGTGTCCGTGGCCCAGGACACCTCCCTGGAGGACGGCACCCTGATCTACAAG<br>GTGAAGCTCCGCGGCACCAACTTCCCTCCTGACGGCCCCGTAATGCAGAAGAA<br>GACAATGGGCTGGGAAGCATCCACCGAGCGGTTGTACCCCGAGGACGTCGTG<br>CTGAAGGGCGACATTAAGCATGCCCTGCGCCTGAAGGACGGCGGCCGCTACC<br>TGGCGGACTTCAAGACCACCTACAGGGCCAAGAAGCCCGTGCAGATGCCCGG<br>CGCCTTCAACATCGACCGCAAGTTGGACATCACATCCCACAACGAGGACTACAC<br>CGTGGTGGAACAGTACGAACGCTCCGTGGCCCGCCACTCCACCGGCGGCATG<br>GACGAGCTGTACAAG |
| SEQ ID NO: 70 | ATGGTGAGCAAGGGCGAGGCAGTGATCAAGGAGTTCATGCGGTTCAAGGTGCA<br>CATGGAGGGCTCCATGAACGGCCACGAGTTCGAGATCGAGGGCGAGGGCGAG<br>GGCCGCCCCTACGAGGGCACCCAGACCGCCAAGCTGAGGGTGACCAAGGGT<br>GGCCCCCTGCCCTTCTCCTGGGACATCCTGTCCCCTCAGTTCATGTACGGCTC<br>CAGGGCCTTCACGAAGCACCCCGCCGACATCCCCGACTACTGGAAGCAGTCCT<br>TCCCCGAGGGCTTCAAGTGGGAGCGCGTGATGAACTTCGAGGACGGCGGCGC<br>CGTGTCCGTGGCCCAGGACACCTCCCTGGAGGACGGCACCCTGATCTACGAA<br>GTGAAGCTCCGCGGCACCAACTTCCCTCCTGACGGCCCCGTAATGCAGAAGAA<br>GACAATGGGCTGGGAAGCATCCACCGAGCGGTTGTACCCCGAGGACGTCGTG<br>CTGAAGGGCGACATTAAGCATGCCCTGCGCCTGAAGGACGGCGGCCGCTACC<br>TGGCGGACTTCAAGACCACCTACAGGGCCAAGAAGCCCGTGCAGATGCCCGG<br>CGCCTTCAACATCGACCGCAAGTTGGACATCACATCCCACAACGAGGACTACAC<br>CGTGGTGGAACAGTACGAACGCTCCGTGGCCCGCCACTCCACCGGCGGCATG<br>GACGAGCTGTACAAG |
| SEQ ID NO: 72 | ATGGTGAGCAAGGGCGAGGCAGTGATCAAGGAGTTCATGCGGTTCAAGGTGCA<br>CATGGAGGGCTCCATGAACGGCCACGAGTTCGAGATCGAGGGCGAGGGCGAG<br>GGCCGCCCCTACGAGGGCACCCAGACCGCCAAGCTGAGGGTGACCAAGGGT<br>GGCCCCCTGCCCTTCTCCTGGGACATCCTGTCCCCTCAGTTCATGTACGGCTC<br>CAGGGCCTTCACGAAGCACCCCGCCGACATCCCCGACTACTGGAAGCAGTCCT<br>TCCCCGAGGGCTTCAAGTGGGAGCGCGTGATGAACTTCGAGGACGGCGGCGC<br>CGTGTCCGTGGCCCAGGACACCTCCCTGGAGGACGGCACCCTGATCTACAAG<br>GTGAAGCTCAAAGGCACCAACTTCCCTCCTGACGGCCCCGTAATGCAGAAGAA |

| | |
|---|---|
| | GACAATGGGCTGGGAAGCATCCACCGAGCGGTTGTACCCCGAGGACGTCGTG CTGAAGGGCGACATTAAGCATGCCCTGCGCCTGAAGGACGGCGGCCGCTACC TGGCGGACTTCAAGACCACCTACAGGGCCAAGAAGCCCGTGCAGATGCCCGG CGCCTTCAACATCGACCGCAAGTTGGACATCACATCCCACAACGAGGACTACAC CGTGGTGGAACAGTACGAACGCTCCGTGGCCCGCCACTCCACCGGCGGCATG GACGAGCTGTACAAG |
| SEQ ID NO: 74 | ATGGTGAGCAAGGGCGAGGCAGTGATCAAGGAGTTCATGCGGTTCAAGGTGCA CATGGAGGGCTCCATGAACGGCCACGAGTTCGAGATCGAGGGCGAGGGCGAG GGCCGCCCCTACGAGGGCACCCAGACCGCCAAGCTGAGGGTGACCAAGGGT GGCCCCCTGCCCTTCTCCTGGGACATCCTGTCCCCTCAGTTCATGTACGGCTC CAGGGCCTTCACGAAGCACCCCGCCGACATCCCCGACTACTGGAAGCAGTCCT TCCCCGAGGGCTTCAAGTGGGAGCGCGTGATGAACTTCGAGGACGGCGGCGC CGTGTCCGTGGCCCAGGACACCTCCCTGGAGGACGGCACCCTGATCTACGAA GTGAAGCTCAAAGGCACCAACTTCCCTCCTGACGGCCCCGTAATGCAGAAGAA GACAATGGGCTGGGAAGCATCCACCGAGCGGTTGTACCCCGAGGACGTCGTG CTGAAGGGCGACATTAAGCATGCCCTGCGCCTGAAGGACGGCGGCCGCTACC TGGCGGACTTCAAGACCACCTACAGGGCCAAGAAGCCCGTGCAGATGCCCGG CGCCTTCAACATCGACCGCAAGTTGGACATCACATCCCACAACGAGGACTACAC CGTGGTGGAACAGTACGAACGCTCCGTGGCCCGCCACTCCACCGGCGGCATG GACGAGCTGTACAAG |
| SEQ ID NO: 76 | ATGGTGAGCAAGGGCGAGGCAGTGATCAAGGAGTTCATGCGGTTCAAGGTGCA CATGGAGGGCTCCATGAACGGCCACGAGTTCGAGATCGAGGGCGAGGGCGAG GGCCGCCCCTACGAGGGCACCCAGACCGCCAAGCTGAGGGTGACCAAGGGT GGCCCCCTGCCCTTCTCCTGGGACATCCTGTCCCCTCAGTTCCAGTACGGCTC CAGGGCCTTCACGAAGCACCCCGCCGACATCCCCGACTACTGGAAGCAGTCCT TCCCCGAGGGCTTCAAGTGGGAGCGCGTGATGAACTTCGAGGACGGCGGCGC CGTGTCCGTGGCCCAGGACACCTCCCTGGAGGACGGCACCCTGATCTACAAG GTGAAGCTCCGCGGCACCAACTTCCCTCCTGACGGCCCCGTAATGCAGAAGAA GACAATGGGCTGGGAAGCATCCACCGAGCGGTTGTACCCCGAGGACGTCGTG CTGAAGGGCGACATTAAGCATGCCCTGCGCCTGAAGGACGGCGGCCGCTACC TGGCGGACTTCAAGACCACCTACAGGGCCAAGAAGCCCGTGCAGATGCCCGG CGCCTTCAACATCGACCGCAAGTTGGACATCACATCCCACAACGAGGACTACAC CGTGGTGGAACAGTACGAACGCTCCGTGGCCCGCCACTCCACCGGCGGCATG GACGAGCTGTACAAG |
| SEQ ID NO: 78 | ATGGTGAGCAAGGGCGAGGCAGTGATCAAGGAGTTCATGCGGTTCAAGGTGCA CATGGAGGGCTCCATGAACGGCCACGAGTTCGAGATCGAGGGCGAGGGCGAG GGCCGCCCCTACGAGGGCACCCAGACCGCCAAGCTGAGGGTGACCAAGGGT GGCCCCCTGCCCTTCTCCTGGGACATCCTGTCCCCTCAGTTCCAGTACGGCTC CAGGGCCTTCACGAAGCACCCCGCCGACATCCCCGACTACTGGAAGCAGTCCT TCCCCGAGGGCTTCAAGTGGGAGCGCGTGATGAACTTCGAGGACGGCGGCGC CGTGTCCGTGGCCCAGGACACCTCCCTGGAGGACGGCACCCTGATCTACGAA GTGAAGCTCCGCGGCACCAACTTCCCTCCTGACGGCCCCGTAATGCAGAAGAA GACAATGGGCTGGGAAGCATCCACCGAGCGGTTGTACCCCGAGGACGTCGTG CTGAAGGGCGACATTAAGCATGCCCTGCGCCTGAAGGACGGCGGCCGCTACC TGGCGGACTTCAAGACCACCTACAGGGCCAAGAAGCCCGTGCAGATGCCCGG CGCCTTCAACATCGACCGCAAGTTGGACATCACATCCCACAACGAGGACTACAC CGTGGTGGAACAGTACGAACGCTCCGTGGCCCGCCACTCCACCGGCGGCATG GACGAGCTGTACAAG |
| SEQ ID NO: 80 | ATGGTGAGCAAGGGCGAGGCAGTGATCAAGGAGTTCATGCGGTTCAAGGTGCA CATGGAGGGCTCCATGAACGGCCACGAGTTCGAGATCGAGGGCGAGGGCGAG GGCCGCCCCTACGAGGGCACCCAGACCGCCAAGCTGAGGGTGACCAAGGGT GGCCCCCTGCCCTTCTCCTGGGACATCCTGTCCCCTCAGTTCCAGTACGGCTC CAGGGCCTTCACGAAGCACCCCGCCGACATCCCCGACTACTGGAAGCAGTCCT TCCCCGAGGGCTTCAAGTGGGAGCGCGTGATGAACTTCGAGGACGGCGGCGC CGTGTCCGTGGCCCAGGACACCTCCCTGGAGGACGGCACCCTGATCTACAAG GTGAAGCTCAAAGGCACCAACTTCCCTCCTGACGGCCCCGTAATGCAGAAGAA |

| | |
|---|---|
| | GACAATGGGCTGGGAAGCATCCACCGAGCGGTTGTACCCCGAGGACGTCGTG<br>CTGAAGGGCGACATTAAGCATGCCCTGCGCCTGAAGGACGGCGGCCGCTACC<br>TGGCGGACTTCAAGACCACCTACAGGGCCAAGAAGCCCGTGCAGATGCCCGG<br>CGCCTTCAACATCGACCGCAAGTTGGACATCACATCCCACAACGAGGACTACAC<br>CGTGGTGGAACAGTACGAACGCTCCGTGGCCCGCCACTCCACCGGCGGCATG<br>GACGAGCTGTACAAG |
| SEQ ID NO: 82 | ATGGTGAGCAAGGGCGAGGCAGTGATCAAGGAGTTCATGCGGTTCAAGGTGCA<br>CATGGAGGGCTCCATGAACGGCCACGAGTTCGAGATCGAGGGCGAGGGCGAG<br>GGCCGCCCCTACGAGGGCACCCAGACCGCCAAGCTGAGGGTGACCAAGGGT<br>GGCCCCCTGCCCTTCTCCTGGGACATCCTGTCCCCTCAGTTCCAGTACGGCTC<br>CAGGGCCTTCACGAAGCACCCCGCCGACATCCCCGACTACTGGAAGCAGTCCT<br>TCCCCGAGGGCTTCAAGTGGGAGCGCGTGATGAACTTCGAGGACGGCGGCGC<br>CGTGTCCGTGGCCCAGGACACCTCCCTGGAGGACGGCACCCTGATCTACGAA<br>GTGAAGCTCAAAGGCACCAACTTCCCTCCTGACGGCCCCGTAATGCAGAAGAA<br>GACAATGGGCTGGGAAGCATCCACCGAGCGGTTGTACCCCGAGGACGTCGTG<br>CTGAAGGGCGACATTAAGCATGCCCTGCGCCTGAAGGACGGCGGCCGCTACC<br>TGGCGGACTTCAAGACCACCTACAGGGCCAAGAAGCCCGTGCAGATGCCCGG<br>CGCCTTCAACATCGACCGCAAGTTGGACATCACATCCCACAACGAGGACTACAC<br>CGTGGTGGAACAGTACGAACGCTCCGTGGCCCGCCACTCCACCGGCGGCATG<br>GACGAGCTGTACAAG |

### Table S6: Crystallographic Data Summary

| Property | Value |
|---|---|
| Entry name | mCranberry2 (construct ID pending) |
| PDB Entry | 36RQ |
| Beamline | SSRL BL12-2 |
| Wavelength (Å) | 0.979 |
| Detector Distance (mm) | 150 mm |
| Space Group | P 21 21 21 |
| Unit Cell Dimensions | 83.14 Å, 137.34 Å, 357.18 Å / 90.00°, 90.00°, 90.00° |
| Resolution Range (Å) | 80.97 – 3.20 |
| Completeness (%) | 85.8 |
| Mean I/σ(I) | 1.65 (highest shell, 3.19 Å); |
| Wilson B-factor (Å²) | 73.1 |
| Rwork / Rfree | 0.201 / 0.267 |
| Reflections used in refinement | 55,890 (work + test) |
| Reflections used for Rfree | 2,997 (5.09%) |
| Number of non-H atoms | 27,588 |
| RMSD Bond Lengths (Å) | 0.010 |
| RMSD Bond Angles (°) | 1.361 |
| Ramachandran Favored (%) | 94.76 |
| Ramachandran Outliers (%) | 0.56 |
| Rotamer Outliers (%) | 0.00 |
| Clashscore | 7.71 |
| Average B, all atoms (Å²) | 82.0 |

#### Table S7: Hydrogen bond occupancies

Hydrogen bond occupancies for key residues and water molecules with the chromophore. Decimals are proportion of frames where at least one hydrogen bond is present between the chromophore and the listed residue, where values over 1 indicate more than 1 hydrogen bond is present some of the time.

| Protein | FQY | Ser69 | Ser146 | Waters |
|---|---|---|---|---|
| mCranberry | 85 | 0.48 | 0.39 | 1.3200 |
| mCranberry2 | 80 | 0.29 | 0.18 | 1.3000 |
| RCB2 | 77 | 0.47 | 0.21 | 1.3800 |
| K114Q K172H | 71 | 0.49 | 0.36 | 1.0900 |
| mCherry-XL | 70 | 0.56 | 0.17 | 1.3900 |
| mCherry | 21 | 0.39 | 0.04 | 1.3000 |

#### Table S8: Bridging hydrogen bond occupancies

Bridging hydrogen bond occupancies for key residues and water molecules with the chromophore. Approximate percentages are listed.

| Protein | FQY | Bridging residues | | | | | |
|---|---|---|---|---|---|---|---|
| mCranberry | 85 | Ser146 | 10.00% | Ser146 | 8.00% | Arg200 | 0.0500 |
| mCranberry2 | 80 | Ser146 | 14.00% | Glu147, Ser146 | 13.00% | | |
| RCB2 | 77 | Ser143 | 12.00% | Ser149 | 8.00% | Glu147 | 8.00% |
| K114Q K172H | 71 | Glu144 | 11.00% | Glu147, Ser146 | 10.00% | Ser149 | 5.00% |
| mCherry-XL | 70 | Glu144 | 18.00% | Glu147, Arg200 | 8.00% | | |
| mCherry | 21 | Glu144 | 10.00% | | | | |

#### Table S9: All chromophore hydrogen bond occupancies

All chromophore hydrogen bond occupancies for each system including differences as compared to the wild type.

| **mCherry** | **21** | | | | | | **Compare To K172H** |
|---|---|---|---|---|---|---|---|
| CH6_66@O2 | Arg | 95 | 21643 | 0.8657 | 2.828 | 162.3725 | |
| CH6_66@N1 | Ser | 69 | 9870 | 0.3948 | 2.9163 | 146.5474 | |
| CH6_66@O3 | Lys | 70 | 8254 | 0.3302 | 2.8618 | 153.2917 | |
| CH6_66@O3 | Trp | 93 | 5336 | 0.2134 | 2.9018 | 152.5817 | |
| CH6_66@OH | Gln | 163 | 3442 | 0.1377 | 2.8506 | 152.5763 | |
| CH6_66@OH | Gln | 163 | 2338 | 0.0935 | 2.8631 | 152.7628 | |
| **mCranberry** | **86** | | | | | | |
| CH6_66@O2 | Arg | 95 | 43510 | 0.8702 | 2.8194 | 162.2298 | 0.0045 |

| | | | | | | | |
|---|---|---|---|---|---|---|---|
| **CH6_66@O3** | **Lys** | **70** | **35790** | **0.7158** | **2.866** | **154.1203** | **0.3856** |
| CH6_66@N1 | Ser | 69 | 24034 | 0.4807 | 2.9261 | 151.4165 | 0.0859 |
| **CH6_66@OH** | **Ser** | **146** | **19444** | **0.3889** | **2.7518** | **162.5648** | **0.3889** |
| **CH6_66@O3** | **Trp** | **93** | **3065** | **0.0613** | **2.9121** | **147.119** | **-0.1521** |
| CH6_66@O2 | Tyr | 181 | 2383 | 0.0477 | 2.791 | 152.1814 | 0.0477 |
| **K114Q K172H** | **71** | | | | | | |
| CH6_66@O2 | Arg | 95 | 44951 | 0.899 | 2.8151 | 162.1707 | 0.0333 |
| CH6_66@N1 | Ser | 69 | 24577 | 0.4915 | 2.913 | 148.9426 | 0.0967 |
| **CH6_66@OH** | **Ser** | **146** | **17787** | **0.3557** | **2.7694** | **158.9263** | **0.3557** |
| CH6_66@O3 | Lys | 70 | 17716 | 0.3543 | 2.8579 | 154.3356 | 0.0241 |
| **CH6_66@O3** | **Trp** | **93** | **4659** | **0.0932** | **2.9082** | **147.7575** | **-0.1202** |
| | **Gln** | **163** | | | | | **-0.21** |
| **RCB2** | **77** | | | | | | |
| CH6_66@O2 | Arg | 95 | 39659 | 0.7932 | 2.8363 | 161.359 | -0.0725 |
| CH6_66@N1 | Ser | 69 | 23581 | 0.4716 | 2.9094 | 148.4747 | 0.0768 |
| CH6_66@O3 | Lys | 70 | 20091 | 0.4018 | 2.8698 | 150.9993 | 0.0716 |
| **CH6_66@OH** | **Ser** | **143** | **10587** | **0.2117** | **2.739** | **160.9261** | **0.2117** |
| CH6_66@O2 | Lys | 70 | 4776 | 0.0955 | 2.857 | 150.5331 | |
| **CH6_66@O3** | **Trp** | **93** | **4518** | **0.0904** | **2.9168** | **149.2552** | **-0.123** |
| CH6_66@O2 | Lys | 70 | 4333 | 0.0867 | 2.8564 | 150.5561 | |
| CH6_66@O2 | Lys | 70 | 4325 | 0.0865 | 2.8593 | 150.7965 | |
| **mCranberry2** | **80** | | | | | | |
| CH6_66@O2 | Arg | 95 | 44679 | 0.8952 | 2.8146 | 162.0508 | 0.0295 |
| CH6_66@N1 | Ser | 69 | 14416 | 0.2888 | 2.9227 | 147.7725 | -0.106 |
| **CH6_66@OH** | **Ser** | **143** | **8844** | **0.1772** | **2.7506** | **160.6044** | **0.1772** |
| **CH6_66@O3** | **Tyr** | **181** | **7814** | **0.1566** | **2.8261** | **158.9229** | **0.1566** |
| **CH6_66@O3** | **Lys** | **70** | **3907** | **0.0783** | **2.8593** | **153.1298** | **-0.2519** |
| **CH6_66@O3** | **Trp** | **93** | **3865** | **0.0774** | **2.9112** | **153.4828** | **-0.136** |
| CH6_66@OH | Ser | 146 | 2316 | 0.0464 | 2.7948 | 157.0467 | 0.0464 |
| **mCherr-XL** | **70** | | | | | | |
| CH6_66@O2 | **Arg** | 95 | 45906 | 0.9181 | 2.8136 | 162.2081 | 0.0524 |
| CH6_66@N1 | Ser | 69 | 28040 | 0.5608 | 2.9067 | 149.4813 | 0.166 |
| **CH6_66@OH** | **Ser** | **146** | **8547** | **0.1709** | **2.7814** | **157.5596** | **0.1709** |
| CH6_66@O3 | Lys | 70 | 5553 | 0.1111 | 2.8595 | 149.0028 | 0.0529 |
| CH6_66@O2 | Lys | 70 | 4893 | 0.0979 | 2.8311 | 151.5689 | |
| CH6_66@O2 | Lys | 70 | 4402 | 0.088 | 2.8286 | 151.5272 | |
| CH6_66@O2 | Lys | 70 | 4306 | 0.0861 | 2.8309 | 151.1222 | |
| **CH6_66@O3** | **Trp** | **93** | **2754** | **0.0551** | **2.9145** | **146.2233** | **-0.1583** |
| CH6_66@O2 | Arg | 95 | 2053 | 0.0411 | 2.9012 | 144.7673 | |
